# *KCND3* is a novel susceptibility locus for early repolarization

**DOI:** 10.1101/673640

**Authors:** Alexander Teumer, Teresa Trenkwalder, Thorsten Kessler, Yalda Jamshidi, Marten E. van den Berg, Bernhard Kaess, Christopher P. Nelson, Rachel Bastiaenen, Marzia De Bortoli, Alessandra Rossini, Isabel Deisenhofer, Klaus Stark, Solmaz Assa, Peter S. Braund, Claudia Cabrera, Anna F. Dominiczak, Martin Gögele, Leanne M. Hall, M. Arfan Ikram, Maryam Kavousi, Karl J. Lackner, Lifelines Cohort Study, Christian Müller, Thomas Münzel, Matthias Nauck, Sandosh Padmanabhan, Norbert Pfeiffer, Tim D. Spector, Andre G. Uitterlinden, Niek Verweij, Uwe Völker, Helen R. Warren, Mobeen Zafar, Stephan B. Felix, Jan A. Kors, Harold Snieder, Patricia B. Munroe, Cristian Pattaro, Christian Fuchsberger, Georg Schmidt, Ilja M. Nolte, Heribert Schunkert, Peter Pramstaller, Philipp S. Wild, Pim van der Harst, Bruno H. Stricker, Renate B. Schnabel, Nilesh J. Samani, Christian Hengstenberg, Marcus Dörr, Elijah R. Behr, Wibke Reinhard

## Abstract

The presence of an early repolarization pattern (ERP) on the surface electrocardiogram (ECG) is associated with risk of ventricular fibrillation and sudden cardiac death. Family studies have shown that ERP is a highly heritable trait but molecular genetic determinants are unknown. We assessed the ERP in 12-lead ECGs of 39,456 individuals and conducted a two-stage meta-analysis of genome-wide association studies (GWAS). In the discovery phase, we included 2,181 cases and 23,641 controls from eight European ancestry studies and identified 19 genome-wide significant (p<5E-8) variants in the *KCND3* (potassium voltage gated channel subfamily D member 3) gene with a p-value of 4.6E-10. Replication of two loci in four additional studies including 1,124 cases and 12,510 controls confirmed the association at the *KCND3* gene locus with a pooled odds ratio of 0.82, p=7.7E-12 (rs1545300 minor allele T). A subsequent GWAS meta-analysis combining all samples did not reveal additional loci. The lead SNP of the discovery stage (rs12090194) was in strong linkage disequilibrium with rs1545300 (r^2^=0.96, D’=1). Summary statistics based conditional analysis did not reveal any secondary signals. Co-localization analyses indicate causal effects of *KCND3* gene expression levels on ERP in both the left ventricle of the heart and in tibial artery.

In this study we identified for the first time a genome-wide significant association of a genetic variant with ERP. Our findings of a locus in the *KCND3* gene not only provide insights into the genetic determinants but also into the pathophysiological mechanism of ERP, revealing a promising candidate for functional studies.

## Introduction

The early repolarization pattern (ERP) is a common ECG finding characterized by an elevation at the QRS-ST junction (J-point) of at least 0.1 mV in two adjacent ECG leads. The prevalence of ERP in the general population ranges from 2 to 13% being more common in young athletic men(1–5). The classical notion of ERP being a benign ECG phenotype was challenged in 2008 by the landmark study of Haissaguerre and colleagues showing an association of ERP with increased risk of ventricular fibrillation and sudden cardiac death(6): the Early Repolarization Syndrome (ERS)(7). Since then several studies demonstrated an elevated risk of cardiovascular and all-cause mortality in individuals with ERP underscoring its arrhythmogenic potential(2, 8, 9). Although the mechanistic basis for malignant arrhythmias in ERS is unclear, it has been suggested that they occur as a result of an augmented transmural electrical dispersion of repolarization. Ex vivo studies point towards a central role of the cardiac transient outward potassium current (I_to_) in the development of both, ERP and ERS(10). Furthermore, case descriptions of ERS identified genetic variations in genes encoding proteins for cardiac ion channels(11–13). Studies among relatives of sudden arrhythmic death syndrome show that ERP is more prevalent in the relatives than in controls indicating that ERP is an important potentially inheritable pro-arrhythmic trait(14, 15). Moreover, in family studies the heritability estimate for the presence of ERP was h^2^=0.49 (16). However, estimates for common SNP heritability from unrelated individuals are lower(17). This may explain why the only available genome-wide association study (GWAS) on ERP failed to identify genetic variants reaching genome-wide significance(18), and indicates the need for GWAS with more power by including a larger number of ERP cases.

In order to identify genetic variations that convey susceptibility to ERP we performed a GWAS and meta-analysis in European ancestry individuals, using a combined two-stage GWAS approach with a discovery phase of 2,181 ERP cases and 23,641 controls from eight cohorts, and replicated the results in 1,124 cases and 12,510 controls from four additional cohorts.

## Results

Clinical characteristics of the study cohorts are depicted in Table 1. The proportion of ERP based on the definition by Haisaguerre and Macfarlane(6, 19) ranged from 6% to 14% which is in line with previously reported prevalence in the general population(2–4).

**Table 1:**
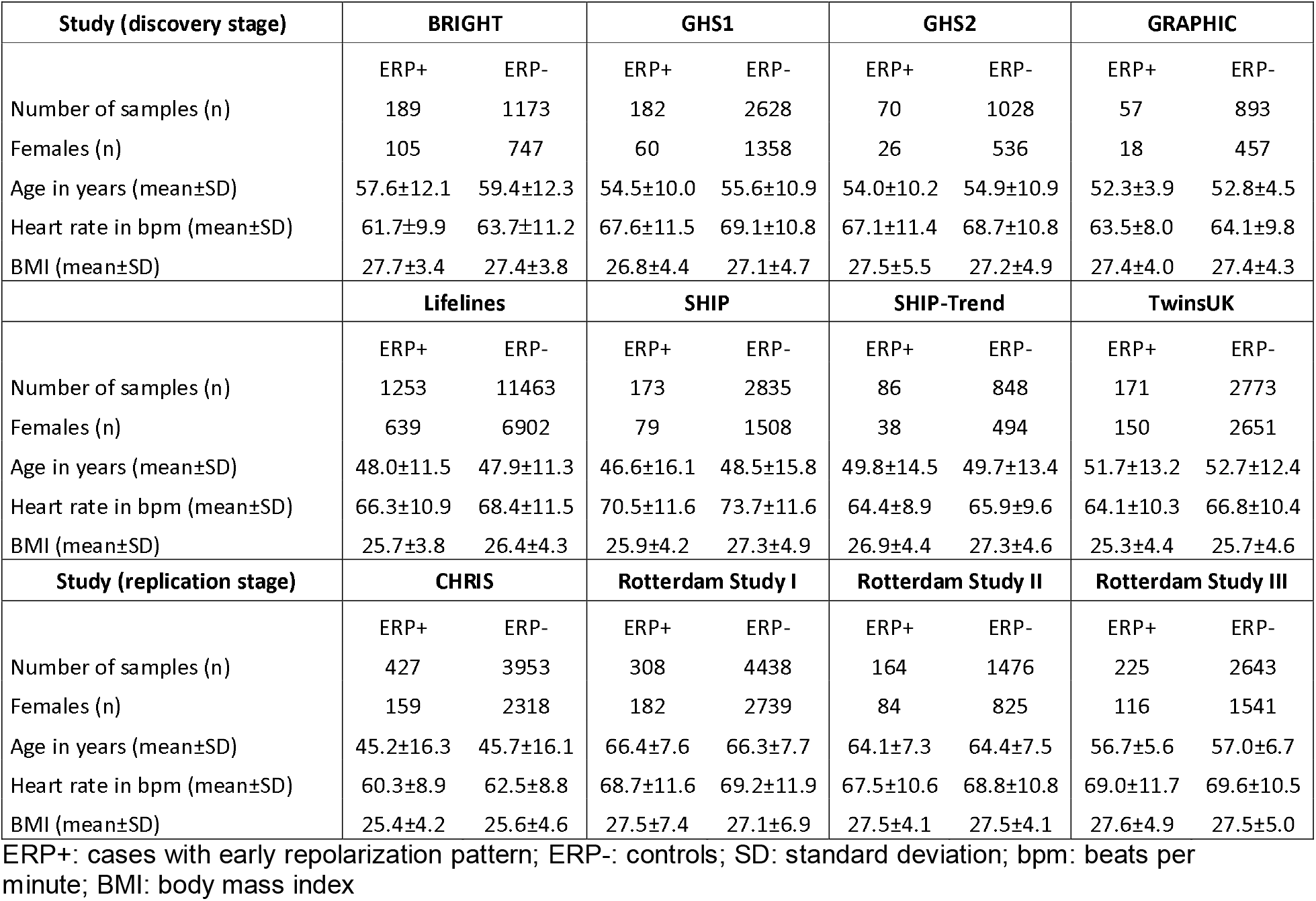
Baseline characteristics of the study populations

### Novel variants associated with ERP

We performed a GWAS meta-analysis in up to 2,181 cases and 23,641 controls from eight discovery cohorts. In total, 6,976,246 SNPs passed quality control (see Methods). We identified 19 variants spanning 49 kb in *KCND3* (Potassium Voltage-Gated Channel Subfamily D Member 3) as well as rs139772527 (effect allele frequency [EAF] 1.4%, OR=2.57, p=2.0E-8) near *HBZ* (Hemoglobin Subunit Zeta) to be genome-wide significantly associated (p<5E-8) with ERP The SNP with the lowest p-value of the region (lead SNP) at *KCND3* was the intronic rs12090194 (EAF 32.5%, OR=0.80, p=4.6E-10), and was replicated in an independent sample of 1,124 cases and 12,510 controls from four additional cohorts (p_replication_=2.5E-3, p_combined_=9.3E-12, Table 2). The SNP rs139772527 near *HBZ* did not fulfil the criteria for replication (p_replication_=0.28, p_combined_=1.4E-6, Table 2) as described in the Methods. The combined meta-analysis of all 12 cohorts including up to 39,456 individuals revealed only the locus at *KCND3* to be genome-wide significantly associated with ERP (**Supplementary Figure 1**). The lead SNP was rs1545300 (EAF 31.9%, OR=0.82, p=7.7E-12), followed by the discovery stage lead SNP rs12090194 being in strong linkage disequilibrium with rs1545300 (r²=0.96, D’=1) (Figure 1). Both SNPs were imputed at very high confidence (imputation quality score >0.97) in all cohorts. The quantile-quantile plots did not show any inflation (individual study λ_GC_ between 0.81 and 1.03, median: 0.91), and overall meta-analysis λ_GC_=1.02 (linkage disequilibrium [LD] score regression intercept: 1.01, see Methods) (**Supplementary Figure 2**). Summary statistics based conditional analysis to select independent hits did not reveal any secondary signals.

**Table 2:** Lead SNPs of the GWAS association results

| variant information |  |  |  | discovery |  |  |  |  | replication |  |  |  | combined |  |  |  |
| --- | --- | --- | --- | --- | --- | --- | --- | --- | --- | --- | --- | --- | --- | --- | --- | --- |
| SNP | chr:position | A1/A2 | nearest gene | AF1 | OR | P | I<sup>2</sup> | N | OR | P | I<sup>2</sup> | N | OR | P | I<sup>2</sup> | N |
| rs12090194 | 1:112,454,822 | t/c | KCND3 | 0.32 | 0.80 | **4.6E-10** | 34 | 25177 | 0.86 | 2.5E-03 | 39 | 13634 | 0.82 | 9.3E-12 | 35 | 38811 |
|  |  |  |  |  | [0.75-0.86] |  |  |  | [0.79-0.95] |  |  |  | [0.78-0.87] |  |  |  |
| rs1545300 | 1:112,464,004 | t/c | KCND3 | 0.32 | 0.81 | 1.4E-09 | 41 | 25172 | 0.85 | 9.4E-04 | 56 | 13634 | 0.82 | **7.7E-12** | 43 | 38806 |
|  |  |  |  |  | [0.75-0.86] |  |  |  | [0.77-0.94] |  |  |  | [0.78-0.87] |  |  |  |
| rs139772527 | 16:208,761 | t/c | HBZ | 0.01 | 2.57 | **2.0E-08** | 0 | 21495 | 1.21 | 2.8E-01 | 0 | 13634 | 1.81 | 1.4E-06 | 11 | 35129 |
|  |  |  |  |  | [1.85-3.58] |  |  |  | [0.85-1.73] |  |  |  | [1.42-2.31] |  |  |  |

A1: effect allele; AF1: allele frequency of A1; OR: odds ratio of A1 [95% confidence interval]; P: association p-value; I<sup>2</sup>: percentage of total variation across studies that is due to heterogeneity; N: sample size. Bold values indicate the lead SNP (lowest p-value) of a significantly associated locus in the corresponding meta-analysis stage.

**Figure 1.**
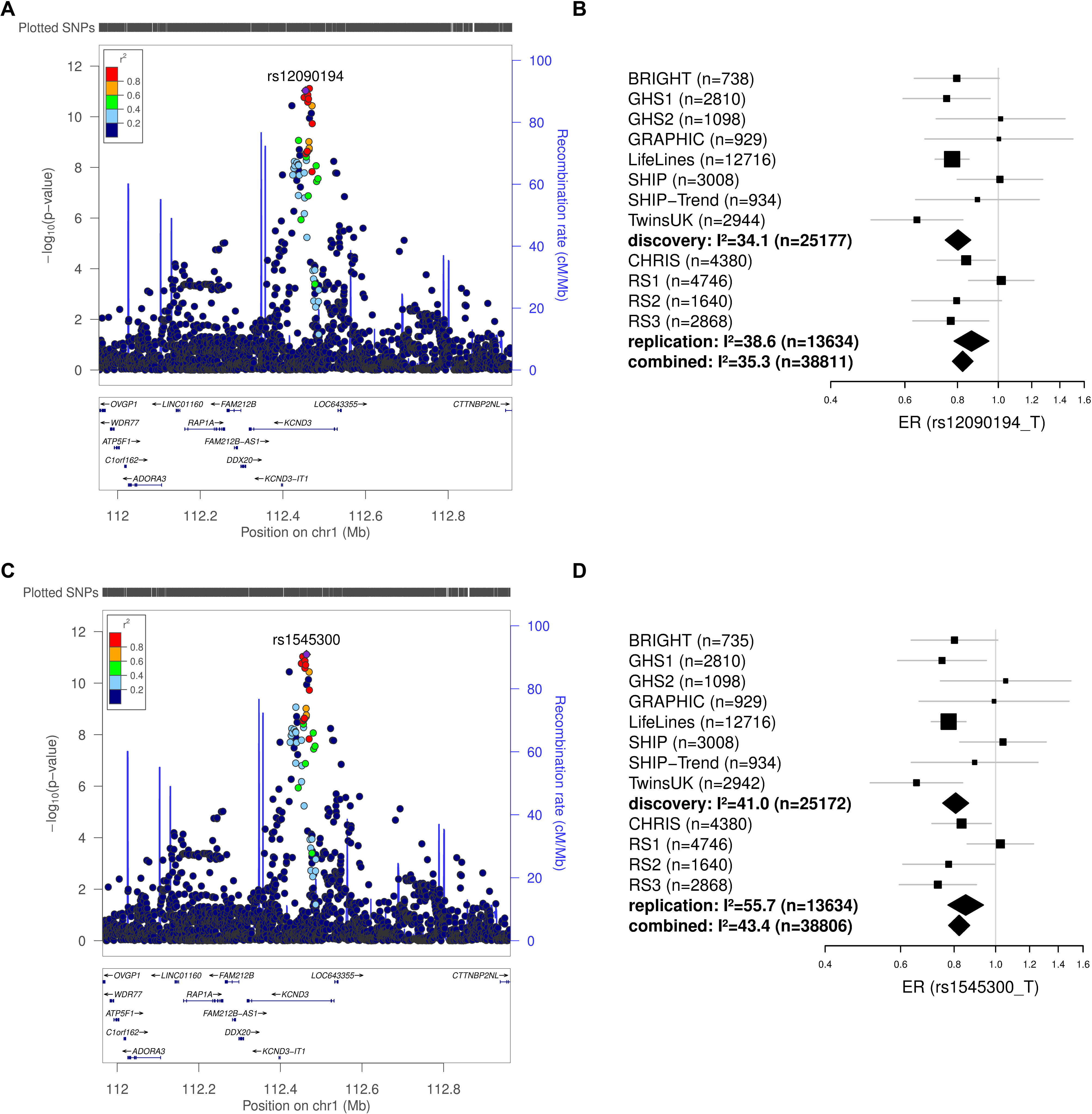
GWAS results of the *KCND3* locus. The results of the combined ERP GWAS results for the *KCND3* locus are shown for the replicated discovery stage lead SNP rs12090194 (A and B), and for the combined GWAS lead SNP rs1545300 (C and D). The regional association plots (A and C) show the association results in a ±500 kb region around the lead SNP. SNPs are plotted on the x-axis according to their chromosomal position with the −log_10_(p-value) of the GWAS association on the y-axis. Correlation with the lead SNP (purple) is estimated based on the 1000 Genomes reference samples. Plots were generated using the website of LocusZoom (Pruim, R. J. *et al.* Bioinformatics, 2010). Genetic positions refer to GRCh37/hg19 coordinates. Forest plots of the respective lead SNPs are provided in (B) and (D), with odds ratios and their 95% confidence intervals plotted on the x-axis. I² is the percentage of total variation across studies that is due to heterogeneity.

### Statistical finemapping of the associated locus

All significantly associated SNPs were located within *KCND3*, the potassium voltage-gated channel subfamily D member 3 gene and were intronic. We used the discovery and replication stage combined GWAS results to assess whether a single SNP or set of variants drive the association signal in *KCND3* (credible set). The 99% credible set was computed based on Approximate Bayes Factors for each SNP, resulting for each in a set of SNPs that with 99% posterior probability contained the variant(s) driving the association signal. For the associated locus at *KCND3* the credible set spanned 49 kb, and contained 19 variants. The two lead SNPs rs1545300 and rs12090194 had a posterior probability of 21% and 19%, respectively, whereas the former candidate SNP rs17029069(18) had a posterior probability of 2% (**Supplementary Table 2**).

To test whether the association in *KCND3* might be driven by heart rate or RR interval, we performed a sensitivity analysis in the 1,253 ERP cases and 11,463 controls of the Lifelines cohort adjusting the genetic association of rs1545300 additionally for these two traits in separate models. The effect estimates were virtually unchanged (OR=0.78) with p=1.2E-7 for both adjustments. In addition, we assessed whether the association of rs1545300 might be related to a specific ERP subtype i.e ST segment or ERP localization. In all subtype-stratified analyses the 95% confidence intervals of the effect sizes overlapped with the overall results not pointing to a subtype driven signal (**Supplementary Table 3**).

### eQTL and co-localization

We searched the Genotype-Tissue Expression (GTEx) project database(20) to look for tissue-specific eQTLs including all genes in vicinity of ±1Mb of the lead SNP rs1545300 and found an association with *KCND3* expression levels in tibial artery (p=3.0E-6, n=388). Two additional eQTL associations of rs1545300 at a false discovery rate (FDR) <0.2 across the 48 tissues tested were found with *KCND3* (ENSG00000171385.5) in the left ventricle (p=2.9E-4, n=272) of the human heart, and with *CEPT1* (ENSG00000134255.9) in the minor salivary gland (p=3.4E-4, n=97) (**Supplementary Table 4**).

Subsequent co-localization analyses of rs1545300 in these three tissues revealed also a significant correlation of gene expression pattern with ERP (p_SMR_≤0.01) (Figure 2, **Supplementary Table 5**), where for the left ventricle the correlation seems to be attributable to the same underlying causative variant (p_HEIDI_≥0.05), and for tibial artery the test was close to nominal significance (p_HEIDI_=0.05). However, the significant p_HEIDI_=1.7E-3 of *CEPT1* in the minor salivary gland points rather towards a pleiotropic effect of rs1545300 than to a causal effect of gene expression on ERP in this tissue. For all three tissues, an increased gene expression level was associated with a higher risk of ERP (**Supplementary Table 5**).

**Figure 2.**
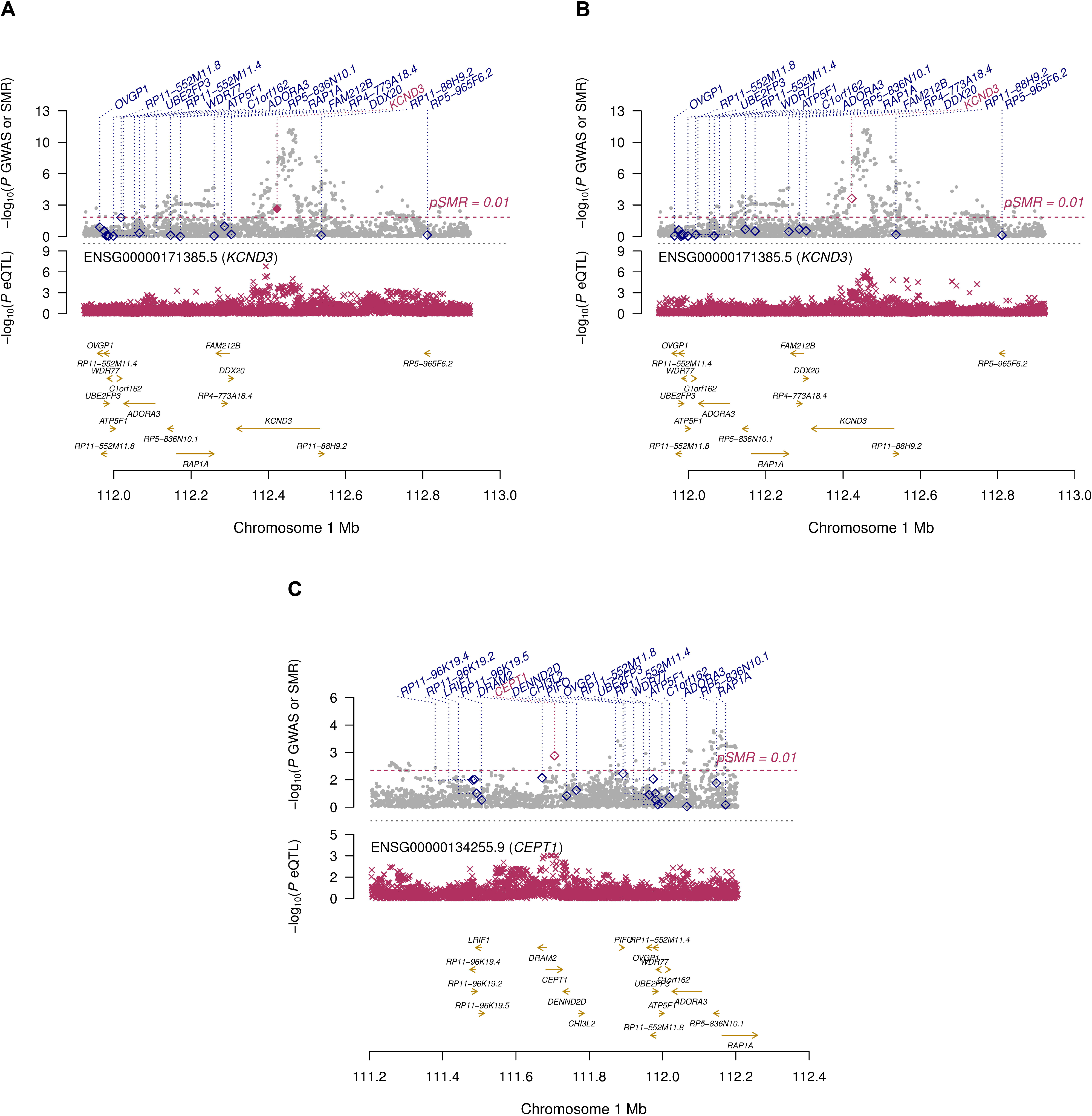
Co-localization results. Illustration of the SMR test for ERP risk and expression QTLs at the rs1545300 locus at chromosome 1p13.2 for (A) left ventricle of the heart, (B) tibial artery, and (C) minor salivary gland tissue. In each panel, the upper box shows the GWAS regional association plot with ERP risk, with level of significance of the SMR test (y-axis) for each transcript in the locus indicated by a diamond positioned at the center of the transcript. A significant SMR test represented by a purple diamond indicates an association of the transcript level of the respective genes (purple label) with the trait. For all three tissues, an increased gene expression level of a significant SMR test was associated with a higher risk of ERP. A filled purple diamond indicates a HEIDI test p-value >0.05, thus a likely co-localization. The lower box shows the regional association distribution with changes in expression of the highlighted (purple) gene transcript in the respective tissue. In both boxes, the x-axis refers to GRCh37/hg19 genomic coordinates.

### Pleiotropic effects of the lead SNPs

To assess pleiotropic effects of the *KCND3* lead SNP rs1545300 or its proxies (r²>0.8), we looked for genome-wide significant associations in the NHGRI-EBI Catalog of published genome-wide association studies(21) (accessed: 05/03/2019). Pleiotropic associations were found for P-wave terminal force (rs12090194 and rs4839185)(22) and for atrial fibrillation (rs1545300 and rs1443926)(23, 24). All these SNPs were in strong linkage disequilibrium (r²>0.97) with the lead SNP. In addition, variants in low to moderate LD with rs1545300 were associated with P-wave duration (rs2798334, r²=0.26)(25) and ST-T-wave amplitudes (rs12145374, r²=0.60)(26).

## Discussion

In this GWAS meta-analysis comprising 3,305 cases and 36,151 controls including independent replication samples, we describe an association of ERP with a locus on chromosome 1 in the *KCND3* gene. This is the first study identifying a robust genome-wide significant association between genetic variants and ERP. Our findings form the genetic basis for further functional studies examining the pathophysiological mechanism of ERP and potentially ERS. The *KCND3* gene encodes the main pore-forming alpha subunit of the voltage-gated rapidly inactivating A-type potassium channel. In the cardiac ventricle *KCND3* contributes to the fast cardiac transient outward potassium current (I_to_), which plays a major role in the early repolarization phase 1 of the cardiac action potential (AP).

To date, two competing theories explain the presence of J waves and ERP: the repolarization and the depolarization theory, both involving the I_to_ channel. On the basis of animal models evidence for the former is more compelling. Thus, J waves result from a transmural voltage gradient created by a more prominent epicardial phase 1 AP notch relative to the endocardial AP notch(10, 27). The I_to_ current notably influences the degree of the transmural heterogeneity of the phase 1 AP notch and consecutively the magnitude of the J wave(10, 27). Pharmacological inhibition of the I_to_ current with 4-aminopyridine results in a reduction of the J wave amplitude(10). The depolarization theory is based on clinical overlap of ERP with Brugada syndrome, which has led to the suggestion of Brugada syndrome being a right ventricular variant of the ERP(28). In theory, deviation from the sequential activation of cardiac currents I_Na_, I_to_, and I_CaL_ can lead to regional conduction slowing and appearance of inferior and/or lateral ERP(27, 29). In patients with ERS, distinct phenotypes of both delayed depolarization and early repolarization have been identified(30).

ERP is a highly heritable trait within families(3, 16), however limited heritability can be attributed to common SNPs in unrelated individuals(17). This might be a reason why the only GWAS to date which included 452 cases failed to replicate any genome-wide significant loci(18). In our study, which includes 3,334 cases, we discovered and replicated variants in the *KCND3* gene. Interestingly, one of these variants (rs17029069), which is in moderate LD (r²=0.18, D’=-1) with our lead SNP rs1545300 (**Supplementary Figure 3**) was reported as a candidate in the earlier GWAS meta-analysis(18). However, this variant did not replicate in their study, which the authors attributed to limited power based on the small sample size and/or heterogeneous phenotyping. In our study, experienced cardiologists centrally adjudicated more than 39,000 ECGs with high reproducibility ensuring a very high phenotyping quality(17). The resulting homogenously assessed phenotype and the substantially increased number of cases are two aspects that elevated the statistical power of our GWAS meta-analysis. All detected variants cluster in intronic regions of the *KCND3* gene, without significant allelic heterogeneity. The annotation of the locus does not point to a direct pathogenic effect, i.e. a protein altering mutation, and also the statistical finemapping revealed no single SNP with a substantial posterior probability (e.g. >80%) of being causal. However, the latter approach has limitations of detecting rare causal variants due to imputation uncertainty and minimum minor allele frequency (MAF). Nevertheless, eQTL analysis suggested that the detected variants may affect gene expression of *KCND3*. Potential mechanisms include modification of gene expression via altered binding of transcription factors at *cis*-elements. This is supported by the results of the test for co-localization showing an increase of ERP risk due to increased gene expression levels of *KCND3* in tissues of the human heart and tibial artery. Similar, pharmacological *ex vivo* data predict gain of function mutations in the I_to_ current to increase the overall transmural outward shift, leading to an increased epicardial AP notch and thereby inducing ERP in the surface ECG(27). Additionally, in close proximity to the lead SNP rs1545300 a long non-coding RNA (lncRNA), KCND3 antisense RNA 1 (*KCND3-AS1*) is described. LncRNAs have been shown to physiologically influence gene regulation through various mechanism e.g. chromatin remodeling, control of transcription initiation and post-transcriptional processing(31, 32). On the other hand, dysregulation of lncRNA control circuits can potentially impact development of disease(33): a very prominent example in cardiovascular diseases is the lncRNA *ANRIL*, which is a key effector of *9p21* in atherosclerotic risk and cardiovascular events(33–35).

Given the high prevalence of ERP in the general population and a high MAF of the identified genetic variants in our study the key question remains why only a very small subset of individuals develops severe ventricular arrhythmias and ERS. The fine interplay of a genetic predisposition and specific precipitating conditions might lead to an electrically vulnerable cardiac state. Insights into the potential origin of ventricular arrhythmias in ERS come from animal models and highlight the role of different ion channels including I_to_(36). A pharmacological model of ERS in canine wedges from the inferior and lateral ventricular wall showed marked regional dispersion of repolarization (loss of phase 2 AP dome and AP shortening in some epicardial regions but not others). Presence of transmural repolarization heterogeneity allowed local re-excitation in form of closely coupled extrasystolic activity (phase 2 re-entry). The combination of an arrhythmogenic substrate, represented by regional electrical instability, and triggering premature ventricular beats resulted in ventricular fibrillation(36). Human data in ERS patients suggest that in a subgroup, the ERP is due to a pure repolarization phenotype and arrhythmia(30) is triggered by Purkinje fiber ectopic beats.

Genetic variants in various ion channel genes have been associated with ERS(37) including the *KCNJ8* and *ABCC9* genes encoding the Kir6.1 and ATP-sensing subunits of the K_ATP_ channel(6, 11, 38, 39). The commonly implicated variant KCNJ8-p.S422L has a population frequency not consistent with ERS, and is predicted to be benign by multiple in silico algorithms according to the ClinVar database(40). A recent study by Chaveau *et al.* has, however, identified a de novo duplication of the *KCND3* gene in a patient who survived sudden cardiac death and in his 2-year-old daughter(12). Both exhibited marked ERP in the inferolateral leads that was augmented by bradycardia and pauses in heart rhythm, in keeping with a repolarization mechanism underlying the ERS phenotype. Studies have suggested that the inferior region of the left ventricle has a higher density of *KCND3* expression and higher intrinsic levels of I_to_(36). This may explain the higher vulnerability of this region for the development of ERS in the setting of a genetically mediated gain-of-function in the I_to_ current. The findings of our study therefore suggest that common variation may play a role in the expression of *KCND3* and the I_to_ current and that it is likely to be relevant in ERS as well. This may in part explain the minimal yield of pathogenic variants in ERS cases. Further GWAS in large collaborative cohorts of ERS patients are therefore necessary to determine the importance of polygenic risk. A systematic evaluation of pleiotropic effects demonstrated known associations of the identified *KCND3* SNPs with ECG phenotypes only. The lead SNP rs1545300 or its proxies in strong LD (r²>0.97) were found to be related with P-wave terminal force(22) and atrial fibrillation(23). This highlights the sharing of underlying mechanisms between *KCND3* variation and cardiac electrical activity.

Our study has some limitations, which need to be acknowledged. Presence of ERP in the ECG can be variable, as it has been described to be dependent on age, heart rate, vagal activity and medication, although our findings were valid after adjusting for some of these factors. Therefore, we cannot exclude that we have missed some individuals with ERP. Second, the tissue-specific gene expression data used for the co-localization analysis is based on a limited sample size. A larger gene expression sample or functional studies are needed to validate the revealed effect of *KCND3* expression on the ERP.

In conclusion, we show for the first time, a robust association of genetic variants with the ERP in a large GWAS of individuals of European ancestry. The locus in the *KCND3* ion channel gene is an intuitive candidate and supports the theory that at least a proportion of ERS is a pure channelopathy. Intensive future research will be needed to extend the discovery of ERP susceptibility loci to individuals of non-European ancestry, and to improve identification and risk stratification of the subset of individuals with the ERP who are at highest risk for potentially lethal ventricular arrhythmias.

## Methods

### Study cohorts and SNP genotyping

The discovery stage included 25,822 subjects (2,181 ERP cases) from eight independent cohorts: the British Genetics of Hypertension (BRIGHT) study, the Gutenberg Health Study (GHS1, GHS2), the Genetic Regulation of Arterial Pressure In humans in the Community (GRAPHIC) study, the Lifelines Cohort Study (Lifelines), the Study of Health in Pomerania (SHIP, SHIP-Trend), and TwinsUK. Additional 13,634 subjects (1,124 ERP cases) from four cohorts (Rotterdam Study I, II, III, and CHRIS) were used as independent replication: the Rotterdam Study (Rotterdam Study I, II, III), and the Cooperative Health Research In South Tyrol (CHRIS) study. The included subjects of all cohorts were of European ancestry, and all cohorts but BRIGHT (which sampled hypertensive cases) were population based (**Supplementary Table 1**). All subjects gave written informed consent and the studies were approved by the local ethics committees.

### Electrocardiogram analysis and ERP evaluation

12-lead ECGs of all 12 studies were analyzed manually by experienced and specifically trained cardiologists for the presence of ERP according to the established definition by Haissaguerre and Macfarlane(6, 19). In case of a QRS duration of >120 ms or rhythm other than sinus rhythm (e.g. atrial fibrillation, pacemaker stimulation) ECGs were excluded from the analysis. The methodology employed and robustness of inter-observer correlations have been presented elsewhere(17).

In detail, ERP was defined as elevation of the J-point above the level of QRS onset of ≥0.1 mV in at least two corresponding leads. To avoid confusion or overlap with Brugada syndrome or arrhythmogenic right ventricular dysplasia, leads V1 to V3 were excluded from ERP scoring. In case of presence of ERP, region, either inferior (leads II, III, aVF), antero-lateral (leads I, aVL, V_4_-V_6_), or both, and the maximum amplitude of J-point elevation was documented. Further, the morphology of ERP was assessed as either notching, slurring or both as well as the ST segment according to Tikkanen and collegues(41) as either concave/rapidly ascending (>0.1 mV elevation 100 ms after J-point peak or persistently elevated ST segment >0.1 mV) or horizontal/descending (≤0.1 mV elevation within 100 ms after J-point peak)(19, 41).

### GWAS in individual studies

The GWAS in each study for both the discovery and replication stage was performed on autosomal imputed SNP genotypes using study-specific quality control protocols which are provided in detail in **Supplementary Table 1**. Association analyses were performed using logistic regression for ERP status as outcome and an additive genetic model on SNP dosages, thus taking genotype uncertainties of imputed SNPs into account. The analyses were adjusted for age, sex, and relevant study-specific covariates such as principal components for population stratification (**Supplementary Table 1**).

### Statistical methods for meta-analysis

The result files from individual studies GWAS underwent extensive quality control before meta-analysis using the gwasqc() function of the GWAtoolbox package v2.2.4(42). The quality control included file format checks as well as plausibility and distributions of association results including effect sizes, standard errors, allele frequencies and imputation quality of the SNPs.

The meta-analyses were conducted using a fixed-effect inverse variance weighting as implemented in Metal(43). Monomorphic SNPs, SNPs with implausible association results (i.e. p≤0, SE≤0, |log(OR)|≥1000), and SNPs with an imputation quality score ≤0.4 were excluded prior to the meta-analyses resulting in a median of 12,839,202 SNPs per cohort (IQR: 10,756,073-13,184,807). During the meta-analysis, the study-specific results were corrected by their specific λ_GC_ if >1. Results were checked for possible errors like use of incorrect association model by plotting the association p-values of the analyses against those from a z-score based meta-analysis for verifying overall concordance. SNPs that were present in <75% of the total sample size contributing to the respective meta-analysis or with a MAF ≤0.01 were excluded from subsequent analyses. Finally, data for up to 6,976,246 SNPs were available after the meta-analysis.

Quantile-quantile plots of the meta-analysis results are provided in **Supplementary Figure 2**. To assess whether there was an inflation of p-values in the meta-analysis results attributed to reasons other than polygenicity, we performed LD score regression(44). The LD score corrected λ_GC_ value of the discovery and replication combined meta-analysis was 1.01, supporting the absence of unaccounted population stratification. Genome-wide significance was defined as a p-value <5E-8, corresponding to a Bonferroni correction of one million independent tests. Unless stated otherwise, all reported p-values are two-sided. The I^2^ statistic was used to evaluate between-study heterogeneity(45).

To evaluate the presence of allelic heterogeneity within each locus, the GCTA stepwise model selection procedure (cojo-slct algorithm) was used to identify independent variants employing a step-wise forward selection approach(46). We used the genotype information of 4,081 SHIP individuals for LD estimation, and set the significance threshold for independent SNPs to 5E-8.

All loci were named according to the nearest gene of the lead SNP. Genomic positions correspond to build 37 (GRCh37).

### Replication analysis

To minimize the burden for multiple testing correction and thus maximizing the power for replication, the lead SNPs of genome-wide significant loci in the discovery stage were taken forward to the replication stage in independent samples (Table 1). SNPs were considered as replicated if the p-value of a one-sided association test was <0.025 which corresponds to a Bonferroni correction for the two lead SNPs tested at 5% significance level.

Finally, the GWAS results from the discovery and replication studies were meta-analyzed to search for additional genome-wide significant loci by maximizing the statistical power for locus discovery.

### Gene expression based analyses

The lead SNP rs1545300 of the *KCND3* locus of the combined discovery and replication GWAS meta-analysis was tested for *cis* eQTLs (±1Mb window around the transcription start site) in 48 tissues available in the GTEx v7 database that included at least 70 samples. Significant associations were selected based on a Bonferroni corrected p-value <3.0E-5 for the number of genes and tissues tested. Subsequently, the SNP rs1545300 was tested and plotted for co-localization in the three tissues with an eQTL FDR<0.2 by applying the SMR method(47) using the GWAS and GTEx eQTL summary statistics. The method includes a test whether the effect on expression observed at a SNP or at its proxies is independent of the signal observed in the GWAS, i.e. that gene expression and y are associated only because of a latent non-genetic confounding variable (SMR test), and a second test that evaluates if the eQTL and GWAS associations can be attributable to the same causative variant (HEIDI test). Significance for co-localization of the gene expression and the GWAS signals was defined by p_SMR_<0.01, where additionally a p_HEIDI_≥0.05 indicates the same underlying causal variant(47).

## Acknowledgments

Detailed acknowledgments are provided in the Supplementary Information.

## Author Contributions

Project design and analysis: W.R., T.T., A.T., M.D, E.R.B. Management of individual study: A.F.D., C.F., C.P., E.R.B., K.J.L., M.A.I., M.D., M.G., M.Z., N.P., N.V., P.B.M., P.P., P.v.d.H., S.A., S.B.F., T.D.S., T.M., T.T., W.R., Y.J. Recruitment of individual study subjects: A.F.D., G.S., H.S., I.D., M.D., M.Z., N.V., P.v.d.H., S.A., S.B.F. Drafting of the manuscript: A.R., A.T., B.H.S., E.R.B., M.D.B., M.E.v.d.B., T.T., W.R. Statistical methods and analysis of individual study: A.T., C.C., C.F., H.R.W., H.S., I.M.N., K.S., M.E.v.d.B., S.P. Genotyping of individual study: A.G.U., C.F., M.N., P.B.M., U.V. Interpretation of the results: A.R., A.T., B.K., C.H., E.R.B., M.D., M.D.B., T.K., T.T., W.R. Critical review of the manuscript: all authors.

## Data availability

Summary genetic association results have been submitted for full download to the CHARGE dbGaP website under accession phs000930 [https://www.ncbi.nlm.nih.gov/gap].

