## Supplementary Data for "*KCND3* is a novel susceptibility locus for early repolarization"

SUPPLEMENTARY INFORMATION

### Supplementary Text

#### *Acknowledgments*

The Genotype-Tissue Expression (GTEx) Project was supported by the Common Fund of the Office of the Director of the National Institutes of Health, and by NCI, NHGRI, NHLBI, NIDA, NIMH, and NINDS.

*BRIGHT*

The BRIGHT study is extremely grateful to all the patients who participated in the study and the BRIGHT nursing team. This works forms part of the research areas contributing to the translational research portfolio of the Cardiovascular Biomedical Research Centre at Barts which is supported and funded by the National Institute for Health Research (Warren, Cabrera, Munroe). This work was supported by the Medical Research Council of Great Britain (grant number G9521010D) and the British Heart Foundation (grant number PG/02/128).

*CHRIS*

Full acknowledgements for the Cooperative Health Research In South Tyrol (CHRIS) study are reported here: http://translational-medicine.biomedcentral.com/articles/10.1186/s12967-015-0704-9#Declarations. The CHRIS study was funded by the Department of Innovation, Research, and University of the Autonomous Province of Bolzano-South Tyrol.

*GHS*

Gutenberg Health Study: The Gutenberg Health Study is funded through the government of Rhineland-Palatinate („Stiftung Rheinland-Pfalz für Innovation“, contract AZ 961-386261/733), the research programs “Wissen schafft Zukunft” and “Center for Translational Vascular Biology (CTVB)” of the Johannes Gutenberg-University of Mainz, and its contract with Boehringer Ingelheim and PHILIPS Medical Systems, including an unrestricted grant for the Gutenberg Health Study. Philipp S. Wild is funded by the Federal Ministry of Education and Research (BMBF 01EO1503) and he is PI of the German Center for Cardiovascular Research (DZHK). This project has received funding from the European Research Council (ERC) under the European Union’s Horizon 2020 research and innovation programme (grant agreement No 648131). This work was performed in the context of the Junior Research Alliance symAtrial project funded by the German Ministry of Research and Education (BMBF 01ZX1408A) e:Med – Systems Medicine program (RBS). RBS was supported by Deutsche Forschungsgemeinschaft (German Research Foundation) Emmy Noether Program SCHN 1149/3-1.

*GRAPHIC*

The GRAPHIC study was funded by the BHF.

*Lifelines*

Lifelines is a multi-disciplinary prospective population-based cohort study examining in a unique three-generation design the health and health-related behaviors of 167,729 persons living in the North of The Netherlands. It employs a broad range of investigative procedures in assessing the biomedical, socio-demographic, behavioral, physical and psychological factors which contribute to the health and disease of the general population, with a special focus on multi-morbidity and complex genetics.

The Lifelines Cohort Study, and generation and management of GWAS genotype data for the Lifelines Cohort Study is supported by the Netherlands Organization of Scientific Research NWO (grant 175.010.2007.006), the Economic Structure Enhancing Fund (FES) of the Dutch government, the Ministry of Economic Affairs, the Ministry of Education, Culture and Science, the Ministry for Health, Welfare and Sports, the Northern Netherlands Collaboration of Provinces (SNN), the Province of Groningen, University Medical Center Groningen, the University of Groningen, Dutch Kidney Foundation and Dutch Diabetes Research Foundation.

The authors wish to acknowledge the services of the Lifelines Cohort Study, the contributing research centers delivering data to Lifelines, and all the study participants.

Lifelines Cohort Study group authors:

Behrooz Z Alizadeh (1), H Marike Boezen (1), Lude Franke (2), Pim van der Harst (3), Gerjan Navis (4), Marianne G Rots (5), Harold Snieder (1), Morris Swertz (2), Bruce HR Wolffenbuttel (6), Cisca Wijmenga (2)

*(1) Department of Epidemiology, University of Groningen, University Medical Center Groningen, The Netherlands*
*(2) Department of Genetics, University of Groningen, University Medical Center Groningen, The Netherlands*
*(3) Department of Cardiology, University of Groningen, University Medical Center Groningen, The Netherlands*
*(4) Department of Internal Medicine, Division of Nephrology, University of Groningen, University Medical Center Groningen, The Netherlands*
*(5) Department of Pathology and Medical Biology, University of Groningen, University Medical Center Groningen, The Netherlands*
*(6) Department of Endocrinology, University of Groningen, University Medical Center Groningen, The Netherlands*

*Rotterdam Study*

The generation and management of GWAS genotype data for the Rotterdam Study (RS I, RS II, RS III) was executed by the Human Genotyping Facility of the Genetic Laboratory of the Department of Internal Medicine, Erasmus MC, Rotterdam, The Netherlands. The GWAS datasets are supported by the Netherlands Organisation of Scientific Research NWO Investments (nr. 175.010.2005.011, 911-03-012), the Genetic Laboratory of the Department of Internal Medicine, Erasmus MC, the Research Institute for Diseases in the Elderly (014-93-015; RIDE2), the Netherlands Genomics Initiative (NGI)/Netherlands Organisation for Scientific Research (NWO) Netherlands Consortium for Healthy Aging (NCHA), project nr. 050-060-810. We thank Pascal Arp, Mila Jhamai, Marijn Verkerk, Lizbeth Herrera and Marjolein Peters, MSc, and Carolina Medina-Gomez, MSc, for their help in creating the GWAS database, and Karol Estrada, PhD, Yurii Aulchenko, PhD, and Carolina Medina-Gomez,

MSc, for the creation and analysis of imputed data. The Rotterdam Study is funded by Erasmus Medical Center and Erasmus University, Rotterdam, Netherlands Organization for the Health Research and Development (ZonMw), the Research Institute for Diseases in the Elderly (RIDE), the Ministry of Education, Culture and Science, the Ministry for Health, Welfare and Sports, the European Commission (DG XII), and the Municipality of Rotterdam. The authors are grateful to the study participants, the staff from the Rotterdam Study and the participating general practitioners and pharmacists.

*SHIP*

The Study of Health in Pomerania (SHIP) is part of the Community Medicine Research net of the University of Greifswald, Germany, which is funded by the Federal Ministry of Education and Research (grants no. 01ZZ9603, 01ZZ0103, and 01ZZ0403), the Ministry of Cultural Affairs as well as the Social Ministry of the Federal State of Mecklenburg-West Pomerania, and the network ‘Greifswald Approach to Individualized Medicine (GANI_MED)’ funded by the Federal Ministry of Education and Research (grant 03IS2061A). Genome-wide data have been supported by the Federal Ministry of Education and Research (grant no. 03ZIK012) and a joint grant from Siemens Healthineers, Erlangen, Germany and the Federal State of Mecklenburg- West Pomerania. The University of Greifswald is a member of the Caché Campus program of the InterSystems GmbH.

*TwinsUK*

The study was funded by the Wellcome Trust; European Community’s Seventh Framework Programme (FP7/2007-2013). The study also receives support from the National Institute for Health Research (NIHR) BioResource Clinical Research Facility and Biomedical Research Centre based at Guy's and St Thomas' NHS Foundation Trust and King's College London and the British Heart Foundation. Tim Spector is holder of an ERC Advanced Principal Investigator award. SNP Genotyping was performed by The Wellcome Trust Sanger Institute and National Eye Institute via NIH/CIDR. Statistical analyses were carried out on the Genetic Cluster Computer (http://www.geneticcluster.org) hosted by SURFsara and financially supported by the Netherlands Scientific Organization (NWO 480-05-003 PI: Posthuma) along with a supplement from the Dutch Brain Foundation and the VU University Amsterdam. Y.J. received funding for the project from the British Heart Foundation (PG/12/38/29615 and PG/06/094/21278). E.B. receives funding from the Robert Lancaster Memorial Fund sponsored by McColl's RG Ltd.

### Supplementary Tables

#### *Supplementary Table 1: Study genotyping information*

| | Study | BRIGHT | CHRIS | GHS1 | GHS2 | GRAPHIC |
|---|---|---|---|---|---|---|
| | Ethnicity | European | European | European | European | European |
| | Country | UK | Italy | Germany | Germany | United Kingdom |
| | Study design | Hypertensive cases | Population-based | Population-based | Population-based | Nuclear families (Offspring excluded) |
| GENOTYPING | Genotyping centre | Affymetrix, Inc., USA and Wellcome Trust Sanger Inst.(UK) | Life and Brain, Bonn (Germany) | Affymetrix, Inc., USA | Affymetrix, Inc., USA | |
| | Genotyping Array | Affymetrix GeneChip 500k array | Illumina Human OmniExpressExome | Affymetrix SNP 6.0 | Affymetrix SNP 6.0 | Illumina human omniexpress 12 V1 |
| | Genotyping calling algorithm | CHIAMO | GenomeStudio, Z call | Birdseed2 | Birdseed2 | Illumina Genomestudio |
| SAMPLE GENOTYPING QC | Sample call rate | ≥0.97 | >99% | >95% | >95% | |
| | Other exclusions | | gender mismatch, excess of heterozygosity, excess of mendelian errors | gender mismatch, excess of IBS (> 95%), excess of heterozygosity (hetFDR > 0.01) | gender mismatch, excess of IBS (> 95%), excess of heterozygosity (hetFDR > 0.01) | |
| SNP QC | MAF (required) | 0% | NA | NA | NA | >1% |
| | HWE (required) | ≥1E-7 | P>10^-6 | 10^-4 | 10^-4 | p>10-6 |
| | Call rate | ≥95% | >99% | >98% | >98% | <95% (or<99% if SNP has MAF<5%) |
| | Exclude duplicates | NA | NA | NA | NA | NA |
| | Other | NA | NA | NA | NA | NA |
| | SNPs for imputation | 446,472 | 610,883 | 662,405 | 673,914 | 612,432 |
| Imputation | Reference Panel | 1000G March 2012 | 1000G Ph1 v3 | 1000G Ph1 v2 | 1000G Ph1 v2 | 1000Gv3 |
| | Build | 37 | 37 | 37 | 37 | 37 |
| | Software for imputation | Minimac | minimac 2 | MACH, minimac | MACH, minimac | IMPUTE v2 |
| | Filters | Imputation quality > 0.1 | Rsq < 0.3 | none | none | MAF>0 |
| | SNPs for analysis | | 14,898,673 | 15,725,774 | 15,616,273 | 14,551,978 |
| Data Analysis | Adjustments | sex, age | sex, age | sex, age | sex, age | sex, age |
| | Software for analysis | mach2qtl | SAIGE | SNPtest | SNPtest | SNPTEST |
| | Lambda GC | 1.01 | 1.01 | 0.93 | 0.85 | 0.81 |
| REFERENCES | Reference study description (PMID) | 12826435 | 26541195, 28374192 | | | 18443236 |
| | Study link/ website | NA | http://www.eurac.edu/en/research/health/biomed/projects/Pages/default.aspx | NA | NA | www2.le.ac.uk/projects/bru/our-research/research-themes/genetics-and-biomarkers/graphic-2 |

| | Study | Lifelines | Rotterdam Study I | Rotterdam Study II | Rotterdam Study III |
|---|---|---|---|---|---|
| | Ethnicity | European | European | European | European |
| | Country | The Netherlands | The Netherlands | The Netherlands | The Netherlands |
| | Study design | Population-based | General Population-Based | General Population-Based | General Population-Based |
| GENOTYPING | Genotyping centre | UMCG | Erasmus University Medical Center (Rotterdam, the Netherlands) | Erasmus University Medical Center (Rotterdam, the Netherlands) | Erasmus University Medical Center (Rotterdam, the Netherlands) |
| | Genotyping Array | Illumina Cyto SNP12 v2 | Illumina 550 duo | Illumina 550 duo | Illumina 610 quad |
| | Genotyping calling algorithm | Illumina Genomestudio | Beadstudio Genecall | Beadstudio Genecall | Beadstudio Genecall |
| SAMPLE GENOTYPING QC | Sample call rate | >95% | ≥97.5% | ≥97.5% | ≥97.5% |
| | Other exclusions | close relatives (pi-hat>0.4), non-caucasian, sex mismatch | 1) Missing DNA<br>2) Gender mismatch with typed X-linked markers<br>3) Excess autosomal heterozygosity > 0.336~FDR>0.1%<br>4) Duplicates and/or 1st or 2nd degree relatives using IBS probabilities >97% from PLINK<br>5) Ethnic outliers using IBS distances > 3SD | | |
| SNP QC | MAF (required) | >1% | >1% | >1% | >1% |
| | HWE (required) | 10-4 | P>10-6 | P>10-6 | P>10-6 |
| | CALL RATE (required) | >95% | ≥97.5% | ≥97.5% | ≥97.5% |
| | Exclude duplicates | NA | NA | NA | NA |
| | Other | NA | NA | NA | NA |
| | SNPs for imputation | 257,581 | 512,849 | 466,389 | 514,073 |
| Imputation | Reference Panel | 1000G v3 | 1000Gv3 | 1000Gv3 | 1000Gv3 |
| | Build | 37 | 37 | 37 | 37 |
| | Software for imputation | BEAGLE v.3.1.0 | MACH | MACH | MACH |
| | Filters | MAF>0 | none | none | none |
| | SNPs for analysis | 28.681.763 | 29,079,054 | 29,079,054 | 29,079,054 |
| Data Analysis | Adjustments | sex, age, PC1-6 | sex, age | sex, age | sex, age |
| | Software for analysis | PLINK | ProbABEL | ProbABEL | ProbABEL |
| | Lambda GC | 0.99 | 0.88 | 0.92 | 0.89 |
| REFERENCES | Reference study description (PMID) | 18075776, 25502107 | 1833235, 29064009 | 1833235, 29064009 | 1833235, 29064009 |
| | Study link/ website | www.lifelines.nl | http://www.epib.nl/research/ergo.htm | http://www.epib.nl/research/ergo.htm | http://www.epib.nl/research/ergo.htm |

| | Study | SHIP | SHIP-Trend | TwinsUK |
|---|---|---|---|---|
| | Ethnicity | European | European | European |
| | Country | Germany | Germany | United Kingdom |
| | Study design | Population-based | Population-based | Twins |
| GENOTYPING | Genotyping centre | Affymetrix, Inc., USA | Helmholtz Zentrum München, Germany | Wellcome Trust Sanger Inst.(UK), CIDR (USA), Centre National de Genotypage (France), Duke University (USA), Helsinki University (Finland) |
| | Genotyping Array | Affymetrix SNP 6.0 | Illumina Omni 2.5 | Illumina TruSeq, Illumina Hap300, Hap550, Hap610 |
| | Genotyping calling algorithm | Birdseed2 | GenCall v1.0 | Illuminus |
| SAMPLE GENOTYPING QC | Sample call rate | >92% | ≥94% | ≥95% |
| | Other exclusions | duplicate samples (by IBS) or reported/genotyped gender mismatch | duplicate samples (by IBS) or reported/genotyped gender mismatch | gender mismatch with X chromosomes genotypes |
| SNP QC | MAF (required) | NA | >0% | >0 % |
| | HWE (required) | P>0.0001 | P>0.0001 | P>10-6 |
| | CALL RATE (required) | >80% | >90% | >95% |
| | Exclude duplicates | chr:position:type | chr:position:type | NA |
| | Other | position mapping problem from b36 to b37 | position mapping problem from b36 to b37 | NA |
| | SNPs for imputation | 905,910 | 1,824,743 | >303,940 |
| Imputation | Reference Panel | 1000Gv3 | 1000Gv3 | 1000Gv3 |
| | Build | 37 | 37 | 37 |
| | Software for imputation | IMPUTE v2.2.2 | IMPUTE v2.2.2 | IMPUTE2 |
| | Filters | eur.maf=0 | eur.maf=0 | Info score >0.4 (imputed) |
| | SNPs for analysis | 17,533,349 | 17,585,496 | 37,426,733 |
| Data Analysis | Adjustments | sex, age | sex, age | sex, age, assay |
| | Software for analysis | QUICKTEST v0.95 | QUICKTEST v0.95 | SNPTEST2 |
| | Lambda GC | 0.90 | 0.86 | 1.03 |
| REFERENCES | Reference study description (PMID) | 20167617 | 20167617 | 17254428 |
| | Study link/ website | http://ship.community-medicine.de/ | http://ship.community-medicine.de/ | http://www.twinsuk.ac.uk |

***Supplementary Table 2: Variants of the KCND3 locus mapping into a 99% credible set***

| SNP | chr | position | A1 | A2 | AF1 | Effect | SE | OR | P | I² | N | nearest gene | function | PP | cumulated PP |
|---|---|---|---|---|---|---|---|---|---|---|---|---|---|---|---|
| rs1545300 | 1 | 112,464,004 | T | C | 0.32 | -0.1961 | 0.0287 | 0.82 | 7.7E-12 | 43.4 | 38806 | *KCND3* | intron | 0.215 | 0.215 |
| rs12090194 | 1 | 112,454,822 | T | C | 0.32 | -0.1949 | 0.0286 | 0.82 | 9.3E-12 | 35.3 | 38811 | *KCND3* | intron | 0.193 | 0.408 |
| rs4839185 | 1 | 112,460,262 | T | C | 0.68 | 0.1929 | 0.0285 | 1.21 | 1.4E-11 | 38.6 | 39293 | *KCND3* | intron | 0.146 | 0.554 |
| rs2120436 | 1 | 112,451,447 | T | C | 0.32 | -0.1938 | 0.0288 | 0.82 | 1.7E-11 | 31.2 | 38805 | *KCND3* | intron | 0.112 | 0.666 |
| rs1443926 | 1 | 112,461,902 | A | G | 0.68 | 0.1918 | 0.0286 | 1.21 | 2.0E-11 | 38.1 | 38799 | *KCND3* | intron | 0.099 | 0.765 |
| rs4839184 | 1 | 112,460,221 | C | G | 0.32 | -0.1909 | 0.0286 | 0.83 | 2.6E-11 | 37.6 | 39307 | *KCND3* | intron | 0.082 | 0.846 |
| rs2010749 | 1 | 112,470,581 | T | C | 0.63 | 0.1835 | 0.0277 | 1.20 | 3.7E-11 | 42.6 | 39229 | *KCND3* | intron | 0.065 | 0.911 |
| rs12119724 | 1 | 112,468,814 | T | C | 0.29 | 0.1902 | 0.0292 | 1.21 | 7.3E-11 | 5.5 | 38772 | *KCND3* | intron | 0.029 | 0.940 |
| rs17029069 | 1 | 112,464,376 | T | C | 0.30 | 0.1875 | 0.0291 | 1.21 | 1.1E-10 | 1.6 | 38786 | *KCND3* | intron | 0.019 | 0.960 |
| rs1443927 | 1 | 112,471,029 | C | G | 0.69 | 0.1864 | 0.0292 | 1.20 | 1.9E-10 | 42.6 | 38818 | *KCND3* | intron | 0.014 | 0.973 |
| rs6682872 | 1 | 112,462,984 | A | G | 0.60 | 0.1649 | 0.027 | 1.18 | 9.6E-10 | 39.2 | 38918 | *KCND3* | intron | 0.003 | 0.977 |
| rs2813865 | 1 | 112,437,956 | A | G | 0.25 | -0.1964 | 0.032 | 0.82 | 8.6E-10 | 66.0 | 39360 | *KCND3* | intron | 0.003 | 0.979 |
| rs583731 | 1 | 112,421,854 | T | C | 0.09 | -0.3726 | 0.0563 | 0.69 | 3.6E-11 | 43.9 | 38199 | *KCND3* | intron | 0.002 | 0.981 |
| rs4838927 | 1 | 112,463,617 | T | C | 0.60 | 0.1623 | 0.027 | 1.18 | 1.7E-09 | 42.1 | 39230 | *KCND3* | intron | 0.002 | 0.983 |
| rs4838926 | 1 | 112,463,323 | C | G | 0.40 | -0.1618 | 0.027 | 0.85 | 2.0E-09 | 41.1 | 39230 | *KCND3* | intron | 0.002 | 0.985 |
| rs7539683 | 1 | 112,439,770 | T | G | 0.27 | 0.1806 | 0.0301 | 1.20 | 2.0E-09 | 0.0 | 38700 | *KCND3* | intron | 0.001 | 0.987 |
| rs72694603 | 1 | 112,458,893 | T | C | 0.32 | -0.1805 | 0.0302 | 0.83 | 2.3E-09 | 30.2 | 35129 | *KCND3* | intron | 0.001 | 0.988 |
| rs72692602 | 1 | 112,458,833 | T | C | 0.68 | 0.1801 | 0.0302 | 1.20 | 2.5E-09 | 29.8 | 35129 | *KCND3* | intron | 0.001 | 0.989 |
| rs72692597 | 1 | 112,455,442 | T | G | 0.32 | -0.1796 | 0.0302 | 0.84 | 2.9E-09 | 31.0 | 35129 | *KCND3* | intron | 0.001 | 0.990 |

A1: effect allele; AF1: allele frequency of A1; Effect: association effect (log OR) of A1; SE: standard error of the effect; OR: odds ratio; P: association p-value; I²: percentage of total variation across studies that is due to heterogeneity; N: sample size; PP: posterior probability of being causal.

***Supplementary Table 3: Association results of rs1545300 stratified by ERP subtypes in the LifeLines cohort***

| Subtype | OR | lower CI | upper CI | P | N cases | N controls |
|---|---|---|---|---|---|---|
| *not stratified* | 0.78 | 0.71 | 0.85 | 1.18E-07 | 1253 | 11463 |
| ST segment ascending | 0.76 | 0.67 | 0.87 | 5.50E-05 | 622 | 11463 |
| ST segment horizontal/descending | 0.79 | 0.69 | 0.90 | 2.66E-04 | 630 | 11463 |
| ERP localization inferior | 0.81 | 0.71 | 0.93 | 2.49E-03 | 560 | 11463 |
| ERP localization lateral | 0.70 | 0.56 | 0.87 | 1.18E-03 | 229 | 11463 |
| ERP localization inferior and lateral | 0.77 | 0.67 | 0.90 | 7.05E-04 | 464 | 11463 |

OR: odds ratio related to minor T allele; CI: 95% confidence interval of the OR; P: association p-value; N cases, N controls: number of cases and controls, respectively.

***Supplementary Table 4: GTEx cis eQTL lookup results of rs1545300 across 48 tissues***

The GTEx variant id 1_112464004_C_T_b37 corresponds to rs1545300, and the transcript ENSG00000171385.5 to *KCND3*. The effect direction of the eQTL slope corresponds to the T allele of rs1545300.

| tissue | transcript | tss_distance | N | MAC | P | slope | slope_se | FDR |
|---|---|---|---|---|---|---|---|---|
| Artery_Tibial | ENSG00000171385.5 | -67773 | 441 | 219 | 2.98E-06 | -0.19 | 0.04 | 0.01 |
| Heart_Left_Ventricle | ENSG00000171385.5 | -67773 | 303 | 154 | 2.92E-04 | -0.12 | 0.03 | 0.19 |
| Minor_Salivary_Gland | ENSG00000134255.9 | 781156 | 97 | 49 | 3.39E-04 | -0.34 | 0.09 | 0.19 |
| Heart_Atrial_Appendage | ENSG00000171385.5 | -67773 | 297 | 147 | 1.58E-03 | -0.13 | 0.04 | 0.49 |
| Skin_Not_Sun_Exposed_Suprapubic | ENSG00000155367.11 | -794095 | 387 | 200 | 1.89E-03 | 0.10 | 0.03 | 0.49 |
| Adipose_Visceral_Omentum | ENSG00000231346.1 | 312659 | 355 | 172 | 1.91E-03 | -0.15 | 0.05 | 0.49 |
| Brain_Amygdala | ENSG00000121933.13 | 357420 | 100 | 47 | 2.03E-03 | 0.23 | 0.07 | 0.49 |
| Artery_Aorta | ENSG00000231346.1 | 312659 | 299 | 152 | 2.94E-03 | 0.15 | 0.05 | 0.62 |
| Esophagus_Muscularis | ENSG00000064886.9 | 720611 | 370 | 188 | 4.12E-03 | 0.13 | 0.05 | 0.71 |
| Brain_Anterior_cingulate_cortex_BA24 | ENSG00000155367.11 | -794095 | 121 | 65 | 4.60E-03 | 0.22 | 0.07 | 0.71 |
| Whole_Blood | ENSG00000155363.14 | -751759 | 407 | 195 | 4.93E-03 | -0.06 | 0.02 | 0.71 |
| Heart_Left_Ventricle | ENSG00000143110.7 | 447590 | 303 | 154 | 6.14E-03 | -0.15 | 0.05 | 0.71 |
| Esophagus_Muscularis | ENSG00000116473.10 | 379164 | 370 | 188 | 6.60E-03 | -0.09 | 0.03 | 0.71 |
| Lung | ENSG00000155366.12 | -786052 | 427 | 222 | 6.67E-03 | -0.07 | 0.02 | 0.71 |
| Artery_Aorta | ENSG00000064886.9 | 720611 | 299 | 152 | 6.72E-03 | 0.17 | 0.06 | 0.71 |
| Brain_Caudate_basal_ganglia | ENSG00000134245.13 | -545159 | 160 | 88 | 7.52E-03 | 0.14 | 0.05 | 0.71 |
| Colon_Sigmoid | ENSG00000121933.13 | 357420 | 233 | 107 | 7.62E-03 | -0.18 | 0.07 | 0.71 |
| Minor_Salivary_Gland | ENSG00000173947.9 | 575094 | 97 | 49 | 8.76E-03 | 0.22 | 0.08 | 0.71 |
| Ovary | ENSG00000260948.1 | 488314 | 133 | 71 | 8.89E-03 | -0.23 | 0.09 | 0.71 |
| Whole_Blood | ENSG00000116489.8 | -698415 | 407 | 195 | 9.00E-03 | -0.04 | 0.01 | 0.71 |
| Brain_Caudate_basal_ganglia | ENSG00000273010.1 | 958080 | 160 | 88 | 9.92E-03 | 0.34 | 0.13 | 0.71 |
| Ovary | ENSG00000085465.11 | 493605 | 133 | 71 | 1.01E-02 | -0.29 | 0.11 | 0.71 |
| Colon_Sigmoid | ENSG00000155363.14 | -751759 | 233 | 107 | 1.02E-02 | -0.12 | 0.05 | 0.71 |
| Brain_Anterior_cingulate_cortex_BA24 | ENSG00000134255.9 | 781156 | 121 | 65 | 1.02E-02 | 0.16 | 0.06 | 0.71 |
| Esophagus_Gastroesophageal_Junction | ENSG00000227811.2 | 181541 | 244 | 110 | 1.09E-02 | -0.24 | 0.09 | 0.71 |
| Brain_Spinal_cord_cervical_c-1 | ENSG00000232811.1 | 977915 | 91 | 49 | 1.21E-02 | 0.48 | 0.18 | 0.71 |
| Colon_Sigmoid | ENSG00000007341.14 | -699443 | 233 | 107 | 1.27E-02 | -0.22 | 0.09 | 0.71 |
| Testis | ENSG00000155367.11 | -794095 | 259 | 128 | 1.32E-02 | 0.07 | 0.03 | 0.71 |
| Small_Intestine_Terminal_Ileum | ENSG00000197852.8 | 168128 | 137 | 54 | 1.46E-02 | 0.22 | 0.09 | 0.71 |
| Brain_Spinal_cord_cervical_c-1 | ENSG00000260948.1 | 488314 | 91 | 49 | 1.47E-02 | -0.36 | 0.14 | 0.71 |
| Breast_Mammary_Tissue | ENSG00000064886.9 | 720611 | 290 | 148 | 1.52E-02 | -0.17 | 0.07 | 0.71 |
| Artery_Aorta | ENSG00000171385.5 | -67773 | 299 | 152 | 1.61E-02 | -0.14 | 0.06 | 0.71 |
| Brain_Cerebellum | ENSG00000143079.10 | -474799 | 173 | 96 | 1.62E-02 | 0.18 | 0.07 | 0.71 |
| Brain_Amygdala | ENSG00000116459.6 | 472490 | 100 | 47 | 1.63E-02 | -0.15 | 0.06 | 0.71 |
| Adipose_Subcutaneous | ENSG00000116459.6 | 472490 | 442 | 214 | 1.64E-02 | 0.08 | 0.03 | 0.71 |
| Adrenal_Gland | ENSG00000273483.1 | -597059 | 190 | 96 | 1.64E-02 | 0.19 | 0.08 | 0.71 |
| Pituitary | ENSG00000064886.9 | 720611 | 183 | 95 | 1.64E-02 | 0.17 | 0.07 | 0.71 |
| Brain_Anterior_cingulate_cortex_BA24 | ENSG00000121933.13 | 357420 | 121 | 65 | 1.72E-02 | 0.18 | 0.07 | 0.71 |
| Breast_Mammary_Tissue | ENSG00000064703.7 | 166137 | 290 | 148 | 1.73E-02 | 0.14 | 0.06 | 0.71 |
| Artery_Coronary | ENSG00000143110.7 | 447590 | 173 | 81 | 1.74E-02 | -0.11 | 0.05 | 0.71 |

| | | | | | | | | |
|---|---|---|---|---|---|---|---|---|
| Minor_Salivary_Gland | ENSG00000156171.10 | 781166 | 97 | 49 | 1.74E-02 | -0.27 | 0.11 | 0.71 |
| Artery_Aorta | ENSG00000155363.14 | -751759 | 299 | 152 | 1.76E-02 | -0.09 | 0.04 | 0.71 |
| Liver | ENSG00000143110.7 | 447590 | 175 | 88 | 1.80E-02 | -0.16 | 0.07 | 0.71 |
| Prostate | ENSG00000155363.14 | -751759 | 152 | 75 | 1.87E-02 | -0.13 | 0.06 | 0.71 |
| Esophagus_Gastroesophageal_Junction | ENSG00000155366.12 | -786052 | 244 | 110 | 1.89E-02 | -0.12 | 0.05 | 0.71 |
| Brain_Spinal_cord_cervical_c-1 | ENSG00000236040.1 | 671128 | 91 | 49 | 2.09E-02 | 0.41 | 0.17 | 0.72 |
| Minor_Salivary_Gland | ENSG00000260948.1 | 488314 | 97 | 49 | 2.09E-02 | 0.24 | 0.10 | 0.72 |
| Brain_Hippocampus | ENSG00000184599.9 | -799037 | 123 | 63 | 2.10E-02 | -0.28 | 0.12 | 0.72 |
| Prostate | ENSG00000143079.10 | -474799 | 152 | 75 | 2.10E-02 | -0.15 | 0.06 | 0.72 |
| Brain_Caudate_basal_ganglia | ENSG00000184599.9 | -799037 | 160 | 88 | 2.19E-02 | -0.28 | 0.12 | 0.74 |
| Minor_Salivary_Gland | ENSG00000232811.1 | 977915 | 97 | 49 | 2.26E-02 | 0.34 | 0.15 | 0.75 |
| Colon_Transverse | ENSG00000134245.13 | -545159 | 274 | 119 | 2.34E-02 | -0.11 | 0.05 | 0.76 |
| Stomach | ENSG00000007341.14 | -699443 | 262 | 107 | 2.40E-02 | -0.19 | 0.08 | 0.76 |
| Minor_Salivary_Gland | ENSG00000261654.1 | 979029 | 97 | 49 | 2.49E-02 | 0.33 | 0.14 | 0.77 |
| Whole_Blood | ENSG00000261654.1 | 979029 | 407 | 195 | 2.53E-02 | 0.13 | 0.06 | 0.77 |
| Small_Intestine_Terminal_Ileum | ENSG00000155367.11 | -794095 | 137 | 54 | 2.54E-02 | -0.20 | 0.09 | 0.77 |
| Pituitary | ENSG00000231346.1 | 312659 | 183 | 95 | 2.73E-02 | -0.19 | 0.09 | 0.78 |
| Testis | ENSG00000261595.1 | -794290 | 259 | 128 | 2.73E-02 | 0.08 | 0.04 | 0.78 |
| Adrenal_Gland | ENSG00000143110.7 | 447590 | 190 | 96 | 2.74E-02 | -0.14 | 0.06 | 0.78 |
| Testis | ENSG00000197852.8 | 168128 | 259 | 128 | 2.87E-02 | -0.17 | 0.08 | 0.78 |
| Brain_Cerebellar_Hemisphere | ENSG00000243960.1 | 482557 | 136 | 74 | 2.88E-02 | -0.25 | 0.11 | 0.78 |
| Skin_Sun_Exposed_Lower_leg | ENSG00000261654.1 | 979029 | 473 | 233 | 2.97E-02 | 0.15 | 0.07 | 0.78 |
| Breast_Mammary_Tissue | ENSG00000162777.12 | 716847 | 290 | 148 | 2.97E-02 | 0.09 | 0.04 | 0.78 |
| Esophagus_Mucosa | ENSG00000243960.1 | 482557 | 407 | 208 | 2.99E-02 | 0.18 | 0.08 | 0.78 |
| Pituitary | ENSG00000007341.14 | -699443 | 183 | 95 | 3.03E-02 | -0.25 | 0.11 | 0.78 |
| Heart_Atrial_Appendage | ENSG00000155363.14 | -751759 | 297 | 147 | 3.10E-02 | -0.07 | 0.03 | 0.78 |
| Nerve_Tibial | ENSG00000155367.11 | -794095 | 414 | 202 | 3.10E-02 | 0.08 | 0.04 | 0.78 |
| Spleen | ENSG00000231346.1 | 312659 | 162 | 55 | 3.14E-02 | -0.28 | 0.13 | 0.78 |
| Small_Intestine_Terminal_Ileum | ENSG00000215867.4 | 271838 | 137 | 54 | 3.31E-02 | -0.34 | 0.16 | 0.81 |
| Esophagus_Mucosa | ENSG00000085465.11 | 493605 | 407 | 208 | 3.34E-02 | 0.08 | 0.04 | 0.81 |
| Pituitary | ENSG00000156171.10 | 781166 | 183 | 95 | 3.42E-02 | -0.20 | 0.09 | 0.81 |
| Brain_Nucleus_accumbens_basal_ganglia | ENSG00000134245.13 | -545159 | 147 | 77 | 3.49E-02 | -0.14 | 0.07 | 0.81 |
| Brain_Spinal_cord_cervical_c-1 | ENSG00000231437.3 | -68388 | 91 | 49 | 3.52E-02 | 0.21 | 0.10 | 0.81 |
| Brain_Caudate_basal_ganglia | ENSG00000121933.13 | 357420 | 160 | 88 | 3.64E-02 | 0.15 | 0.07 | 0.81 |
| Brain_Substantia_nigra | ENSG00000116459.6 | 472490 | 88 | 49 | 3.67E-02 | 0.15 | 0.07 | 0.81 |
| Brain_Frontal_Cortex_BA9 | ENSG00000134245.13 | -545159 | 129 | 73 | 3.67E-02 | -0.22 | 0.10 | 0.81 |
| Liver | ENSG00000116489.8 | -698415 | 175 | 88 | 3.76E-02 | -0.10 | 0.05 | 0.81 |
| Esophagus_Mucosa | ENSG00000184599.9 | -799037 | 407 | 208 | 3.79E-02 | -0.15 | 0.07 | 0.81 |
| Minor_Salivary_Gland | ENSG00000231346.1 | 312659 | 97 | 49 | 3.83E-02 | -0.25 | 0.12 | 0.81 |
| Adrenal_Gland | ENSG00000064703.7 | 166137 | 190 | 96 | 3.84E-02 | 0.15 | 0.07 | 0.81 |
| Esophagus_Mucosa | ENSG00000231437.3 | -68388 | 407 | 208 | 4.08E-02 | 0.11 | 0.06 | 0.84 |
| Cells_EBV-transformed_lymphocytes | ENSG00000007341.14 | -699443 | 130 | 68 | 4.10E-02 | -0.24 | 0.11 | 0.84 |
| Small_Intestine_Terminal_Ileum | ENSG00000273010.1 | 958080 | 137 | 54 | 4.32E-02 | 0.29 | 0.14 | 0.86 |
| Artery_Coronary | ENSG00000116455.9 | 472519 | 173 | 81 | 4.33E-02 | -0.09 | 0.04 | 0.86 |
| Adipose_Subcutaneous | ENSG00000116473.10 | 379164 | 442 | 214 | 4.35E-02 | -0.07 | 0.03 | 0.86 |
| Adrenal_Gland | ENSG00000155366.12 | -786052 | 190 | 96 | 4.44E-02 | 0.12 | 0.06 | 0.86 |
| Artery_Tibial | ENSG00000007341.14 | -699443 | 441 | 219 | 4.56E-02 | -0.12 | 0.06 | 0.86 |

| | | | | | | | | |
|---|---|---|---|---|---|---|---|---|
| Brain_Hippocampus | ENSG00000155367.11 | -794095 | 123 | 63 | 4.66E-02 | -0.15 | 0.08 | 0.86 |
| Heart_Atrial_Appendage | ENSG00000116489.8 | -698415 | 297 | 147 | 4.72E-02 | 0.06 | 0.03 | 0.86 |
| Brain_Hypothalamus | ENSG00000085465.11 | 493605 | 121 | 61 | 4.78E-02 | 0.14 | 0.07 | 0.86 |
| Testis | ENSG00000064703.7 | 166137 | 259 | 128 | 4.81E-02 | 0.05 | 0.03 | 0.86 |
| Cells_Transformed_fibroblasts | ENSG00000243960.1 | 482557 | 343 | 175 | 4.85E-02 | 0.14 | 0.07 | 0.86 |
| Thyroid | ENSG00000007341.14 | -699443 | 446 | 224 | 4.87E-02 | -0.11 | 0.06 | 0.86 |
| Cells_EBV-transformed_lymphocytes | ENSG00000197852.8 | 168128 | 130 | 68 | 4.97E-02 | 0.20 | 0.10 | 0.86 |
| Nerve_Tibial | ENSG00000232811.1 | 977915 | 414 | 202 | 5.07E-02 | -0.15 | 0.07 | 0.86 |
| Cells_EBV-transformed_lymphocytes | ENSG00000155363.14 | -751759 | 130 | 68 | 5.17E-02 | 0.16 | 0.08 | 0.86 |
| Brain_Cerebellum | ENSG00000231437.3 | -68388 | 173 | 96 | 5.18E-02 | 0.15 | 0.08 | 0.86 |
| Skin_Sun_Exposed_Lower_leg | ENSG00000116489.8 | -698415 | 473 | 233 | 5.29E-02 | -0.05 | 0.02 | 0.86 |
| Colon_Transverse | ENSG00000156171.10 | 781166 | 274 | 119 | 5.33E-02 | -0.10 | 0.05 | 0.86 |
| Nerve_Tibial | ENSG00000007341.14 | -699443 | 414 | 202 | 5.36E-02 | -0.12 | 0.06 | 0.86 |
| Cells_Transformed_fibroblasts | ENSG00000261654.1 | 979029 | 343 | 175 | 5.50E-02 | -0.10 | 0.05 | 0.86 |
| Esophagus_Gastroesophageal_Junction | ENSG00000064703.7 | 166137 | 244 | 110 | 5.62E-02 | 0.11 | 0.06 | 0.86 |
| Brain_Hippocampus | ENSG00000233337.1 | 483868 | 123 | 63 | 5.65E-02 | -0.22 | 0.12 | 0.86 |
| Esophagus_Muscularis | ENSG00000007341.14 | -699443 | 370 | 188 | 5.67E-02 | -0.13 | 0.07 | 0.86 |
| Breast_Mammary_Tissue | ENSG00000121933.13 | 357420 | 290 | 148 | 5.71E-02 | -0.10 | 0.05 | 0.86 |
| Stomach | ENSG00000121931.11 | 968421 | 262 | 107 | 5.75E-02 | -0.15 | 0.08 | 0.86 |
| Prostate | ENSG00000173947.9 | 575094 | 152 | 75 | 5.75E-02 | -0.12 | 0.06 | 0.86 |
| Pancreas | ENSG00000273483.1 | -597059 | 248 | 108 | 5.77E-02 | -0.18 | 0.09 | 0.86 |
| Muscle_Skeletal | ENSG00000273010.1 | 958080 | 564 | 276 | 5.82E-02 | 0.10 | 0.05 | 0.86 |
| Colon_Sigmoid | ENSG00000064703.7 | 166137 | 233 | 107 | 5.84E-02 | 0.10 | 0.05 | 0.86 |
| Brain_Nucleus_accumbens_basal_ganglia | ENSG00000116489.8 | -698415 | 147 | 77 | 5.90E-02 | -0.13 | 0.07 | 0.86 |
| Stomach | ENSG00000143110.7 | 447590 | 262 | 107 | 5.92E-02 | -0.10 | 0.05 | 0.86 |
| Cells_Transformed_fibroblasts | ENSG00000155366.12 | -786052 | 343 | 175 | 5.95E-02 | -0.04 | 0.02 | 0.86 |
| Brain_Hypothalamus | ENSG00000121933.13 | 357420 | 121 | 61 | 5.99E-02 | 0.11 | 0.06 | 0.86 |
| Breast_Mammary_Tissue | ENSG00000233337.1 | 483868 | 290 | 148 | 6.05E-02 | 0.15 | 0.08 | 0.86 |
| Brain_Nucleus_accumbens_basal_ganglia | ENSG00000224167.1 | -955558 | 147 | 77 | 6.09E-02 | 0.25 | 0.13 | 0.86 |
| Colon_Sigmoid | ENSG00000231246.1 | -439146 | 233 | 107 | 6.11E-02 | 0.18 | 0.10 | 0.86 |
| Spleen | ENSG00000232811.1 | 977915 | 162 | 55 | 6.19E-02 | 0.26 | 0.14 | 0.86 |
| Liver | ENSG00000116455.9 | 472519 | 175 | 88 | 6.34E-02 | -0.10 | 0.06 | 0.86 |
| Adipose_Visceral_Omentum | ENSG00000155363.14 | -751759 | 355 | 172 | 6.38E-02 | -0.06 | 0.03 | 0.86 |
| Cells_Transformed_fibroblasts | ENSG00000007341.14 | -699443 | 343 | 175 | 6.42E-02 | -0.14 | 0.07 | 0.86 |
| Brain_Cerebellum | ENSG00000116455.9 | 472519 | 173 | 96 | 6.48E-02 | 0.13 | 0.07 | 0.86 |
| Colon_Transverse | ENSG00000215866.3 | -929261 | 274 | 119 | 6.59E-02 | 0.15 | 0.08 | 0.86 |
| Breast_Mammary_Tissue | ENSG00000243960.1 | 482557 | 290 | 148 | 6.74E-02 | 0.17 | 0.09 | 0.86 |
| Skin_Sun_Exposed_Lower_leg | ENSG00000171385.5 | -67773 | 473 | 233 | 6.76E-02 | 0.07 | 0.04 | 0.86 |
| Esophagus_Gastroesophageal_Junction | ENSG00000155363.14 | -751759 | 244 | 110 | 6.79E-02 | -0.09 | 0.05 | 0.86 |
| Pancreas | ENSG00000143110.7 | 447590 | 248 | 108 | 6.79E-02 | -0.11 | 0.06 | 0.86 |
| Heart_Atrial_Appendage | ENSG00000197852.8 | 168128 | 297 | 147 | 6.82E-02 | 0.08 | 0.04 | 0.86 |
| Cells_EBV-transformed_lymphocytes | ENSG00000116489.8 | -698415 | 130 | 68 | 6.90E-02 | -0.10 | 0.05 | 0.86 |
| Brain_Nucleus_accumbens_basal_ganglia | ENSG00000227811.2 | 181541 | 147 | 77 | 6.92E-02 | -0.20 | 0.11 | 0.86 |
| Brain_Hippocampus | ENSG00000243960.1 | 482557 | 123 | 63 | 6.93E-02 | 0.29 | 0.16 | 0.86 |
| Skin_Sun_Exposed_Lower_leg | ENSG00000064703.7 | 166137 | 473 | 233 | 7.03E-02 | 0.08 | 0.04 | 0.86 |
| Brain_Caudate_basal_ganglia | ENSG00000231437.3 | -68388 | 160 | 88 | 7.05E-02 | 0.15 | 0.08 | 0.86 |

| | | | | | | | | |
|---|---|---|---|---|---|---|---|---|
| Colon_Sigmoid | ENSG00000116455.9 | 472519 | 233 | 107 | 7.07E-02 | -0.08 | 0.05 | 0.86 |
| Brain_Amygdala | ENSG00000184599.9 | -799037 | 100 | 47 | 7.09E-02 | 0.35 | 0.19 | 0.86 |
| Pancreas | ENSG00000233337.1 | 483868 | 248 | 108 | 7.09E-02 | 0.16 | 0.09 | 0.86 |
| Pancreas | ENSG00000173947.9 | 575094 | 248 | 108 | 7.10E-02 | 0.14 | 0.08 | 0.86 |
| Skin_Sun_Exposed_Lower_leg | ENSG00000143079.10 | -474799 | 473 | 233 | 7.12E-02 | 0.04 | 0.02 | 0.86 |
| Spleen | ENSG00000238975.1 | -731206 | 162 | 55 | 7.25E-02 | -0.23 | 0.12 | 0.86 |
| Whole_Blood | ENSG00000156171.10 | 781166 | 407 | 195 | 7.27E-02 | -0.06 | 0.03 | 0.86 |
| Small_Intestine_Terminal_Ileum | ENSG00000116473.10 | 379164 | 137 | 54 | 7.29E-02 | -0.15 | 0.08 | 0.86 |
| Uterus | ENSG00000273010.1 | 958080 | 111 | 63 | 7.31E-02 | 0.26 | 0.14 | 0.86 |
| Prostate | ENSG00000232811.1 | 977915 | 152 | 75 | 7.45E-02 | -0.22 | 0.12 | 0.86 |
| Brain_Putamen_basal_ganglia | ENSG00000231346.1 | 312659 | 124 | 71 | 7.48E-02 | -0.25 | 0.14 | 0.86 |
| Vagina | ENSG00000116459.6 | 472490 | 115 | 59 | 7.61E-02 | 0.06 | 0.04 | 0.86 |
| Testis | ENSG00000116455.9 | 472519 | 259 | 128 | 7.62E-02 | 0.10 | 0.06 | 0.86 |
| Pancreas | ENSG00000116455.9 | 472519 | 248 | 108 | 7.72E-02 | 0.12 | 0.07 | 0.86 |
| Adrenal_Gland | ENSG00000143079.10 | -474799 | 190 | 96 | 7.75E-02 | 0.13 | 0.07 | 0.86 |
| Cells_EBV-transformed_lymphocytes | ENSG00000261654.1 | 979029 | 130 | 68 | 7.75E-02 | 0.18 | 0.10 | 0.86 |
| Prostate | ENSG00000261654.1 | 979029 | 152 | 75 | 7.82E-02 | -0.21 | 0.12 | 0.86 |
| Muscle_Skeletal | ENSG00000116455.9 | 472519 | 564 | 276 | 7.83E-02 | 0.05 | 0.03 | 0.86 |
| Heart_Left_Ventricle | ENSG00000227811.2 | 181541 | 303 | 154 | 7.85E-02 | 0.12 | 0.07 | 0.86 |
| Skin_Not_Sun_Exposed_Suprapubic | ENSG00000155366.12 | -786052 | 387 | 200 | 7.92E-02 | -0.06 | 0.03 | 0.86 |
| Cells_Transformed_fibroblasts | ENSG00000184599.9 | -799037 | 343 | 175 | 7.99E-02 | -0.13 | 0.07 | 0.86 |
| Cells_Transformed_fibroblasts | ENSG00000116473.10 | 379164 | 343 | 175 | 8.13E-02 | -0.05 | 0.03 | 0.86 |
| Thyroid | ENSG00000273010.1 | 958080 | 446 | 224 | 8.15E-02 | -0.13 | 0.07 | 0.86 |
| Colon_Sigmoid | ENSG00000121931.11 | 968421 | 233 | 107 | 8.17E-02 | 0.14 | 0.08 | 0.86 |
| Adipose_Visceral_Omentum | ENSG00000116473.10 | 379164 | 355 | 172 | 8.24E-02 | -0.05 | 0.03 | 0.86 |
| Pancreas | ENSG00000134245.13 | -545159 | 248 | 108 | 8.38E-02 | 0.12 | 0.07 | 0.86 |
| Ovary | ENSG00000116455.9 | 472519 | 133 | 71 | 8.41E-02 | 0.11 | 0.06 | 0.86 |
| Thyroid | ENSG00000116455.9 | 472519 | 446 | 224 | 8.45E-02 | 0.05 | 0.03 | 0.86 |
| Breast_Mammary_Tissue | ENSG00000155366.12 | -786052 | 290 | 148 | 8.49E-02 | -0.06 | 0.03 | 0.86 |
| Thyroid | ENSG00000116473.10 | 379164 | 446 | 224 | 8.49E-02 | -0.05 | 0.03 | 0.86 |
| Minor_Salivary_Gland | ENSG00000007341.14 | -699443 | 97 | 49 | 8.50E-02 | -0.28 | 0.16 | 0.86 |
| Lung | ENSG00000238975.1 | -731206 | 427 | 222 | 8.65E-02 | 0.09 | 0.05 | 0.86 |
| Spleen | ENSG00000116459.6 | 472490 | 162 | 55 | 8.73E-02 | 0.08 | 0.05 | 0.86 |
| Pituitary | ENSG00000171385.5 | -67773 | 183 | 95 | 8.78E-02 | 0.15 | 0.09 | 0.86 |
| Brain_Cerebellar_Hemisphere | ENSG00000116459.6 | 472490 | 136 | 74 | 8.85E-02 | -0.08 | 0.05 | 0.86 |
| Skin_Sun_Exposed_Lower_leg | ENSG00000064886.9 | 720611 | 473 | 233 | 8.88E-02 | 0.07 | 0.04 | 0.86 |
| Brain_Amygdala | ENSG00000134255.9 | 781156 | 100 | 47 | 8.93E-02 | 0.19 | 0.11 | 0.86 |
| Esophagus_Muscularis | ENSG00000225075.1 | -775239 | 370 | 188 | 8.98E-02 | 0.13 | 0.08 | 0.86 |
| Brain_Cerebellar_Hemisphere | ENSG00000227811.2 | 181541 | 136 | 74 | 8.99E-02 | 0.23 | 0.13 | 0.86 |
| Esophagus_Gastroesophageal_Junction | ENSG00000116489.8 | -698415 | 244 | 110 | 9.09E-02 | -0.08 | 0.05 | 0.86 |
| Brain_Substantia_nigra | ENSG00000155366.12 | -786052 | 88 | 49 | 9.18E-02 | 0.10 | 0.06 | 0.86 |
| Colon_Transverse | ENSG00000134255.9 | 781156 | 274 | 119 | 9.20E-02 | -0.06 | 0.04 | 0.86 |
| Pituitary | ENSG00000143110.7 | 447590 | 183 | 95 | 9.21E-02 | 0.11 | 0.07 | 0.86 |
| Pituitary | ENSG00000143079.10 | -474799 | 183 | 95 | 9.23E-02 | 0.12 | 0.07 | 0.86 |
| Liver | ENSG00000116473.10 | 379164 | 175 | 88 | 9.26E-02 | -0.13 | 0.08 | 0.86 |
| Brain_Substantia_nigra | ENSG00000162777.12 | 716847 | 88 | 49 | 9.27E-02 | 0.16 | 0.09 | 0.86 |
| Testis | ENSG00000155363.14 | -751759 | 259 | 128 | 9.43E-02 | 0.04 | 0.03 | 0.86 |

| | | | | | | | | |
|---|---|---|---|---|---|---|---|---|
| Brain_Caudate_basal_ganglia | ENSG00000171385.5 | -67773 | 160 | 88 | 9.58E-02 | 0.13 | 0.08 | 0.86 |
| Uterus | ENSG00000260948.1 | 488314 | 111 | 63 | 9.59E-02 | -0.18 | 0.11 | 0.86 |
| Adipose_Visceral_Omentum | ENSG00000155367.11 | -794095 | 355 | 172 | 9.59E-02 | -0.08 | 0.05 | 0.86 |
| Esophagus_Mucosa | ENSG00000134245.13 | -545159 | 407 | 208 | 9.63E-02 | -0.06 | 0.04 | 0.86 |
| Brain_Substantia_nigra | ENSG00000243960.1 | 482557 | 88 | 49 | 9.65E-02 | 0.25 | 0.15 | 0.86 |
| Adipose_Visceral_Omentum | ENSG00000227811.2 | 181541 | 355 | 172 | 9.71E-02 | -0.13 | 0.08 | 0.86 |
| Brain_Putamen_basal_ganglia | ENSG00000231437.3 | -68388 | 124 | 71 | 9.77E-02 | 0.15 | 0.09 | 0.86 |
| Ovary | ENSG00000134245.13 | -545159 | 133 | 71 | 9.79E-02 | 0.14 | 0.08 | 0.86 |
| Brain_Hypothalamus | ENSG00000231346.1 | 312659 | 121 | 61 | 9.94E-02 | 0.19 | 0.12 | 0.86 |
| Testis | ENSG00000231246.1 | -439146 | 259 | 128 | 9.95E-02 | 0.11 | 0.07 | 0.86 |
| Brain_Cortex | ENSG00000116459.6 | 472490 | 158 | 79 | 9.99E-02 | -0.12 | 0.07 | 0.86 |
| Spleen | ENSG00000116473.10 | 379164 | 162 | 55 | 1.00E-01 | -0.13 | 0.08 | 0.86 |
| Heart_Left_Ventricle | ENSG00000085465.11 | 493605 | 303 | 154 | 1.00E-01 | -0.08 | 0.05 | 0.86 |
| Stomach | ENSG00000238975.1 | -731206 | 262 | 107 | 1.01E-01 | 0.15 | 0.09 | 0.86 |
| Esophagus_Mucosa | ENSG00000225075.1 | -775239 | 407 | 208 | 1.01E-01 | -0.12 | 0.07 | 0.86 |
| Brain_Amygdala | ENSG00000231437.3 | -68388 | 100 | 47 | 1.02E-01 | 0.23 | 0.14 | 0.86 |
| Brain_Cortex | ENSG00000197852.8 | 168128 | 158 | 79 | 1.02E-01 | -0.09 | 0.05 | 0.86 |
| Brain_Frontal_Cortex_BA9 | ENSG00000233337.1 | 483868 | 129 | 73 | 1.03E-01 | 0.23 | 0.14 | 0.86 |
| Thyroid | ENSG00000215866.3 | -929261 | 446 | 224 | 1.03E-01 | 0.11 | 0.07 | 0.86 |
| Nerve_Tibial | ENSG00000121931.11 | 968421 | 414 | 202 | 1.04E-01 | -0.06 | 0.04 | 0.86 |
| Artery_Coronary | ENSG00000143079.10 | -474799 | 173 | 81 | 1.04E-01 | 0.12 | 0.07 | 0.86 |
| Esophagus_Mucosa | ENSG00000143079.10 | -474799 | 407 | 208 | 1.05E-01 | 0.04 | 0.03 | 0.86 |
| Small_Intestine_Terminal_Ileum | ENSG00000173947.9 | 575094 | 137 | 54 | 1.07E-01 | -0.13 | 0.08 | 0.86 |
| Pituitary | ENSG00000162777.12 | 716847 | 183 | 95 | 1.07E-01 | 0.13 | 0.08 | 0.86 |
| Pituitary | ENSG00000116455.9 | 472519 | 183 | 95 | 1.07E-01 | -0.10 | 0.06 | 0.86 |
| Brain_Spinal_cord_cervical_c-1 | ENSG00000171385.5 | -67773 | 91 | 49 | 1.07E-01 | 0.17 | 0.11 | 0.86 |
| Testis | ENSG00000116459.6 | 472490 | 259 | 128 | 1.07E-01 | 0.08 | 0.05 | 0.86 |
| Skin_Sun_Exposed_Lower_leg | ENSG00000273010.1 | 958080 | 473 | 233 | 1.08E-01 | -0.12 | 0.07 | 0.86 |
| Small_Intestine_Terminal_Ileum | ENSG00000156171.10 | 781166 | 137 | 54 | 1.08E-01 | 0.10 | 0.06 | 0.86 |
| Minor_Salivary_Gland | ENSG00000064886.9 | 720611 | 97 | 49 | 1.08E-01 | -0.11 | 0.07 | 0.86 |
| Uterus | ENSG00000064703.7 | 166137 | 111 | 63 | 1.09E-01 | 0.15 | 0.09 | 0.86 |
| Heart_Atrial_Appendage | ENSG00000233337.1 | 483868 | 297 | 147 | 1.09E-01 | -0.13 | 0.08 | 0.86 |
| Artery_Tibial | ENSG00000232811.1 | 977915 | 441 | 219 | 1.10E-01 | -0.11 | 0.07 | 0.86 |
| Lung | ENSG00000232811.1 | 977915 | 427 | 222 | 1.10E-01 | 0.11 | 0.07 | 0.86 |
| Brain_Nucleus_accumbens_basal_ganglia | ENSG00000273483.1 | -597059 | 147 | 77 | 1.10E-01 | 0.12 | 0.08 | 0.86 |
| Minor_Salivary_Gland | ENSG00000273483.1 | -597059 | 97 | 49 | 1.11E-01 | 0.22 | 0.14 | 0.86 |
| Esophagus_Muscularis | ENSG00000155363.14 | -751759 | 370 | 188 | 1.12E-01 | -0.05 | 0.03 | 0.86 |
| Cells_Transformed_fibroblasts | ENSG00000225075.1 | -775239 | 343 | 175 | 1.12E-01 | -0.13 | 0.08 | 0.86 |
| Pancreas | ENSG00000121933.13 | 357420 | 248 | 108 | 1.12E-01 | -0.08 | 0.05 | 0.86 |
| Heart_Atrial_Appendage | ENSG00000243960.1 | 482557 | 297 | 147 | 1.14E-01 | 0.14 | 0.09 | 0.87 |
| Brain_Anterior_cingulate_cortex_BA24 | ENSG00000085465.11 | 493605 | 121 | 65 | 1.14E-01 | 0.11 | 0.07 | 0.87 |
| Pituitary | ENSG00000197852.8 | 168128 | 183 | 95 | 1.15E-01 | -0.17 | 0.11 | 0.87 |
| Pituitary | ENSG00000155367.11 | -794095 | 183 | 95 | 1.16E-01 | 0.15 | 0.10 | 0.88 |
| Brain_Cerebellar_Hemisphere | ENSG00000155367.11 | -794095 | 136 | 74 | 1.17E-01 | -0.13 | 0.08 | 0.88 |
| Ovary | ENSG00000232811.1 | 977915 | 133 | 71 | 1.17E-01 | -0.20 | 0.12 | 0.88 |
| Thyroid | ENSG00000064703.7 | 166137 | 446 | 224 | 1.18E-01 | -0.04 | 0.03 | 0.88 |
| Prostate | ENSG00000273010.1 | 958080 | 152 | 75 | 1.19E-01 | -0.19 | 0.12 | 0.88 |

| | | | | | | | | |
|---|---|---|---|---|---|---|---|---|
| Brain_Cerebellum | ENSG00000215866.3 | -929261 | 173 | 96 | 1.21E-01 | 0.18 | 0.11 | 0.88 |
| Artery_Aorta | ENSG00000116489.8 | -698415 | 299 | 152 | 1.22E-01 | 0.05 | 0.03 | 0.88 |
| Brain_Amygdala | ENSG00000231346.1 | 312659 | 100 | 47 | 1.22E-01 | 0.22 | 0.14 | 0.88 |
| Brain_Spinal_cord_cervical_c-1 | ENSG00000134255.9 | 781156 | 91 | 49 | 1.22E-01 | 0.12 | 0.08 | 0.88 |
| Spleen | ENSG00000121931.11 | 968421 | 162 | 55 | 1.25E-01 | -0.17 | 0.11 | 0.88 |
| Brain_Hippocampus | ENSG00000121933.13 | 357420 | 123 | 63 | 1.25E-01 | 0.08 | 0.05 | 0.88 |
| Brain_Cerebellum | ENSG00000215867.4 | 271838 | 173 | 96 | 1.25E-01 | -0.18 | 0.12 | 0.88 |
| Esophagus_Muscularis | ENSG00000171385.5 | -67773 | 370 | 188 | 1.25E-01 | -0.04 | 0.03 | 0.88 |
| Spleen | ENSG00000243960.1 | 482557 | 162 | 55 | 1.25E-01 | -0.23 | 0.15 | 0.88 |
| Brain_Putamen_basal_ganglia | ENSG00000232811.1 | 977915 | 124 | 71 | 1.26E-01 | 0.21 | 0.14 | 0.88 |
| Testis | ENSG00000134245.13 | -545159 | 259 | 128 | 1.27E-01 | 0.07 | 0.04 | 0.88 |
| Nerve_Tibial | ENSG00000143079.10 | -474799 | 414 | 202 | 1.27E-01 | -0.05 | 0.03 | 0.88 |
| Thyroid | ENSG00000064886.9 | 720611 | 446 | 224 | 1.28E-01 | 0.08 | 0.05 | 0.88 |
| Brain_Substantia_nigra | ENSG00000227811.2 | 181541 | 88 | 49 | 1.30E-01 | 0.28 | 0.18 | 0.88 |
| Brain_Hippocampus | ENSG00000173947.9 | 575094 | 123 | 63 | 1.30E-01 | 0.10 | 0.07 | 0.88 |
| Breast_Mammary_Tissue | ENSG00000227811.2 | 181541 | 290 | 148 | 1.30E-01 | 0.13 | 0.09 | 0.88 |
| Esophagus_Mucosa | ENSG00000156171.10 | 781166 | 407 | 208 | 1.30E-01 | -0.08 | 0.06 | 0.88 |
| Nerve_Tibial | ENSG00000121933.13 | 357420 | 414 | 202 | 1.32E-01 | -0.06 | 0.04 | 0.88 |
| Brain_Hypothalamus | ENSG00000231437.3 | -68388 | 121 | 61 | 1.32E-01 | 0.15 | 0.10 | 0.88 |
| Ovary | ENSG00000171385.5 | -67773 | 133 | 71 | 1.33E-01 | -0.14 | 0.09 | 0.88 |
| Lung | ENSG00000121931.11 | 968421 | 427 | 222 | 1.33E-01 | -0.04 | 0.03 | 0.88 |
| Adrenal_Gland | ENSG00000134245.13 | -545159 | 190 | 96 | 1.33E-01 | 0.11 | 0.07 | 0.88 |
| Artery_Aorta | ENSG00000143079.10 | -474799 | 299 | 152 | 1.33E-01 | 0.07 | 0.04 | 0.88 |
| Esophagus_Gastroesophageal_Junction | ENSG00000261654.1 | 979029 | 244 | 110 | 1.34E-01 | -0.14 | 0.09 | 0.88 |
| Colon_Transverse | ENSG00000007341.14 | -699443 | 274 | 119 | 1.34E-01 | -0.13 | 0.09 | 0.88 |
| Artery_Tibial | ENSG00000155367.11 | -794095 | 441 | 219 | 1.34E-01 | 0.06 | 0.04 | 0.88 |
| Brain_Cortex | ENSG00000231437.3 | -68388 | 158 | 79 | 1.34E-01 | 0.18 | 0.12 | 0.88 |
| Thyroid | ENSG00000238975.1 | -731206 | 446 | 224 | 1.35E-01 | 0.09 | 0.06 | 0.88 |
| Vagina | ENSG00000116455.9 | 472519 | 115 | 59 | 1.36E-01 | 0.11 | 0.07 | 0.88 |
| Prostate | ENSG00000116473.10 | 379164 | 152 | 75 | 1.37E-01 | -0.07 | 0.05 | 0.88 |
| Artery_Coronary | ENSG00000156171.10 | 781166 | 173 | 81 | 1.38E-01 | -0.10 | 0.07 | 0.88 |
| Brain_Substantia_nigra | ENSG00000173947.9 | 575094 | 88 | 49 | 1.38E-01 | 0.18 | 0.12 | 0.88 |
| Adipose_Subcutaneous | ENSG00000171385.5 | -67773 | 442 | 214 | 1.39E-01 | -0.06 | 0.04 | 0.88 |
| Pancreas | ENSG00000143079.10 | -474799 | 248 | 108 | 1.41E-01 | 0.09 | 0.06 | 0.88 |
| Brain_Substantia_nigra | ENSG00000134255.9 | 781156 | 88 | 49 | 1.41E-01 | 0.10 | 0.07 | 0.88 |
| Stomach | ENSG00000229283.1 | 603455 | 262 | 107 | 1.42E-01 | 0.10 | 0.07 | 0.88 |
| Adipose_Subcutaneous | ENSG00000162777.12 | 716847 | 442 | 214 | 1.42E-01 | 0.05 | 0.03 | 0.88 |
| Testis | ENSG00000237556.1 | 12046 | 259 | 128 | 1.42E-01 | 0.07 | 0.05 | 0.88 |
| Brain_Substantia_nigra | ENSG00000116473.10 | 379164 | 88 | 49 | 1.43E-01 | 0.09 | 0.06 | 0.88 |
| Brain_Frontal_Cortex_BA9 | ENSG00000116473.10 | 379164 | 129 | 73 | 1.43E-01 | -0.08 | 0.05 | 0.88 |
| Nerve_Tibial | ENSG00000261654.1 | 979029 | 414 | 202 | 1.44E-01 | -0.09 | 0.06 | 0.88 |
| Testis | ENSG00000156171.10 | 781166 | 259 | 128 | 1.44E-01 | -0.06 | 0.04 | 0.88 |
| Skin_Sun_Exposed_Lower_leg | ENSG00000227811.2 | 181541 | 473 | 233 | 1.44E-01 | -0.11 | 0.08 | 0.88 |
| Brain_Caudate_basal_ganglia | ENSG00000143079.10 | -474799 | 160 | 88 | 1.45E-01 | 0.11 | 0.07 | 0.88 |
| Heart_Atrial_Appendage | ENSG00000227811.2 | 181541 | 297 | 147 | 1.45E-01 | 0.09 | 0.06 | 0.88 |
| Esophagus_Gastroesophageal_Junction | ENSG00000231346.1 | 312659 | 244 | 110 | 1.46E-01 | 0.08 | 0.06 | 0.88 |
| Small_Intestine_Terminal_Ileum | ENSG00000116459.6 | 472490 | 137 | 54 | 1.46E-01 | -0.06 | 0.04 | 0.88 |

| | | | | | | | | |
|---|---|---|---|---|---|---|---|---|
| Vagina | ENSG00000231437.3 | -68388 | 115 | 59 | 1.47E-01 | 0.16 | 0.11 | 0.88 |
| Brain_Cerebellar_Hemisphere | ENSG00000116489.8 | -698415 | 136 | 74 | 1.47E-01 | -0.14 | 0.10 | 0.88 |
| Brain_Cortex | ENSG00000261654.1 | 979029 | 158 | 79 | 1.47E-01 | 0.16 | 0.11 | 0.88 |
| Testis | ENSG00000273010.1 | 958080 | 259 | 128 | 1.48E-01 | -0.09 | 0.06 | 0.88 |
| Esophagus_Muscularis | ENSG00000116455.9 | 472519 | 370 | 188 | 1.49E-01 | 0.05 | 0.03 | 0.88 |
| Brain_Substantia_nigra | ENSG00000184599.9 | -799037 | 88 | 49 | 1.49E-01 | 0.14 | 0.09 | 0.88 |
| Brain_Spinal_cord_cervical_c-1 | ENSG00000064703.7 | 166137 | 91 | 49 | 1.50E-01 | 0.19 | 0.13 | 0.88 |
| Nerve_Tibial | ENSG00000116489.8 | -698415 | 414 | 202 | 1.50E-01 | -0.05 | 0.03 | 0.88 |
| Brain_Spinal_cord_cervical_c-1 | ENSG00000134245.13 | -545159 | 91 | 49 | 1.51E-01 | -0.20 | 0.14 | 0.88 |
| Brain_Anterior_cingulate_cortex_BA24 | ENSG00000232811.1 | 977915 | 121 | 65 | 1.51E-01 | -0.20 | 0.14 | 0.88 |
| Cells_EBV-transformed_lymphocytes | ENSG00000273010.1 | 958080 | 130 | 68 | 1.51E-01 | -0.20 | 0.14 | 0.88 |
| Brain_Nucleus_accumbens_basal_ganglia | ENSG00000231346.1 | 312659 | 147 | 77 | 1.52E-01 | 0.18 | 0.13 | 0.88 |
| Esophagus_Mucosa | ENSG00000007341.14 | -699443 | 407 | 208 | 1.53E-01 | -0.08 | 0.05 | 0.88 |
| Lung | ENSG00000215867.4 | 271838 | 427 | 222 | 1.53E-01 | -0.10 | 0.07 | 0.88 |
| Brain_Frontal_Cortex_BA9 | ENSG00000143110.7 | 447590 | 129 | 73 | 1.53E-01 | 0.11 | 0.08 | 0.88 |
| Artery_Aorta | ENSG00000225075.1 | -775239 | 299 | 152 | 1.55E-01 | 0.13 | 0.09 | 0.88 |
| Brain_Caudate_basal_ganglia | ENSG00000231346.1 | 312659 | 160 | 88 | 1.56E-01 | 0.15 | 0.11 | 0.88 |
| Uterus | ENSG00000227811.2 | 181541 | 111 | 63 | 1.56E-01 | -0.20 | 0.14 | 0.88 |
| Brain_Cortex | ENSG00000155367.11 | -794095 | 158 | 79 | 1.58E-01 | 0.13 | 0.09 | 0.88 |
| Heart_Atrial_Appendage | ENSG00000173947.9 | 575094 | 297 | 147 | 1.59E-01 | -0.05 | 0.03 | 0.88 |
| Brain_Hypothalamus | ENSG00000134255.9 | 781156 | 121 | 61 | 1.59E-01 | -0.08 | 0.05 | 0.88 |
| Cells_Transformed_fibroblasts | ENSG00000260948.1 | 488314 | 343 | 175 | 1.60E-01 | 0.05 | 0.04 | 0.88 |
| Brain_Caudate_basal_ganglia | ENSG00000121931.11 | 968421 | 160 | 88 | 1.60E-01 | 0.13 | 0.09 | 0.88 |
| Lung | ENSG00000085465.11 | 493605 | 427 | 222 | 1.61E-01 | 0.04 | 0.03 | 0.88 |
| Liver | ENSG00000121931.11 | 968421 | 175 | 88 | 1.61E-01 | 0.11 | 0.07 | 0.88 |
| Skin_Sun_Exposed_Lower_leg | ENSG00000273483.1 | -597059 | 473 | 233 | 1.61E-01 | -0.10 | 0.07 | 0.88 |
| Liver | ENSG00000231437.3 | -68388 | 175 | 88 | 1.61E-01 | -0.14 | 0.10 | 0.88 |
| Prostate | ENSG00000162777.12 | 716847 | 152 | 75 | 1.62E-01 | -0.08 | 0.05 | 0.88 |
| Ovary | ENSG00000064703.7 | 166137 | 133 | 71 | 1.62E-01 | 0.10 | 0.07 | 0.88 |
| Artery_Tibial | ENSG00000156171.10 | 781166 | 441 | 219 | 1.63E-01 | -0.05 | 0.04 | 0.88 |
| Skin_Sun_Exposed_Lower_leg | ENSG00000238975.1 | -731206 | 473 | 233 | 1.65E-01 | -0.08 | 0.06 | 0.88 |
| Minor_Salivary_Gland | ENSG00000233337.1 | 483868 | 97 | 49 | 1.65E-01 | 0.13 | 0.09 | 0.88 |
| Esophagus_Gastroesophageal_Junction | ENSG00000143079.10 | -474799 | 244 | 110 | 1.65E-01 | -0.10 | 0.07 | 0.88 |
| Cells_Transformed_fibroblasts | ENSG00000273010.1 | 958080 | 343 | 175 | 1.65E-01 | -0.13 | 0.09 | 0.88 |
| Artery_Tibial | ENSG00000227811.2 | 181541 | 441 | 219 | 1.66E-01 | 0.10 | 0.07 | 0.88 |
| Small_Intestine_Terminal_Ileum | ENSG00000121933.13 | 357420 | 137 | 54 | 1.66E-01 | 0.15 | 0.11 | 0.88 |
| Brain_Spinal_cord_cervical_c-1 | ENSG00000173947.9 | 575094 | 91 | 49 | 1.66E-01 | 0.20 | 0.14 | 0.88 |
| Small_Intestine_Terminal_Ileum | ENSG00000231437.3 | -68388 | 137 | 54 | 1.67E-01 | 0.13 | 0.09 | 0.88 |
| Breast_Mammary_Tissue | ENSG00000231346.1 | 312659 | 290 | 148 | 1.67E-01 | 0.10 | 0.07 | 0.88 |
| Minor_Salivary_Gland | ENSG00000121931.11 | 968421 | 97 | 49 | 1.68E-01 | 0.15 | 0.11 | 0.88 |
| Colon_Sigmoid | ENSG00000156171.10 | 781166 | 233 | 107 | 1.68E-01 | -0.10 | 0.07 | 0.88 |
| Spleen | ENSG00000273010.1 | 958080 | 162 | 55 | 1.68E-01 | -0.22 | 0.16 | 0.88 |
| Skin_Not_Sun_Exposed_Suprapubic | ENSG00000238975.1 | -731206 | 387 | 200 | 1.70E-01 | -0.08 | 0.06 | 0.88 |
| Brain_Frontal_Cortex_BA9 | ENSG00000064703.7 | 166137 | 129 | 73 | 1.70E-01 | -0.15 | 0.11 | 0.88 |
| Testis | ENSG00000243960.1 | 482557 | 259 | 128 | 1.70E-01 | -0.13 | 0.09 | 0.88 |
| Cells_EBV-transformed_lymphocytes | ENSG00000143110.7 | 447590 | 130 | 68 | 1.71E-01 | 0.13 | 0.09 | 0.88 |
| Ovary | ENSG00000231437.3 | -68388 | 133 | 71 | 1.72E-01 | -0.15 | 0.11 | 0.88 |

| | | | | | | | | |
|---|---|---|---|---|---|---|---|---|
| Brain_Putamen_basal_ganglia | ENSG00000007341.14 | -699443 | 124 | 71 | 1.72E-01 | -0.19 | 0.14 | 0.88 |
| Brain_Hypothalamus | ENSG00000162777.12 | 716847 | 121 | 61 | 1.72E-01 | 0.14 | 0.10 | 0.88 |
| Cells_Transformed_fibroblasts | ENSG00000143079.10 | -474799 | 343 | 175 | 1.73E-01 | -0.05 | 0.03 | 0.88 |
| Breast_Mammary_Tissue | ENSG00000007341.14 | -699443 | 290 | 148 | 1.73E-01 | -0.10 | 0.07 | 0.88 |
| Artery_Tibial | ENSG00000231346.1 | 312659 | 441 | 219 | 1.73E-01 | 0.05 | 0.03 | 0.88 |
| Liver | ENSG00000171385.5 | -67773 | 175 | 88 | 1.74E-01 | -0.10 | 0.07 | 0.88 |
| Minor_Salivary_Gland | ENSG00000171385.5 | -67773 | 97 | 49 | 1.74E-01 | -0.13 | 0.09 | 0.88 |
| Pancreas | ENSG00000197852.8 | 168128 | 248 | 108 | 1.74E-01 | 0.11 | 0.08 | 0.88 |
| Cells_Transformed_fibroblasts | ENSG00000215866.3 | -929261 | 343 | 175 | 1.75E-01 | 0.10 | 0.07 | 0.88 |
| Brain_Frontal_Cortex_BA9 | ENSG00000224167.1 | -955558 | 129 | 73 | 1.75E-01 | -0.19 | 0.14 | 0.88 |
| Esophagus_Gastroesophageal_Junction | ENSG00000134245.13 | -545159 | 244 | 110 | 1.76E-01 | -0.09 | 0.06 | 0.88 |
| Brain_Putamen_basal_ganglia | ENSG00000227811.2 | 181541 | 124 | 71 | 1.76E-01 | -0.20 | 0.15 | 0.88 |
| Brain_Hypothalamus | ENSG00000116455.9 | 472519 | 121 | 61 | 1.76E-01 | -0.13 | 0.09 | 0.88 |
| Brain_Hippocampus | ENSG00000064703.7 | 166137 | 123 | 63 | 1.77E-01 | -0.15 | 0.11 | 0.88 |
| Lung | ENSG00000156171.10 | 781166 | 427 | 222 | 1.78E-01 | -0.06 | 0.04 | 0.88 |
| Brain_Nucleus_accumbens_basal_ganglia | ENSG00000243960.1 | 482557 | 147 | 77 | 1.78E-01 | 0.19 | 0.14 | 0.88 |
| Brain_Hippocampus | ENSG00000231437.3 | -68388 | 123 | 63 | 1.78E-01 | 0.11 | 0.08 | 0.88 |
| Prostate | ENSG00000143110.7 | 447590 | 152 | 75 | 1.78E-01 | -0.11 | 0.08 | 0.88 |
| Uterus | ENSG00000134255.9 | 781156 | 111 | 63 | 1.79E-01 | -0.13 | 0.09 | 0.88 |
| Brain_Putamen_basal_ganglia | ENSG00000162777.12 | 716847 | 124 | 71 | 1.80E-01 | -0.15 | 0.11 | 0.88 |
| Brain_Cortex | ENSG00000121933.13 | 357420 | 158 | 79 | 1.80E-01 | 0.12 | 0.09 | 0.88 |
| Brain_Hypothalamus | ENSG00000243960.1 | 482557 | 121 | 61 | 1.80E-01 | -0.18 | 0.14 | 0.88 |
| Brain_Caudate_basal_ganglia | ENSG00000064886.9 | 720611 | 160 | 88 | 1.80E-01 | -0.14 | 0.10 | 0.88 |
| Esophagus_Muscularis | ENSG00000156171.10 | 781166 | 370 | 188 | 1.81E-01 | -0.08 | 0.06 | 0.89 |
| Lung | ENSG00000121933.13 | 357420 | 427 | 222 | 1.83E-01 | 0.04 | 0.03 | 0.89 |
| Adipose_Subcutaneous | ENSG00000134245.13 | -545159 | 442 | 214 | 1.84E-01 | -0.06 | 0.05 | 0.89 |
| Skin_Not_Sun_Exposed_Suprapubic | ENSG00000225075.1 | -775239 | 387 | 200 | 1.85E-01 | 0.09 | 0.07 | 0.89 |
| Adrenal_Gland | ENSG00000197852.8 | 168128 | 190 | 96 | 1.85E-01 | 0.12 | 0.09 | 0.89 |
| Artery_Coronary | ENSG00000233337.1 | 483868 | 173 | 81 | 1.85E-01 | -0.15 | 0.11 | 0.89 |
| Adipose_Visceral_Omentum | ENSG00000171385.5 | -67773 | 355 | 172 | 1.85E-01 | 0.08 | 0.06 | 0.89 |
| Esophagus_Gastroesophageal_Junction | ENSG00000233337.1 | 483868 | 244 | 110 | 1.87E-01 | -0.12 | 0.09 | 0.89 |
| Brain_Nucleus_accumbens_basal_ganglia | ENSG00000121931.11 | 968421 | 147 | 77 | 1.88E-01 | 0.11 | 0.09 | 0.89 |
| Artery_Coronary | ENSG00000261654.1 | 979029 | 173 | 81 | 1.89E-01 | -0.15 | 0.11 | 0.89 |
| Brain_Hippocampus | ENSG00000134245.13 | -545159 | 123 | 63 | 1.89E-01 | 0.15 | 0.11 | 0.89 |
| Brain_Amygdala | ENSG00000143079.10 | -474799 | 100 | 47 | 1.89E-01 | 0.10 | 0.08 | 0.89 |
| Small_Intestine_Terminal_Ileum | ENSG00000243960.1 | 482557 | 137 | 54 | 1.90E-01 | -0.16 | 0.12 | 0.89 |
| Brain_Substantia_nigra | ENSG00000232811.1 | 977915 | 88 | 49 | 1.91E-01 | -0.22 | 0.17 | 0.89 |
| Brain_Frontal_Cortex_BA9 | ENSG00000116459.6 | 472490 | 129 | 73 | 1.92E-01 | -0.11 | 0.08 | 0.89 |
| Artery_Tibial | ENSG00000155363.14 | -751759 | 441 | 219 | 1.93E-01 | -0.04 | 0.03 | 0.89 |
| Esophagus_Gastroesophageal_Junction | ENSG00000162777.12 | 716847 | 244 | 110 | 1.94E-01 | -0.08 | 0.06 | 0.89 |
| Adipose_Visceral_Omentum | ENSG00000231437.3 | -68388 | 355 | 172 | 1.94E-01 | 0.07 | 0.05 | 0.89 |
| Brain_Anterior_cingulate_cortex_BA24 | ENSG00000233337.1 | 483868 | 121 | 65 | 1.94E-01 | 0.18 | 0.14 | 0.89 |
| Muscle_Skeletal | ENSG00000231346.1 | 312659 | 564 | 276 | 1.94E-01 | -0.08 | 0.06 | 0.89 |
| Brain_Cerebellar_Hemisphere | ENSG00000260948.1 | 488314 | 136 | 74 | 1.94E-01 | -0.12 | 0.09 | 0.89 |
| Brain_Spinal_cord_cervical_c-1 | ENSG00000116459.6 | 472490 | 91 | 49 | 1.95E-01 | -0.12 | 0.09 | 0.89 |
| Minor_Salivary_Gland | ENSG00000085465.11 | 493605 | 97 | 49 | 1.97E-01 | 0.12 | 0.09 | 0.89 |

| | | | | | | | | |
|---|---|---|---|---|---|---|---|---|
| Skin_Not_Sun_Exposed_Suprapubic | ENSG00000197852.8 | 168128 | 387 | 200 | 1.97E-01 | 0.05 | 0.04 | 0.89 |
| Nerve_Tibial | ENSG00000143110.7 | 447590 | 414 | 202 | 1.97E-01 | -0.04 | 0.03 | 0.89 |
| Nerve_Tibial | ENSG00000273483.1 | -597059 | 414 | 202 | 1.98E-01 | 0.07 | 0.06 | 0.89 |
| Uterus | ENSG00000273483.1 | -597059 | 111 | 63 | 1.99E-01 | 0.13 | 0.10 | 0.89 |
| Pituitary | ENSG00000173947.9 | 575094 | 183 | 95 | 1.99E-01 | -0.08 | 0.07 | 0.89 |
| Uterus | ENSG00000155363.14 | -751759 | 111 | 63 | 1.99E-01 | -0.09 | 0.07 | 0.89 |
| Whole_Blood | ENSG00000007341.14 | -699443 | 407 | 195 | 2.01E-01 | -0.07 | 0.05 | 0.89 |
| Lung | ENSG00000134255.9 | 781156 | 427 | 222 | 2.01E-01 | -0.05 | 0.04 | 0.89 |
| Heart_Atrial_Appendage | ENSG00000162777.12 | 716847 | 297 | 147 | 2.01E-01 | 0.07 | 0.05 | 0.89 |
| Brain_Cortex | ENSG00000134245.13 | -545159 | 158 | 79 | 2.01E-01 | -0.15 | 0.12 | 0.89 |
| Esophagus_Mucosa | ENSG00000064886.9 | 720611 | 407 | 208 | 2.02E-01 | 0.06 | 0.05 | 0.89 |
| Skin_Not_Sun_Exposed_Suprapubic | ENSG00000116473.10 | 379164 | 387 | 200 | 2.02E-01 | -0.04 | 0.03 | 0.89 |
| Brain_Anterior_cingulate_cortex_BA24 | ENSG00000197852.8 | 168128 | 121 | 65 | 2.02E-01 | -0.08 | 0.06 | 0.89 |
| Brain_Cortex | ENSG00000273483.1 | -597059 | 158 | 79 | 2.02E-01 | 0.11 | 0.08 | 0.89 |
| Uterus | ENSG00000261654.1 | 979029 | 111 | 63 | 2.02E-01 | 0.15 | 0.12 | 0.89 |
| Brain_Frontal_Cortex_BA9 | ENSG00000134255.9 | 781156 | 129 | 73 | 2.03E-01 | 0.08 | 0.06 | 0.89 |
| Artery_Tibial | ENSG00000260948.1 | 488314 | 441 | 219 | 2.03E-01 | -0.07 | 0.05 | 0.89 |
| Brain_Spinal_cord_cervical_c-1 | ENSG00000121931.11 | 968421 | 91 | 49 | 2.04E-01 | 0.21 | 0.17 | 0.89 |
| Pancreas | ENSG00000155367.11 | -794095 | 248 | 108 | 2.04E-01 | -0.08 | 0.07 | 0.89 |
| Breast_Mammary_Tissue | ENSG00000231437.3 | -68388 | 290 | 148 | 2.06E-01 | 0.09 | 0.07 | 0.89 |
| Artery_Tibial | ENSG00000155366.12 | -786052 | 441 | 219 | 2.06E-01 | -0.03 | 0.02 | 0.89 |
| Nerve_Tibial | ENSG00000155366.12 | -786052 | 414 | 202 | 2.06E-01 | -0.03 | 0.03 | 0.89 |
| Brain_Hypothalamus | ENSG00000233337.1 | 483868 | 121 | 61 | 2.07E-01 | 0.14 | 0.11 | 0.90 |
| Thyroid | ENSG00000143079.10 | -474799 | 446 | 224 | 2.08E-01 | 0.04 | 0.03 | 0.90 |
| Adrenal_Gland | ENSG00000007341.14 | -699443 | 190 | 96 | 2.09E-01 | -0.13 | 0.10 | 0.90 |
| Brain_Cerebellar_Hemisphere | ENSG00000273483.1 | -597059 | 136 | 74 | 2.11E-01 | 0.07 | 0.06 | 0.91 |
| Heart_Left_Ventricle | ENSG00000156171.10 | 781166 | 303 | 154 | 2.13E-01 | -0.07 | 0.06 | 0.91 |
| Colon_Transverse | ENSG00000116489.8 | -698415 | 274 | 119 | 2.13E-01 | 0.03 | 0.02 | 0.91 |
| Spleen | ENSG00000155367.11 | -794095 | 162 | 55 | 2.14E-01 | 0.13 | 0.10 | 0.91 |
| Breast_Mammary_Tissue | ENSG00000134255.9 | 781156 | 290 | 148 | 2.16E-01 | 0.05 | 0.04 | 0.92 |
| Brain_Hypothalamus | ENSG00000143110.7 | 447590 | 121 | 61 | 2.16E-01 | -0.09 | 0.07 | 0.92 |
| Uterus | ENSG00000155366.12 | -786052 | 111 | 63 | 2.17E-01 | -0.08 | 0.07 | 0.92 |
| Colon_Transverse | ENSG00000116459.6 | 472490 | 274 | 119 | 2.18E-01 | -0.03 | 0.02 | 0.92 |
| Brain_Caudate_basal_ganglia | ENSG00000155367.11 | -794095 | 160 | 88 | 2.18E-01 | -0.10 | 0.08 | 0.92 |
| Heart_Left_Ventricle | ENSG00000134255.9 | 781156 | 303 | 154 | 2.18E-01 | -0.07 | 0.06 | 0.92 |
| Colon_Transverse | ENSG00000238975.1 | -731206 | 274 | 119 | 2.20E-01 | 0.10 | 0.08 | 0.92 |
| Brain_Putamen_basal_ganglia | ENSG00000233337.1 | 483868 | 124 | 71 | 2.20E-01 | -0.16 | 0.13 | 0.92 |
| Skin_Sun_Exposed_Lower_leg | ENSG00000007341.14 | -699443 | 473 | 233 | 2.21E-01 | 0.05 | 0.04 | 0.92 |
| Brain_Hippocampus | ENSG00000273483.1 | -597059 | 123 | 63 | 2.21E-01 | 0.10 | 0.08 | 0.92 |
| Artery_Tibial | ENSG00000261654.1 | 979029 | 441 | 219 | 2.22E-01 | -0.08 | 0.07 | 0.92 |
| Cells_Transformed_fibroblasts | ENSG00000121931.11 | 968421 | 343 | 175 | 2.25E-01 | 0.04 | 0.04 | 0.93 |
| Brain_Cortex | ENSG00000260948.1 | 488314 | 158 | 79 | 2.25E-01 | 0.12 | 0.10 | 0.93 |
| Stomach | ENSG00000231346.1 | 312659 | 262 | 107 | 2.26E-01 | 0.07 | 0.06 | 0.93 |
| Lung | ENSG00000227811.2 | 181541 | 427 | 222 | 2.27E-01 | 0.09 | 0.07 | 0.93 |
| Skin_Sun_Exposed_Lower_leg | ENSG00000116455.9 | 472519 | 473 | 233 | 2.27E-01 | -0.04 | 0.03 | 0.93 |
| Cells_Transformed_fibroblasts | ENSG00000064886.9 | 720611 | 343 | 175 | 2.29E-01 | 0.05 | 0.04 | 0.93 |
| Brain_Putamen_basal_ganglia | ENSG00000215866.3 | -929261 | 124 | 71 | 2.29E-01 | 0.16 | 0.13 | 0.93 |

| | | | | | | | | |
|---|---|---|---|---|---|---|---|---|
| Minor_Salivary_Gland | ENSG00000162777.12 | 716847 | 97 | 49 | 2.31E-01 | -0.11 | 0.09 | 0.94 |
| Brain_Anterior_cingulate_cortex_BA24 | ENSG00000273010.1 | 958080 | 121 | 65 | 2.32E-01 | 0.16 | 0.13 | 0.94 |
| Esophagus_Muscularis | ENSG00000064703.7 | 166137 | 370 | 188 | 2.35E-01 | 0.05 | 0.04 | 0.94 |
| Adipose_Visceral_Omentum | ENSG00000134255.9 | 781156 | 355 | 172 | 2.35E-01 | 0.05 | 0.04 | 0.94 |
| Brain_Nucleus_accumbens_basal_ganglia | ENSG00000085465.11 | 493605 | 147 | 77 | 2.35E-01 | 0.08 | 0.06 | 0.94 |
| Esophagus_Muscularis | ENSG00000232811.1 | 977915 | 370 | 188 | 2.35E-01 | -0.10 | 0.08 | 0.94 |
| Skin_Not_Sun_Exposed_Suprapubic | ENSG00000162777.12 | 716847 | 387 | 200 | 2.36E-01 | -0.04 | 0.03 | 0.94 |
| Brain_Cerebellum | ENSG00000116473.10 | 379164 | 173 | 96 | 2.36E-01 | 0.06 | 0.05 | 0.94 |
| Colon_Transverse | ENSG00000233337.1 | 483868 | 274 | 119 | 2.36E-01 | -0.10 | 0.09 | 0.94 |
| Small_Intestine_Terminal_Ileum | ENSG00000155363.14 | -751759 | 137 | 54 | 2.36E-01 | -0.07 | 0.06 | 0.94 |
| Brain_Putamen_basal_ganglia | ENSG00000121933.13 | 357420 | 124 | 71 | 2.37E-01 | 0.12 | 0.10 | 0.94 |
| Esophagus_Gastroesophageal_Junction | ENSG00000143110.7 | 447590 | 244 | 110 | 2.39E-01 | -0.06 | 0.05 | 0.94 |
| Minor_Salivary_Gland | ENSG00000155363.14 | -751759 | 97 | 49 | 2.39E-01 | -0.10 | 0.08 | 0.94 |
| Esophagus_Gastroesophageal_Junction | ENSG00000007341.14 | -699443 | 244 | 110 | 2.40E-01 | -0.11 | 0.09 | 0.94 |
| Colon_Transverse | ENSG00000197852.8 | 168128 | 274 | 119 | 2.41E-01 | 0.07 | 0.06 | 0.94 |
| Esophagus_Mucosa | ENSG00000261654.1 | 979029 | 407 | 208 | 2.41E-01 | -0.09 | 0.08 | 0.94 |
| Breast_Mammary_Tissue | ENSG00000171385.5 | -67773 | 290 | 148 | 2.43E-01 | -0.08 | 0.07 | 0.94 |
| Stomach | ENSG00000197852.8 | 168128 | 262 | 107 | 2.43E-01 | -0.05 | 0.05 | 0.94 |
| Vagina | ENSG00000273010.1 | 958080 | 115 | 59 | 2.45E-01 | 0.17 | 0.15 | 0.94 |
| Thyroid | ENSG00000116489.8 | -698415 | 446 | 224 | 2.46E-01 | -0.03 | 0.03 | 0.94 |
| Lung | ENSG00000155367.11 | -794095 | 427 | 222 | 2.46E-01 | 0.03 | 0.03 | 0.94 |
| Adipose_Subcutaneous | ENSG00000116455.9 | 472519 | 442 | 214 | 2.46E-01 | -0.03 | 0.03 | 0.94 |
| Adipose_Visceral_Omentum | ENSG00000225075.1 | -775239 | 355 | 172 | 2.46E-01 | 0.09 | 0.08 | 0.94 |
| Ovary | ENSG00000261654.1 | 979029 | 133 | 71 | 2.48E-01 | -0.12 | 0.10 | 0.94 |
| Thyroid | ENSG00000227811.2 | 181541 | 446 | 224 | 2.48E-01 | 0.09 | 0.08 | 0.94 |
| Brain_Anterior_cingulate_cortex_BA24 | ENSG00000143079.10 | -474799 | 121 | 65 | 2.49E-01 | 0.08 | 0.07 | 0.94 |
| Stomach | ENSG00000231437.3 | -68388 | 262 | 107 | 2.49E-01 | 0.07 | 0.06 | 0.94 |
| Uterus | ENSG00000143110.7 | 447590 | 111 | 63 | 2.49E-01 | 0.13 | 0.11 | 0.94 |
| Uterus | ENSG00000116459.6 | 472490 | 111 | 63 | 2.49E-01 | -0.06 | 0.05 | 0.94 |
| Adrenal_Gland | ENSG00000156171.10 | 781166 | 190 | 96 | 2.51E-01 | -0.08 | 0.07 | 0.94 |
| Artery_Tibial | ENSG00000064703.7 | 166137 | 441 | 219 | 2.51E-01 | 0.03 | 0.03 | 0.94 |
| Skin_Not_Sun_Exposed_Suprapubic | ENSG00000085465.11 | 493605 | 387 | 200 | 2.52E-01 | -0.04 | 0.03 | 0.94 |
| Prostate | ENSG00000156171.10 | 781166 | 152 | 75 | 2.52E-01 | -0.12 | 0.10 | 0.94 |
| Brain_Nucleus_accumbens_basal_ganglia | ENSG00000197852.8 | 168128 | 147 | 77 | 2.52E-01 | -0.11 | 0.10 | 0.94 |
| Brain_Substantia_nigra | ENSG00000231346.1 | 312659 | 88 | 49 | 2.52E-01 | 0.17 | 0.14 | 0.94 |
| Brain_Caudate_basal_ganglia | ENSG00000064703.7 | 166137 | 160 | 88 | 2.54E-01 | -0.10 | 0.09 | 0.94 |
| Brain_Spinal_cord_cervical_c-1 | ENSG00000231346.1 | 312659 | 91 | 49 | 2.54E-01 | -0.20 | 0.17 | 0.94 |
| Heart_Atrial_Appendage | ENSG00000156171.10 | 781166 | 297 | 147 | 2.55E-01 | -0.07 | 0.06 | 0.94 |
| Whole_Blood | ENSG00000116473.10 | 379164 | 407 | 195 | 2.55E-01 | -0.02 | 0.02 | 0.94 |
| Brain_Substantia_nigra | ENSG00000215866.3 | -929261 | 88 | 49 | 2.57E-01 | 0.15 | 0.13 | 0.94 |
| Brain_Frontal_Cortex_BA9 | ENSG00000197852.8 | 168128 | 129 | 73 | 2.57E-01 | -0.07 | 0.06 | 0.94 |
| Vagina | ENSG00000116489.8 | -698415 | 115 | 59 | 2.58E-01 | -0.07 | 0.06 | 0.94 |
| Prostate | ENSG00000155366.12 | -786052 | 152 | 75 | 2.59E-01 | -0.09 | 0.08 | 0.94 |
| Heart_Left_Ventricle | ENSG00000260948.1 | 488314 | 303 | 154 | 2.59E-01 | 0.09 | 0.08 | 0.94 |
| Nerve_Tibial | ENSG00000231437.3 | -68388 | 414 | 202 | 2.60E-01 | 0.07 | 0.06 | 0.94 |
| Esophagus_Mucosa | ENSG00000173947.9 | 575094 | 407 | 208 | 2.61E-01 | -0.06 | 0.05 | 0.94 |

| | | | | | | | | |
|---|---|---|---|---|---|---|---|---|
| Brain_Hypothalamus | ENSG00000121931.11 | 968421 | 121 | 61 | 2.61E-01 | 0.10 | 0.09 | 0.94 |
| Adrenal_Gland | ENSG00000116473.10 | 379164 | 190 | 96 | 2.61E-01 | 0.06 | 0.06 | 0.94 |
| Breast_Mammary_Tissue | ENSG00000273010.1 | 958080 | 290 | 148 | 2.61E-01 | -0.10 | 0.09 | 0.94 |
| Ovary | ENSG00000243960.1 | 482557 | 133 | 71 | 2.62E-01 | -0.15 | 0.13 | 0.94 |
| Brain_Cerebellar_Hemisphere | ENSG00000134245.13 | -545159 | 136 | 74 | 2.63E-01 | -0.08 | 0.07 | 0.94 |
| Colon_Sigmoid | ENSG00000243960.1 | 482557 | 233 | 107 | 2.67E-01 | -0.12 | 0.10 | 0.94 |
| Esophagus_Gastroesophageal_Junction | ENSG00000116459.6 | 472490 | 244 | 110 | 2.67E-01 | 0.04 | 0.03 | 0.94 |
| Esophagus_Muscularis | ENSG00000243960.1 | 482557 | 370 | 188 | 2.67E-01 | 0.10 | 0.09 | 0.94 |
| Breast_Mammary_Tissue | ENSG00000232811.1 | 977915 | 290 | 148 | 2.68E-01 | -0.09 | 0.08 | 0.94 |
| Brain_Anterior_cingulate_cortex_BA24 | ENSG00000134245.13 | -545159 | 121 | 65 | 2.69E-01 | -0.13 | 0.11 | 0.94 |
| Adrenal_Gland | ENSG00000085465.11 | 493605 | 190 | 96 | 2.69E-01 | -0.07 | 0.07 | 0.94 |
| Artery_Tibial | ENSG00000116473.10 | 379164 | 441 | 219 | 2.69E-01 | -0.03 | 0.02 | 0.94 |
| Artery_Coronary | ENSG00000064886.9 | 720611 | 173 | 81 | 2.71E-01 | -0.10 | 0.09 | 0.94 |
| Cells_Transformed_fibroblasts | ENSG00000231437.3 | -68388 | 343 | 175 | 2.72E-01 | 0.05 | 0.05 | 0.94 |
| Stomach | ENSG00000261654.1 | 979029 | 262 | 107 | 2.72E-01 | -0.10 | 0.09 | 0.94 |
| Brain_Putamen_basal_ganglia | ENSG00000273483.1 | -597059 | 124 | 71 | 2.73E-01 | 0.12 | 0.11 | 0.94 |
| Brain_Putamen_basal_ganglia | ENSG00000143079.10 | -474799 | 124 | 71 | 2.73E-01 | 0.09 | 0.08 | 0.94 |
| Esophagus_Gastroesophageal_Junction | ENSG00000116473.10 | 379164 | 244 | 110 | 2.73E-01 | -0.04 | 0.04 | 0.94 |
| Uterus | ENSG00000155367.11 | -794095 | 111 | 63 | 2.73E-01 | -0.12 | 0.11 | 0.94 |
| Brain_Anterior_cingulate_cortex_BA24 | ENSG00000116459.6 | 472490 | 121 | 65 | 2.73E-01 | -0.07 | 0.07 | 0.94 |
| Pancreas | ENSG00000064703.7 | 166137 | 248 | 108 | 2.75E-01 | -0.08 | 0.08 | 0.94 |
| Pancreas | ENSG00000156171.10 | 781166 | 248 | 108 | 2.75E-01 | -0.07 | 0.07 | 0.94 |
| Lung | ENSG00000173947.9 | 575094 | 427 | 222 | 2.77E-01 | -0.03 | 0.03 | 0.94 |
| Brain_Nucleus_accumbens_basal_ganglia | ENSG00000143110.7 | 447590 | 147 | 77 | 2.78E-01 | -0.08 | 0.07 | 0.94 |
| Small_Intestine_Terminal_Ileum | ENSG00000116455.9 | 472519 | 137 | 54 | 2.78E-01 | -0.07 | 0.07 | 0.94 |
| Lung | ENSG00000197852.8 | 168128 | 427 | 222 | 2.79E-01 | 0.05 | 0.04 | 0.94 |
| Liver | ENSG00000156171.10 | 781166 | 175 | 88 | 2.79E-01 | -0.12 | 0.11 | 0.94 |
| Brain_Cerebellum | ENSG00000155363.14 | -751759 | 173 | 96 | 2.80E-01 | -0.05 | 0.05 | 0.94 |
| Brain_Frontal_Cortex_BA9 | ENSG00000260948.1 | 488314 | 129 | 73 | 2.80E-01 | -0.11 | 0.10 | 0.94 |
| Adipose_Visceral_Omentum | ENSG00000162777.12 | 716847 | 355 | 172 | 2.80E-01 | 0.05 | 0.04 | 0.94 |
| Minor_Salivary_Gland | ENSG00000143110.7 | 447590 | 97 | 49 | 2.81E-01 | 0.10 | 0.09 | 0.94 |
| Vagina | ENSG00000155366.12 | -786052 | 115 | 59 | 2.81E-01 | 0.06 | 0.06 | 0.94 |
| Lung | ENSG00000260948.1 | 488314 | 427 | 222 | 2.81E-01 | 0.04 | 0.04 | 0.94 |
| Esophagus_Mucosa | ENSG00000273483.1 | -597059 | 407 | 208 | 2.83E-01 | 0.07 | 0.06 | 0.94 |
| Spleen | ENSG00000064703.7 | 166137 | 162 | 55 | 2.83E-01 | 0.13 | 0.12 | 0.94 |
| Brain_Amygdala | ENSG00000162777.12 | 716847 | 100 | 47 | 2.83E-01 | -0.12 | 0.11 | 0.94 |
| Liver | ENSG00000243960.1 | 482557 | 175 | 88 | 2.84E-01 | -0.12 | 0.12 | 0.94 |
| Skin_Not_Sun_Exposed_Suprapubic | ENSG00000143079.10 | -474799 | 387 | 200 | 2.84E-01 | 0.03 | 0.02 | 0.94 |
| Brain_Cerebellum | ENSG00000197852.8 | 168128 | 173 | 96 | 2.85E-01 | -0.08 | 0.08 | 0.94 |
| Lung | ENSG00000243960.1 | 482557 | 427 | 222 | 2.86E-01 | 0.07 | 0.07 | 0.94 |
| Brain_Cerebellum | ENSG00000243960.1 | 482557 | 173 | 96 | 2.87E-01 | 0.12 | 0.11 | 0.94 |
| Brain_Hypothalamus | ENSG00000156171.10 | 781166 | 121 | 61 | 2.87E-01 | -0.11 | 0.10 | 0.94 |
| Brain_Hippocampus | ENSG00000064886.9 | 720611 | 123 | 63 | 2.88E-01 | 0.11 | 0.10 | 0.94 |
| Skin_Sun_Exposed_Lower_leg | ENSG00000231437.3 | -68388 | 473 | 233 | 2.88E-01 | 0.05 | 0.05 | 0.94 |
| Esophagus_Gastroesophageal_Junction | ENSG00000085465.11 | 493605 | 244 | 110 | 2.89E-01 | -0.05 | 0.04 | 0.94 |
| Colon_Transverse | ENSG00000143110.7 | 447590 | 274 | 119 | 2.89E-01 | 0.05 | 0.05 | 0.94 |
| Stomach | ENSG00000143079.10 | -474799 | 262 | 107 | 2.89E-01 | -0.04 | 0.04 | 0.94 |

| | | | | | | | | |
|---|---|---|---|---|---|---|---|---|
| Brain_Caudate_basal_ganglia | ENSG00000143110.7 | 447590 | 160 | 88 | 2.89E-01 | -0.06 | 0.06 | 0.94 |
| Brain_Frontal_Cortex_BA9 | ENSG00000231437.3 | -68388 | 129 | 73 | 2.90E-01 | 0.13 | 0.12 | 0.94 |
| Brain_Cortex | ENSG00000143079.10 | -474799 | 158 | 79 | 2.92E-01 | 0.09 | 0.08 | 0.94 |
| Cells_EBV-transformed_lymphocytes | ENSG00000238975.1 | -731206 | 130 | 68 | 2.92E-01 | -0.10 | 0.09 | 0.94 |
| Breast_Mammary_Tissue | ENSG00000143079.10 | -474799 | 290 | 148 | 2.93E-01 | 0.03 | 0.03 | 0.94 |
| Nerve_Tibial | ENSG00000116473.10 | 379164 | 414 | 202 | 2.94E-01 | -0.04 | 0.03 | 0.94 |
| Liver | ENSG00000232811.1 | 977915 | 175 | 88 | 2.94E-01 | 0.12 | 0.11 | 0.94 |
| Minor_Salivary_Gland | ENSG00000155366.12 | -786052 | 97 | 49 | 2.95E-01 | 0.06 | 0.06 | 0.94 |
| Pancreas | ENSG00000238975.1 | -731206 | 248 | 108 | 2.95E-01 | -0.11 | 0.11 | 0.94 |
| Brain_Amygdala | ENSG00000171385.5 | -67773 | 100 | 47 | 2.95E-01 | -0.07 | 0.07 | 0.94 |
| Colon_Sigmoid | ENSG00000225075.1 | -775239 | 233 | 107 | 2.96E-01 | -0.11 | 0.10 | 0.94 |
| Artery_Tibial | ENSG00000134255.9 | 781156 | 441 | 219 | 2.96E-01 | -0.04 | 0.04 | 0.94 |
| Brain_Caudate_basal_ganglia | ENSG00000007341.14 | -699443 | 160 | 88 | 2.97E-01 | -0.12 | 0.11 | 0.94 |
| Testis | ENSG00000134255.9 | 781156 | 259 | 128 | 2.97E-01 | 0.03 | 0.03 | 0.94 |
| Heart_Atrial_Appendage | ENSG00000064703.7 | 166137 | 297 | 147 | 2.97E-01 | 0.04 | 0.04 | 0.94 |
| Adipose_Visceral_Omentum | ENSG00000243960.1 | 482557 | 355 | 172 | 2.98E-01 | -0.07 | 0.07 | 0.94 |
| Artery_Aorta | ENSG00000007341.14 | -699443 | 299 | 152 | 2.98E-01 | -0.06 | 0.06 | 0.94 |
| Adrenal_Gland | ENSG00000231437.3 | -68388 | 190 | 96 | 2.99E-01 | 0.10 | 0.10 | 0.94 |
| Adipose_Visceral_Omentum | ENSG00000116489.8 | -698415 | 355 | 172 | 2.99E-01 | -0.03 | 0.03 | 0.94 |
| Breast_Mammary_Tissue | ENSG00000238975.1 | -731206 | 290 | 148 | 2.99E-01 | -0.08 | 0.08 | 0.94 |
| Brain_Cerebellar_Hemisphere | ENSG00000273010.1 | 958080 | 136 | 74 | 3.00E-01 | 0.13 | 0.12 | 0.94 |
| Liver | ENSG00000155363.14 | -751759 | 175 | 88 | 3.01E-01 | -0.07 | 0.06 | 0.94 |
| Muscle_Skeletal | ENSG00000197852.8 | 168128 | 564 | 276 | 3.01E-01 | -0.04 | 0.04 | 0.94 |
| Artery_Aorta | ENSG00000121933.13 | 357420 | 299 | 152 | 3.01E-01 | -0.04 | 0.04 | 0.94 |
| Brain_Nucleus_accumbens_basal_ganglia | ENSG00000155363.14 | -751759 | 147 | 77 | 3.01E-01 | -0.08 | 0.07 | 0.94 |
| Colon_Sigmoid | ENSG00000162777.12 | 716847 | 233 | 107 | 3.02E-01 | 0.06 | 0.05 | 0.94 |
| Spleen | ENSG00000273483.1 | -597059 | 162 | 55 | 3.02E-01 | 0.12 | 0.11 | 0.94 |
| Liver | ENSG00000261654.1 | 979029 | 175 | 88 | 3.02E-01 | -0.14 | 0.13 | 0.94 |
| Artery_Aorta | ENSG00000116473.10 | 379164 | 299 | 152 | 3.02E-01 | -0.04 | 0.04 | 0.94 |
| Artery_Tibial | ENSG00000197852.8 | 168128 | 441 | 219 | 3.02E-01 | -0.04 | 0.04 | 0.94 |
| Ovary | ENSG00000225075.1 | -775239 | 133 | 71 | 3.02E-01 | 0.13 | 0.12 | 0.94 |
| Heart_Left_Ventricle | ENSG00000155366.12 | -786052 | 303 | 154 | 3.03E-01 | -0.03 | 0.03 | 0.94 |
| Uterus | ENSG00000171385.5 | -67773 | 111 | 63 | 3.03E-01 | -0.11 | 0.11 | 0.94 |
| Testis | ENSG00000232811.1 | 977915 | 259 | 128 | 3.05E-01 | -0.07 | 0.07 | 0.94 |
| Ovary | ENSG00000273010.1 | 958080 | 133 | 71 | 3.05E-01 | 0.13 | 0.12 | 0.94 |
| Adipose_Visceral_Omentum | ENSG00000064886.9 | 720611 | 355 | 172 | 3.06E-01 | -0.06 | 0.06 | 0.94 |
| Skin_Not_Sun_Exposed_Suprapubic | ENSG00000116459.6 | 472490 | 387 | 200 | 3.06E-01 | 0.03 | 0.02 | 0.94 |
| Uterus | ENSG00000233337.1 | 483868 | 111 | 63 | 3.06E-01 | 0.12 | 0.11 | 0.94 |
| Brain_Cerebellum | ENSG00000064703.7 | 166137 | 173 | 96 | 3.06E-01 | -0.08 | 0.08 | 0.94 |
| Brain_Substantia_nigra | ENSG00000121933.13 | 357420 | 88 | 49 | 3.07E-01 | 0.07 | 0.07 | 0.94 |
| Brain_Frontal_Cortex_BA9 | ENSG00000064886.9 | 720611 | 129 | 73 | 3.08E-01 | 0.11 | 0.10 | 0.94 |
| Cells_Transformed_fibroblasts | ENSG00000231346.1 | 312659 | 343 | 175 | 3.08E-01 | 0.07 | 0.07 | 0.94 |
| Muscle_Skeletal | ENSG00000162777.12 | 716847 | 564 | 276 | 3.08E-01 | -0.04 | 0.04 | 0.94 |
| Cells_EBV-transformed_lymphocytes | ENSG00000116459.6 | 472490 | 130 | 68 | 3.09E-01 | 0.05 | 0.05 | 0.94 |
| Testis | ENSG00000231437.3 | -68388 | 259 | 128 | 3.09E-01 | 0.08 | 0.08 | 0.94 |
| Brain_Cortex | ENSG00000007341.14 | -699443 | 158 | 79 | 3.09E-01 | 0.11 | 0.11 | 0.94 |
| Heart_Atrial_Appendage | ENSG00000007341.14 | -699443 | 297 | 147 | 3.11E-01 | -0.08 | 0.08 | 0.94 |

| | | | | | | | | |
|---|---|---|---|---|---|---|---|---|
| Brain_Nucleus_accumbens_basal_ganglia | ENSG00000231246.1 | -439146 | 147 | 77 | 3.11E-01 | -0.12 | 0.12 | 0.94 |
| Brain_Frontal_Cortex_BA9 | ENSG00000007341.14 | -699443 | 129 | 73 | 3.12E-01 | -0.13 | 0.12 | 0.94 |
| Artery_Coronary | ENSG00000121933.13 | 357420 | 173 | 81 | 3.16E-01 | -0.06 | 0.06 | 0.94 |
| Brain_Substantia_nigra | ENSG00000116455.9 | 472519 | 88 | 49 | 3.16E-01 | 0.10 | 0.10 | 0.94 |
| Brain_Spinal_cord_cervical_c-1 | ENSG00000215866.3 | -929261 | 91 | 49 | 3.16E-01 | -0.13 | 0.13 | 0.94 |
| Adipose_Subcutaneous | ENSG00000156171.10 | 781166 | 442 | 214 | 3.16E-01 | -0.05 | 0.04 | 0.94 |
| Brain_Nucleus_accumbens_basal_ganglia | ENSG00000116473.10 | 379164 | 147 | 77 | 3.17E-01 | -0.05 | 0.05 | 0.94 |
| Muscle_Skeletal | ENSG00000116489.8 | -698415 | 564 | 276 | 3.17E-01 | 0.03 | 0.03 | 0.94 |
| Brain_Cortex | ENSG00000116473.10 | 379164 | 158 | 79 | 3.18E-01 | -0.06 | 0.06 | 0.94 |
| Brain_Cortex | ENSG00000155363.14 | -751759 | 158 | 79 | 3.19E-01 | -0.06 | 0.06 | 0.94 |
| Prostate | ENSG00000184599.9 | -799037 | 152 | 75 | 3.20E-01 | -0.10 | 0.10 | 0.94 |
| Artery_Aorta | ENSG00000273483.1 | -597059 | 299 | 152 | 3.20E-01 | 0.07 | 0.07 | 0.94 |
| Brain_Hippocampus | ENSG00000232811.1 | 977915 | 123 | 63 | 3.21E-01 | -0.15 | 0.15 | 0.94 |
| Pancreas | ENSG00000243960.1 | 482557 | 248 | 108 | 3.23E-01 | 0.09 | 0.09 | 0.94 |
| Minor_Salivary_Gland | ENSG00000134245.13 | -545159 | 97 | 49 | 3.24E-01 | -0.10 | 0.10 | 0.94 |
| Brain_Frontal_Cortex_BA9 | ENSG00000227811.2 | 181541 | 129 | 73 | 3.25E-01 | 0.12 | 0.12 | 0.94 |
| Skin_Sun_Exposed_Lower_leg | ENSG00000173947.9 | 575094 | 473 | 233 | 3.25E-01 | 0.05 | 0.05 | 0.94 |
| Pituitary | ENSG00000116473.10 | 379164 | 183 | 95 | 3.26E-01 | -0.06 | 0.06 | 0.94 |
| Esophagus_Gastroesophageal_Junction | ENSG00000225075.1 | -775239 | 244 | 110 | 3.26E-01 | -0.10 | 0.10 | 0.94 |
| Pituitary | ENSG00000231437.3 | -68388 | 183 | 95 | 3.26E-01 | 0.10 | 0.10 | 0.94 |
| Artery_Aorta | ENSG00000231437.3 | -68388 | 299 | 152 | 3.26E-01 | 0.06 | 0.06 | 0.94 |
| Pancreas | ENSG00000232811.1 | 977915 | 248 | 108 | 3.26E-01 | 0.11 | 0.11 | 0.94 |
| Thyroid | ENSG00000173947.9 | 575094 | 446 | 224 | 3.27E-01 | 0.03 | 0.04 | 0.94 |
| Thyroid | ENSG00000273483.1 | -597059 | 446 | 224 | 3.27E-01 | -0.06 | 0.06 | 0.94 |
| Pituitary | ENSG00000121931.11 | 968421 | 183 | 95 | 3.28E-01 | 0.08 | 0.08 | 0.94 |
| Artery_Coronary | ENSG00000260948.1 | 488314 | 173 | 81 | 3.29E-01 | -0.09 | 0.09 | 0.94 |
| Brain_Hypothalamus | ENSG00000184599.9 | -799037 | 121 | 61 | 3.29E-01 | 0.14 | 0.14 | 0.94 |
| Brain_Cerebellum | ENSG00000232558.1 | 64477 | 173 | 96 | 3.30E-01 | 0.12 | 0.12 | 0.94 |
| Heart_Left_Ventricle | ENSG00000155367.11 | -794095 | 303 | 154 | 3.30E-01 | 0.05 | 0.05 | 0.94 |
| Colon_Sigmoid | ENSG00000116473.10 | 379164 | 233 | 107 | 3.30E-01 | -0.05 | 0.06 | 0.94 |
| Breast_Mammary_Tissue | ENSG00000156171.10 | 781166 | 290 | 148 | 3.30E-01 | 0.04 | 0.04 | 0.94 |
| Artery_Coronary | ENSG00000121931.11 | 968421 | 173 | 81 | 3.31E-01 | -0.09 | 0.10 | 0.94 |
| Ovary | ENSG00000116473.10 | 379164 | 133 | 71 | 3.31E-01 | -0.07 | 0.07 | 0.94 |
| Colon_Sigmoid | ENSG00000173947.9 | 575094 | 233 | 107 | 3.32E-01 | -0.06 | 0.06 | 0.94 |
| Cells_Transformed_fibroblasts | ENSG00000224167.1 | -955558 | 343 | 175 | 3.32E-01 | 0.07 | 0.08 | 0.94 |
| Colon_Sigmoid | ENSG00000171385.5 | -67773 | 233 | 107 | 3.33E-01 | -0.07 | 0.07 | 0.94 |
| Artery_Tibial | ENSG00000121931.11 | 968421 | 441 | 219 | 3.34E-01 | -0.04 | 0.04 | 0.94 |
| Heart_Left_Ventricle | ENSG00000162777.12 | 716847 | 303 | 154 | 3.34E-01 | -0.05 | 0.05 | 0.94 |
| Adipose_Subcutaneous | ENSG00000231437.3 | -68388 | 442 | 214 | 3.35E-01 | 0.05 | 0.06 | 0.94 |
| Nerve_Tibial | ENSG00000064886.9 | 720611 | 414 | 202 | 3.35E-01 | -0.05 | 0.05 | 0.94 |
| Colon_Sigmoid | ENSG00000116489.8 | -698415 | 233 | 107 | 3.36E-01 | -0.05 | 0.06 | 0.94 |
| Artery_Aorta | ENSG00000261654.1 | 979029 | 299 | 152 | 3.36E-01 | 0.08 | 0.08 | 0.94 |
| Adipose_Subcutaneous | ENSG00000134216.14 | 630520 | 442 | 214 | 3.37E-01 | -0.06 | 0.07 | 0.94 |
| Brain_Hippocampus | ENSG00000155363.14 | -751759 | 123 | 63 | 3.38E-01 | -0.06 | 0.06 | 0.94 |
| Artery_Tibial | ENSG00000143079.10 | -474799 | 441 | 219 | 3.38E-01 | -0.03 | 0.03 | 0.94 |
| Vagina | ENSG00000134255.9 | 781156 | 115 | 59 | 3.39E-01 | -0.08 | 0.09 | 0.94 |

| | | | | | | | | |
|---|---|---|---|---|---|---|---|---|
| Vagina | ENSG00000162777.12 | 716847 | 115 | 59 | 3.39E-01 | 0.05 | 0.05 | 0.94 |
| Thyroid | ENSG00000116459.6 | 472490 | 446 | 224 | 3.39E-01 | -0.01 | 0.01 | 0.94 |
| Brain_Putamen_basal_ganglia | ENSG00000155367.11 | -794095 | 124 | 71 | 3.39E-01 | -0.10 | 0.10 | 0.94 |
| Ovary | ENSG00000155363.14 | -751759 | 133 | 71 | 3.40E-01 | 0.06 | 0.07 | 0.94 |
| Brain_Hippocampus | ENSG00000171385.5 | -67773 | 123 | 63 | 3.42E-01 | 0.06 | 0.06 | 0.94 |
| Muscle_Skeletal | ENSG00000007341.14 | -699443 | 564 | 276 | 3.42E-01 | -0.05 | 0.05 | 0.94 |
| Brain_Amygdala | ENSG00000227811.2 | 181541 | 100 | 47 | 3.43E-01 | -0.17 | 0.18 | 0.94 |
| Prostate | ENSG00000260948.1 | 488314 | 152 | 75 | 3.43E-01 | 0.10 | 0.10 | 0.94 |
| Thyroid | ENSG00000155367.11 | -794095 | 446 | 224 | 3.43E-01 | -0.04 | 0.04 | 0.94 |
| Pituitary | ENSG00000224167.1 | -955558 | 183 | 95 | 3.44E-01 | -0.14 | 0.14 | 0.94 |
| Brain_Frontal_Cortex_BA9 | ENSG00000273483.1 | -597059 | 129 | 73 | 3.45E-01 | 0.07 | 0.07 | 0.94 |
| Brain_Spinal_cord_cervical_c-1 | ENSG00000184599.9 | -799037 | 91 | 49 | 3.45E-01 | -0.14 | 0.15 | 0.94 |
| Testis | ENSG00000215866.3 | -929261 | 259 | 128 | 3.45E-01 | 0.04 | 0.04 | 0.94 |
| Brain_Hippocampus | ENSG00000224167.1 | -955558 | 123 | 63 | 3.45E-01 | -0.13 | 0.14 | 0.94 |
| Adipose_Subcutaneous | ENSG00000243960.1 | 482557 | 442 | 214 | 3.46E-01 | -0.07 | 0.08 | 0.94 |
| Nerve_Tibial | ENSG00000171385.5 | -67773 | 414 | 202 | 3.47E-01 | 0.04 | 0.05 | 0.94 |
| Artery_Coronary | ENSG00000134255.9 | 781156 | 173 | 81 | 3.47E-01 | -0.06 | 0.06 | 0.94 |
| Uterus | ENSG00000116473.10 | 379164 | 111 | 63 | 3.47E-01 | -0.05 | 0.06 | 0.94 |
| Minor_Salivary_Gland | ENSG00000184599.9 | -799037 | 97 | 49 | 3.49E-01 | 0.13 | 0.14 | 0.94 |
| Cells_Transformed_fibroblasts | ENSG00000155363.14 | -751759 | 343 | 175 | 3.49E-01 | -0.03 | 0.03 | 0.94 |
| Brain_Caudate_basal_ganglia | ENSG00000156171.10 | 781166 | 160 | 88 | 3.49E-01 | -0.07 | 0.08 | 0.94 |
| Stomach | ENSG00000233337.1 | 483868 | 262 | 107 | 3.50E-01 | -0.07 | 0.08 | 0.94 |
| Whole_Blood | ENSG00000143110.7 | 447590 | 407 | 195 | 3.50E-01 | -0.02 | 0.02 | 0.94 |
| Adrenal_Gland | ENSG00000243960.1 | 482557 | 190 | 96 | 3.50E-01 | -0.10 | 0.11 | 0.94 |
| Adipose_Subcutaneous | ENSG00000227179.2 | 529563 | 442 | 214 | 3.52E-01 | 0.05 | 0.05 | 0.94 |
| Lung | ENSG00000064886.9 | 720611 | 427 | 222 | 3.52E-01 | 0.05 | 0.05 | 0.94 |
| Brain_Hippocampus | ENSG00000116473.10 | 379164 | 123 | 63 | 3.53E-01 | -0.05 | 0.05 | 0.94 |
| Brain_Spinal_cord_cervical_c-1 | ENSG00000007341.14 | -699443 | 91 | 49 | 3.53E-01 | -0.14 | 0.15 | 0.94 |
| Minor_Salivary_Gland | ENSG00000231437.3 | -68388 | 97 | 49 | 3.53E-01 | -0.11 | 0.12 | 0.94 |
| Brain_Hippocampus | ENSG00000162777.12 | 716847 | 123 | 63 | 3.55E-01 | 0.09 | 0.10 | 0.94 |
| Prostate | ENSG00000064886.9 | 720611 | 152 | 75 | 3.55E-01 | -0.08 | 0.08 | 0.94 |
| Liver | ENSG00000197852.8 | 168128 | 175 | 88 | 3.56E-01 | -0.10 | 0.10 | 0.94 |
| Brain_Spinal_cord_cervical_c-1 | ENSG00000224167.1 | -955558 | 91 | 49 | 3.57E-01 | -0.15 | 0.16 | 0.94 |
| Adipose_Visceral_Omentum | ENSG00000231246.1 | -439146 | 355 | 172 | 3.57E-01 | 0.07 | 0.07 | 0.94 |
| Brain_Nucleus_accumbens_basal_ganglia | ENSG00000121933.13 | 357420 | 147 | 77 | 3.58E-01 | -0.07 | 0.07 | 0.94 |
| Brain_Amygdala | ENSG00000224167.1 | -955558 | 100 | 47 | 3.58E-01 | -0.15 | 0.16 | 0.94 |
| Cells_Transformed_fibroblasts | ENSG00000238975.1 | -731206 | 343 | 175 | 3.58E-01 | -0.07 | 0.08 | 0.94 |
| Stomach | ENSG00000243960.1 | 482557 | 262 | 107 | 3.58E-01 | -0.08 | 0.08 | 0.94 |
| Esophagus_Gastroesophageal_Junction | ENSG00000121931.11 | 968421 | 244 | 110 | 3.58E-01 | -0.06 | 0.07 | 0.94 |
| Vagina | ENSG00000231346.1 | 312659 | 115 | 59 | 3.59E-01 | 0.08 | 0.08 | 0.94 |
| Brain_Frontal_Cortex_BA9 | ENSG00000121933.13 | 357420 | 129 | 73 | 3.59E-01 | 0.09 | 0.10 | 0.94 |
| Artery_Tibial | ENSG00000064886.9 | 720611 | 441 | 219 | 3.61E-01 | -0.05 | 0.05 | 0.94 |
| Brain_Cortex | ENSG00000233337.1 | 483868 | 158 | 79 | 3.62E-01 | -0.12 | 0.13 | 0.94 |
| Artery_Coronary | ENSG00000197852.8 | 168128 | 173 | 81 | 3.62E-01 | -0.06 | 0.06 | 0.94 |
| Brain_Substantia_nigra | ENSG00000156171.10 | 781166 | 88 | 49 | 3.63E-01 | 0.09 | 0.10 | 0.94 |
| Ovary | ENSG00000007341.14 | -699443 | 133 | 71 | 3.63E-01 | 0.08 | 0.08 | 0.94 |
| Liver | ENSG00000134245.13 | -545159 | 175 | 88 | 3.64E-01 | -0.09 | 0.10 | 0.94 |

| | | | | | | | | |
|---|---|---|---|---|---|---|---|---|
| Skin_Sun_Exposed_Lower_leg | ENSG00000231246.1 | -439146 | 473 | 233 | 3.64E-01 | 0.05 | 0.06 | 0.94 |
| Testis | ENSG00000234020.1 | 553973 | 259 | 128 | 3.65E-01 | -0.08 | 0.09 | 0.94 |
| Testis | ENSG00000121933.13 | 357420 | 259 | 128 | 3.66E-01 | -0.03 | 0.03 | 0.94 |
| Adipose_Visceral_Omentum | ENSG00000134245.13 | -545159 | 355 | 172 | 3.66E-01 | -0.03 | 0.04 | 0.94 |
| Brain_Caudate_basal_ganglia | ENSG00000273483.1 | -597059 | 160 | 88 | 3.67E-01 | 0.07 | 0.08 | 0.94 |
| Breast_Mammary_Tissue | ENSG00000143110.7 | 447590 | 290 | 148 | 3.67E-01 | -0.03 | 0.03 | 0.94 |
| Uterus | ENSG00000116455.9 | 472519 | 111 | 63 | 3.68E-01 | 0.05 | 0.05 | 0.94 |
| Heart_Left_Ventricle | ENSG00000273010.1 | 958080 | 303 | 154 | 3.69E-01 | 0.08 | 0.09 | 0.94 |
| Adrenal_Gland | ENSG00000215866.3 | -929261 | 190 | 96 | 3.70E-01 | -0.09 | 0.10 | 0.94 |
| Uterus | ENSG00000184599.9 | -799037 | 111 | 63 | 3.71E-01 | -0.10 | 0.12 | 0.94 |
| Brain_Substantia_nigra | ENSG00000155363.14 | -751759 | 88 | 49 | 3.71E-01 | 0.09 | 0.10 | 0.94 |
| Artery_Aorta | ENSG00000134245.13 | -545159 | 299 | 152 | 3.72E-01 | -0.05 | 0.06 | 0.94 |
| Uterus | ENSG00000007341.14 | -699443 | 111 | 63 | 3.73E-01 | -0.07 | 0.08 | 0.94 |
| Colon_Sigmoid | ENSG00000155366.12 | -786052 | 233 | 107 | 3.73E-01 | 0.04 | 0.05 | 0.94 |
| Brain_Nucleus_accumbens_basal_ganglia | ENSG00000064703.7 | 166137 | 147 | 77 | 3.73E-01 | -0.09 | 0.10 | 0.94 |
| Adrenal_Gland | ENSG00000232811.1 | 977915 | 190 | 96 | 3.74E-01 | 0.09 | 0.10 | 0.94 |
| Breast_Mammary_Tissue | ENSG00000116455.9 | 472519 | 290 | 148 | 3.74E-01 | 0.03 | 0.03 | 0.94 |
| Adrenal_Gland | ENSG00000173947.9 | 575094 | 190 | 96 | 3.74E-01 | -0.05 | 0.06 | 0.94 |
| Skin_Not_Sun_Exposed_Suprapubic | ENSG00000116455.9 | 472519 | 387 | 200 | 3.75E-01 | 0.03 | 0.03 | 0.94 |
| Skin_Sun_Exposed_Lower_leg | ENSG00000156171.10 | 781166 | 473 | 233 | 3.75E-01 | -0.04 | 0.04 | 0.94 |
| Colon_Sigmoid | ENSG00000261654.1 | 979029 | 233 | 107 | 3.76E-01 | 0.08 | 0.09 | 0.94 |
| Brain_Nucleus_accumbens_basal_ganglia | ENSG00000155366.12 | -786052 | 147 | 77 | 3.76E-01 | -0.04 | 0.05 | 0.94 |
| Esophagus_Muscularis | ENSG00000155367.11 | -794095 | 370 | 188 | 3.78E-01 | 0.04 | 0.04 | 0.94 |
| Whole_Blood | ENSG00000231346.1 | 312659 | 407 | 195 | 3.78E-01 | 0.04 | 0.05 | 0.94 |
| Brain_Amygdala | ENSG00000243960.1 | 482557 | 100 | 47 | 3.79E-01 | -0.18 | 0.20 | 0.94 |
| Brain_Cerebellum | ENSG00000227811.2 | 181541 | 173 | 96 | 3.79E-01 | 0.11 | 0.13 | 0.94 |
| Spleen | ENSG00000171385.5 | -67773 | 162 | 55 | 3.80E-01 | -0.10 | 0.12 | 0.94 |
| Brain_Cerebellum | ENSG00000224167.1 | -955558 | 173 | 96 | 3.80E-01 | 0.12 | 0.13 | 0.94 |
| Heart_Left_Ventricle | ENSG00000134245.13 | -545159 | 303 | 154 | 3.80E-01 | -0.05 | 0.06 | 0.94 |
| Esophagus_Mucosa | ENSG00000260948.1 | 488314 | 407 | 208 | 3.82E-01 | 0.05 | 0.06 | 0.94 |
| Brain_Spinal_cord_cervical_c-1 | ENSG00000143079.10 | -474799 | 91 | 49 | 3.83E-01 | 0.10 | 0.11 | 0.94 |
| Skin_Not_Sun_Exposed_Suprapubic | ENSG00000143110.7 | 447590 | 387 | 200 | 3.84E-01 | -0.04 | 0.04 | 0.94 |
| Thyroid | ENSG00000155363.14 | -751759 | 446 | 224 | 3.84E-01 | -0.02 | 0.03 | 0.94 |
| Pancreas | ENSG00000225075.1 | -775239 | 248 | 108 | 3.84E-01 | -0.08 | 0.09 | 0.94 |
| Pancreas | ENSG00000231437.3 | -68388 | 248 | 108 | 3.84E-01 | 0.06 | 0.07 | 0.94 |
| Adipose_Visceral_Omentum | ENSG00000116459.6 | 472490 | 355 | 172 | 3.84E-01 | 0.02 | 0.02 | 0.94 |
| Brain_Substantia_nigra | ENSG00000155367.11 | -794095 | 88 | 49 | 3.85E-01 | 0.07 | 0.07 | 0.94 |
| Brain_Cortex | ENSG00000064886.9 | 720611 | 158 | 79 | 3.85E-01 | -0.08 | 0.09 | 0.94 |
| Heart_Atrial_Appendage | ENSG00000155366.12 | -786052 | 297 | 147 | 3.85E-01 | -0.03 | 0.04 | 0.94 |
| Brain_Putamen_basal_ganglia | ENSG00000155363.14 | -751759 | 124 | 71 | 3.85E-01 | -0.06 | 0.07 | 0.94 |
| Thyroid | ENSG00000143110.7 | 447590 | 446 | 224 | 3.86E-01 | 0.02 | 0.03 | 0.94 |
| Esophagus_Muscularis | ENSG00000143079.10 | -474799 | 370 | 188 | 3.86E-01 | -0.05 | 0.05 | 0.94 |
| Cells_EBV-transformed_lymphocytes | ENSG00000225075.1 | -775239 | 130 | 68 | 3.86E-01 | 0.10 | 0.11 | 0.94 |
| Heart_Left_Ventricle | ENSG00000116489.8 | -698415 | 303 | 154 | 3.87E-01 | -0.03 | 0.03 | 0.94 |
| Nerve_Tibial | ENSG00000225075.1 | -775239 | 414 | 202 | 3.88E-01 | 0.07 | 0.08 | 0.94 |
| Cells_EBV-transformed_lymphocytes | ENSG00000064886.9 | 720611 | 130 | 68 | 3.89E-01 | -0.11 | 0.12 | 0.94 |

| | | | | | | | | |
|---|---|---|---|---|---|---|---|---|
| Colon_Sigmoid | ENSG00000134255.9 | 781156 | 233 | 107 | 3.89E-01 | -0.05 | 0.06 | 0.94 |
| Colon_Transverse | ENSG00000171385.5 | -67773 | 274 | 119 | 3.89E-01 | -0.04 | 0.04 | 0.94 |
| Adipose_Subcutaneous | ENSG00000116489.8 | -698415 | 442 | 214 | 3.89E-01 | -0.02 | 0.02 | 0.94 |
| Esophagus_Gastroesophageal_Junction | ENSG00000231246.1 | -439146 | 244 | 110 | 3.90E-01 | -0.07 | 0.08 | 0.94 |
| Vagina | ENSG00000173947.9 | 575094 | 115 | 59 | 3.90E-01 | 0.08 | 0.10 | 0.94 |
| Adipose_Visceral_Omentum | ENSG00000116455.9 | 472519 | 355 | 172 | 3.91E-01 | 0.02 | 0.02 | 0.94 |
| Esophagus_Gastroesophageal_Junction | ENSG00000173947.9 | 575094 | 244 | 110 | 3.92E-01 | -0.05 | 0.06 | 0.94 |
| Brain_Frontal_Cortex_BA9 | ENSG00000184599.9 | -799037 | 129 | 73 | 3.93E-01 | -0.12 | 0.14 | 0.94 |
| Brain_Amygdala | ENSG00000215866.3 | -929261 | 100 | 47 | 3.93E-01 | 0.11 | 0.13 | 0.94 |
| Minor_Salivary_Gland | ENSG00000243960.1 | 482557 | 97 | 49 | 3.93E-01 | 0.10 | 0.11 | 0.94 |
| Brain_Substantia_nigra | ENSG00000231246.1 | -439146 | 88 | 49 | 3.95E-01 | -0.13 | 0.15 | 0.94 |
| Adipose_Visceral_Omentum | ENSG00000143079.10 | -474799 | 355 | 172 | 3.95E-01 | -0.03 | 0.03 | 0.94 |
| Brain_Cerebellar_Hemisphere | ENSG00000007341.14 | -699443 | 136 | 74 | 3.95E-01 | -0.09 | 0.10 | 0.94 |
| Testis | ENSG00000227811.2 | 181541 | 259 | 128 | 3.96E-01 | -0.07 | 0.08 | 0.94 |
| Brain_Spinal_cord_cervical_c-1 | ENSG00000143110.7 | 447590 | 91 | 49 | 3.96E-01 | 0.07 | 0.08 | 0.94 |
| Brain_Cerebellar_Hemisphere | ENSG00000224167.1 | -955558 | 136 | 74 | 3.96E-01 | -0.10 | 0.12 | 0.94 |
| Testis | ENSG00000171385.5 | -67773 | 259 | 128 | 3.96E-01 | 0.07 | 0.08 | 0.94 |
| Brain_Spinal_cord_cervical_c-1 | ENSG00000261654.1 | 979029 | 91 | 49 | 3.97E-01 | -0.13 | 0.16 | 0.94 |
| Pancreas | ENSG00000231346.1 | 312659 | 248 | 108 | 3.97E-01 | 0.08 | 0.09 | 0.94 |
| Esophagus_Muscularis | ENSG00000197852.8 | 168128 | 370 | 188 | 3.97E-01 | -0.04 | 0.05 | 0.94 |
| Cells_EBV-transformed_lymphocytes | ENSG00000243960.1 | 482557 | 130 | 68 | 3.98E-01 | -0.11 | 0.13 | 0.94 |
| Artery_Aorta | ENSG00000121931.11 | 968421 | 299 | 152 | 3.98E-01 | -0.04 | 0.05 | 0.94 |
| Cells_Transformed_fibroblasts | ENSG00000143110.7 | 447590 | 343 | 175 | 3.99E-01 | 0.06 | 0.07 | 0.94 |
| Minor_Salivary_Gland | ENSG00000273010.1 | 958080 | 97 | 49 | 4.00E-01 | -0.12 | 0.14 | 0.94 |
| Brain_Hypothalamus | ENSG00000155367.11 | -794095 | 121 | 61 | 4.00E-01 | 0.07 | 0.09 | 0.94 |
| Muscle_Skeletal | ENSG00000134255.9 | 781156 | 564 | 276 | 4.01E-01 | 0.04 | 0.04 | 0.94 |
| Brain_Cerebellar_Hemisphere | ENSG00000197852.8 | 168128 | 136 | 74 | 4.01E-01 | -0.08 | 0.09 | 0.94 |
| Heart_Atrial_Appendage | ENSG00000260948.1 | 488314 | 297 | 147 | 4.01E-01 | 0.06 | 0.07 | 0.94 |
| Ovary | ENSG00000064886.9 | 720611 | 133 | 71 | 4.03E-01 | -0.08 | 0.09 | 0.94 |
| Artery_Aorta | ENSG00000155367.11 | -794095 | 299 | 152 | 4.03E-01 | -0.03 | 0.04 | 0.94 |
| Brain_Substantia_nigra | ENSG00000233337.1 | 483868 | 88 | 49 | 4.03E-01 | -0.13 | 0.15 | 0.94 |
| Artery_Coronary | ENSG00000116473.10 | 379164 | 173 | 81 | 4.04E-01 | 0.04 | 0.05 | 0.94 |
| Thyroid | ENSG00000261654.1 | 979029 | 446 | 224 | 4.05E-01 | 0.05 | 0.06 | 0.94 |
| Colon_Sigmoid | ENSG00000232811.1 | 977915 | 233 | 107 | 4.05E-01 | 0.09 | 0.11 | 0.94 |
| Esophagus_Mucosa | ENSG00000116455.9 | 472519 | 407 | 208 | 4.05E-01 | 0.03 | 0.03 | 0.94 |
| Liver | ENSG00000064886.9 | 720611 | 175 | 88 | 4.06E-01 | 0.10 | 0.12 | 0.94 |
| Breast_Mammary_Tissue | ENSG00000116489.8 | -698415 | 290 | 148 | 4.06E-01 | -0.03 | 0.03 | 0.94 |
| Small_Intestine_Terminal_Ileum | ENSG00000116489.8 | -698415 | 137 | 54 | 4.07E-01 | -0.06 | 0.07 | 0.94 |
| Testis | ENSG00000260948.1 | 488314 | 259 | 128 | 4.07E-01 | 0.05 | 0.07 | 0.94 |
| Cells_EBV-transformed_lymphocytes | ENSG00000143079.10 | -474799 | 130 | 68 | 4.07E-01 | -0.08 | 0.10 | 0.94 |
| Brain_Cerebellar_Hemisphere | ENSG00000064703.7 | 166137 | 136 | 74 | 4.07E-01 | -0.07 | 0.08 | 0.94 |
| Brain_Cerebellar_Hemisphere | ENSG00000121931.11 | 968421 | 136 | 74 | 4.08E-01 | 0.07 | 0.09 | 0.94 |
| Artery_Tibial | ENSG00000273483.1 | -597059 | 441 | 219 | 4.08E-01 | -0.05 | 0.07 | 0.94 |
| Spleen | ENSG00000197852.8 | 168128 | 162 | 55 | 4.08E-01 | -0.07 | 0.08 | 0.94 |
| Cells_EBV-transformed_lymphocytes | ENSG00000064703.7 | 166137 | 130 | 68 | 4.09E-01 | 0.05 | 0.07 | 0.94 |
| Prostate | ENSG00000116455.9 | 472519 | 152 | 75 | 4.10E-01 | -0.05 | 0.06 | 0.94 |
| Pituitary | ENSG00000232811.1 | 977915 | 183 | 95 | 4.11E-01 | 0.11 | 0.13 | 0.94 |

| | | | | | | | | |
|---|---|---|---|---|---|---|---|---|
| Brain_Cerebellum | ENSG00000171385.5 | -67773 | 173 | 96 | 4.11E-01 | 0.05 | 0.06 | 0.94 |
| Skin_Sun_Exposed_Lower_leg | ENSG00000197852.8 | 168128 | 473 | 233 | 4.11E-01 | 0.03 | 0.04 | 0.94 |
| Lung | ENSG00000261654.1 | 979029 | 427 | 222 | 4.12E-01 | 0.04 | 0.05 | 0.94 |
| Skin_Not_Sun_Exposed_Suprapubic | ENSG00000231246.1 | -439146 | 387 | 200 | 4.13E-01 | 0.05 | 0.07 | 0.94 |
| Brain_Amygdala | ENSG00000155363.14 | -751759 | 100 | 47 | 4.14E-01 | -0.08 | 0.10 | 0.94 |
| Breast_Mammary_Tissue | ENSG00000116459.6 | 472490 | 290 | 148 | 4.14E-01 | -0.02 | 0.02 | 0.94 |
| Nerve_Tibial | ENSG00000260948.1 | 488314 | 414 | 202 | 4.15E-01 | 0.04 | 0.05 | 0.94 |
| Small_Intestine_Terminal_Ileum | ENSG00000171385.5 | -67773 | 137 | 54 | 4.15E-01 | 0.07 | 0.08 | 0.94 |
| Vagina | ENSG00000233337.1 | 483868 | 115 | 59 | 4.15E-01 | 0.11 | 0.14 | 0.94 |
| Brain_Cerebellum | ENSG00000121931.11 | 968421 | 173 | 96 | 4.17E-01 | 0.06 | 0.07 | 0.94 |
| Muscle_Skeletal | ENSG00000085465.11 | 493605 | 564 | 276 | 4.18E-01 | 0.03 | 0.03 | 0.94 |
| Brain_Putamen_basal_ganglia | ENSG00000134245.13 | -545159 | 124 | 71 | 4.18E-01 | 0.04 | 0.05 | 0.94 |
| Brain_Putamen_basal_ganglia | ENSG00000085465.11 | 493605 | 124 | 71 | 4.18E-01 | 0.06 | 0.07 | 0.94 |
| Adipose_Subcutaneous | ENSG00000225075.1 | -775239 | 442 | 214 | 4.23E-01 | -0.06 | 0.07 | 0.94 |
| Esophagus_Mucosa | ENSG00000232811.1 | 977915 | 407 | 208 | 4.24E-01 | -0.06 | 0.08 | 0.94 |
| Skin_Not_Sun_Exposed_Suprapubic | ENSG00000155363.14 | -751759 | 387 | 200 | 4.25E-01 | 0.02 | 0.03 | 0.94 |
| Brain_Cerebellum | ENSG00000156171.10 | 781166 | 173 | 96 | 4.26E-01 | 0.07 | 0.09 | 0.94 |
| Spleen | ENSG00000116489.8 | -698415 | 162 | 55 | 4.26E-01 | 0.05 | 0.07 | 0.94 |
| Nerve_Tibial | ENSG00000173947.9 | 575094 | 414 | 202 | 4.26E-01 | 0.04 | 0.05 | 0.94 |
| Prostate | ENSG00000273483.1 | -597059 | 152 | 75 | 4.26E-01 | -0.06 | 0.08 | 0.94 |
| Whole_Blood | ENSG00000134245.13 | -545159 | 407 | 195 | 4.27E-01 | -0.05 | 0.06 | 0.94 |
| Brain_Anterior_cingulate_cortex_BA24 | ENSG00000184599.9 | -799037 | 121 | 65 | 4.27E-01 | 0.10 | 0.12 | 0.94 |
| Brain_Substantia_nigra | ENSG00000171385.5 | -67773 | 88 | 49 | 4.27E-01 | 0.05 | 0.06 | 0.94 |
| Brain_Cerebellar_Hemisphere | ENSG00000134255.9 | 781156 | 136 | 74 | 4.28E-01 | -0.05 | 0.06 | 0.94 |
| Brain_Hypothalamus | ENSG00000064886.9 | 720611 | 121 | 61 | 4.28E-01 | 0.08 | 0.10 | 0.94 |
| Heart_Atrial_Appendage | ENSG00000064886.9 | 720611 | 297 | 147 | 4.29E-01 | -0.05 | 0.07 | 0.94 |
| Adipose_Visceral_Omentum | ENSG00000007341.14 | -699443 | 355 | 172 | 4.29E-01 | -0.05 | 0.07 | 0.94 |
| Brain_Substantia_nigra | ENSG00000116489.8 | -698415 | 88 | 49 | 4.30E-01 | 0.07 | 0.09 | 0.94 |
| Stomach | ENSG00000155363.14 | -751759 | 262 | 107 | 4.31E-01 | -0.04 | 0.05 | 0.94 |
| Adipose_Subcutaneous | ENSG00000173947.9 | 575094 | 442 | 214 | 4.31E-01 | -0.04 | 0.05 | 0.94 |
| Brain_Anterior_cingulate_cortex_BA24 | ENSG00000273483.1 | -597059 | 121 | 65 | 4.31E-01 | 0.07 | 0.08 | 0.94 |
| Brain_Cortex | ENSG00000171385.5 | -67773 | 158 | 79 | 4.32E-01 | -0.05 | 0.06 | 0.94 |
| Cells_Transformed_fibroblasts | ENSG00000116459.6 | 472490 | 343 | 175 | 4.32E-01 | -0.01 | 0.02 | 0.94 |
| Minor_Salivary_Gland | ENSG00000116489.8 | -698415 | 97 | 49 | 4.33E-01 | -0.05 | 0.06 | 0.94 |
| Brain_Hypothalamus | ENSG00000171385.5 | -67773 | 121 | 61 | 4.33E-01 | 0.07 | 0.09 | 0.94 |
| Brain_Cortex | ENSG00000143110.7 | 447590 | 158 | 79 | 4.34E-01 | 0.06 | 0.07 | 0.94 |
| Skin_Sun_Exposed_Lower_leg | ENSG00000232811.1 | 977915 | 473 | 233 | 4.35E-01 | 0.06 | 0.07 | 0.95 |
| Stomach | ENSG00000116489.8 | -698415 | 262 | 107 | 4.36E-01 | -0.03 | 0.04 | 0.95 |
| Small_Intestine_Terminal_Ileum | ENSG00000225075.1 | -775239 | 137 | 54 | 4.37E-01 | 0.07 | 0.10 | 0.95 |
| Colon_Sigmoid | ENSG00000134245.13 | -545159 | 233 | 107 | 4.38E-01 | -0.05 | 0.06 | 0.95 |
| Spleen | ENSG00000007341.14 | -699443 | 162 | 55 | 4.39E-01 | 0.10 | 0.12 | 0.95 |
| Brain_Nucleus_accumbens_basal_ganglia | ENSG00000215866.3 | -929261 | 147 | 77 | 4.41E-01 | 0.09 | 0.11 | 0.95 |
| Brain_Cerebellum | ENSG00000273483.1 | -597059 | 173 | 96 | 4.43E-01 | 0.06 | 0.08 | 0.95 |
| Brain_Amygdala | ENSG00000233337.1 | 483868 | 100 | 47 | 4.43E-01 | -0.15 | 0.20 | 0.95 |
| Cells_Transformed_fibroblasts | ENSG00000231246.1 | -439146 | 343 | 175 | 4.44E-01 | -0.04 | 0.06 | 0.95 |
| Skin_Sun_Exposed_Lower_leg | ENSG00000143110.7 | 447590 | 473 | 233 | 4.45E-01 | -0.02 | 0.03 | 0.95 |
| Skin_Sun_Exposed_Lower_leg | ENSG00000243960.1 | 482557 | 473 | 233 | 4.45E-01 | 0.05 | 0.07 | 0.95 |

| | | | | | | | | |
|---|---|---|---|---|---|---|---|---|
| Brain_Amygdala | ENSG00000155367.11 | -794095 | 100 | 47 | 4.45E-01 | 0.11 | 0.15 | 0.95 |
| Brain_Amygdala | ENSG00000156171.10 | 781166 | 100 | 47 | 4.46E-01 | -0.09 | 0.12 | 0.95 |
| Brain_Nucleus_accumbens_basal_ganglia | ENSG00000007341.14 | -699443 | 147 | 77 | 4.47E-01 | -0.10 | 0.13 | 0.95 |
| Brain_Hypothalamus | ENSG00000064703.7 | 166137 | 121 | 61 | 4.47E-01 | -0.08 | 0.11 | 0.95 |
| Colon_Transverse | ENSG00000273483.1 | -597059 | 274 | 119 | 4.47E-01 | 0.05 | 0.07 | 0.95 |
| Stomach | ENSG00000085465.11 | 493605 | 262 | 107 | 4.49E-01 | -0.05 | 0.06 | 0.95 |
| Adipose_Subcutaneous | ENSG00000007341.14 | -699443 | 442 | 214 | 4.51E-01 | -0.05 | 0.06 | 0.95 |
| Small_Intestine_Terminal_Ileum | ENSG00000064886.9 | 720611 | 137 | 54 | 4.51E-01 | 0.06 | 0.08 | 0.95 |
| Brain_Amygdala | ENSG00000197852.8 | 168128 | 100 | 47 | 4.52E-01 | 0.07 | 0.09 | 0.95 |
| Brain_Spinal_cord_cervical_c-1 | ENSG00000156171.10 | 781166 | 91 | 49 | 4.52E-01 | -0.08 | 0.11 | 0.95 |
| Stomach | ENSG00000260948.1 | 488314 | 262 | 107 | 4.52E-01 | -0.06 | 0.08 | 0.95 |
| Brain_Anterior_cingulate_cortex_BA24 | ENSG00000155363.14 | -751759 | 121 | 65 | 4.52E-01 | -0.04 | 0.05 | 0.95 |
| Small_Intestine_Terminal_Ileum | ENSG00000162777.12 | 716847 | 137 | 54 | 4.53E-01 | -0.04 | 0.05 | 0.95 |
| Esophagus_Mucosa | ENSG00000064703.7 | 166137 | 407 | 208 | 4.53E-01 | 0.03 | 0.03 | 0.95 |
| Brain_Putamen_basal_ganglia | ENSG00000064886.9 | 720611 | 124 | 71 | 4.53E-01 | -0.09 | 0.12 | 0.95 |
| Brain_Cerebellum | ENSG00000261654.1 | 979029 | 173 | 96 | 4.54E-01 | 0.07 | 0.10 | 0.95 |
| Adipose_Visceral_Omentum | ENSG00000143110.7 | 447590 | 355 | 172 | 4.55E-01 | -0.02 | 0.03 | 0.95 |
| Brain_Caudate_basal_ganglia | ENSG00000085465.11 | 493605 | 160 | 88 | 4.55E-01 | 0.05 | 0.06 | 0.95 |
| Liver | ENSG00000143079.10 | -474799 | 175 | 88 | 4.55E-01 | -0.05 | 0.06 | 0.95 |
| Vagina | ENSG00000143079.10 | -474799 | 115 | 59 | 4.56E-01 | 0.05 | 0.07 | 0.95 |
| Brain_Cerebellar_Hemisphere | ENSG00000121933.13 | 357420 | 136 | 74 | 4.56E-01 | -0.08 | 0.11 | 0.95 |
| Testis | ENSG00000143110.7 | 447590 | 259 | 128 | 4.57E-01 | 0.06 | 0.07 | 0.95 |
| Testis | ENSG00000116473.10 | 379164 | 259 | 128 | 4.57E-01 | -0.02 | 0.02 | 0.95 |
| Muscle_Skeletal | ENSG00000134245.13 | -545159 | 564 | 276 | 4.59E-01 | 0.03 | 0.05 | 0.95 |
| Adrenal_Gland | ENSG00000233337.1 | 483868 | 190 | 96 | 4.60E-01 | -0.07 | 0.09 | 0.95 |
| Brain_Frontal_Cortex_BA9 | ENSG00000273010.1 | 958080 | 129 | 73 | 4.61E-01 | -0.10 | 0.13 | 0.95 |
| Adrenal_Gland | ENSG00000184599.9 | -799037 | 190 | 96 | 4.62E-01 | 0.04 | 0.05 | 0.95 |
| Vagina | ENSG00000238975.1 | -731206 | 115 | 59 | 4.62E-01 | 0.11 | 0.15 | 0.95 |
| Heart_Left_Ventricle | ENSG00000121933.13 | 357420 | 303 | 154 | 4.63E-01 | -0.04 | 0.05 | 0.95 |
| Adipose_Visceral_Omentum | ENSG00000260948.1 | 488314 | 355 | 172 | 4.64E-01 | 0.05 | 0.06 | 0.95 |
| Lung | ENSG00000134216.14 | 630520 | 427 | 222 | 4.64E-01 | 0.04 | 0.05 | 0.95 |
| Muscle_Skeletal | ENSG00000116459.6 | 472490 | 564 | 276 | 4.65E-01 | -0.01 | 0.02 | 0.95 |
| Colon_Sigmoid | ENSG00000233337.1 | 483868 | 233 | 107 | 4.65E-01 | -0.07 | 0.10 | 0.95 |
| Pancreas | ENSG00000171385.5 | -67773 | 248 | 108 | 4.65E-01 | 0.04 | 0.05 | 0.95 |
| Breast_Mammary_Tissue | ENSG00000273483.1 | -597059 | 290 | 148 | 4.66E-01 | -0.06 | 0.09 | 0.95 |
| Spleen | ENSG00000231437.3 | -68388 | 162 | 55 | 4.67E-01 | -0.08 | 0.12 | 0.95 |
| Vagina | ENSG00000227811.2 | 181541 | 115 | 59 | 4.67E-01 | -0.12 | 0.16 | 0.95 |
| Adipose_Visceral_Omentum | ENSG00000121933.13 | 357420 | 355 | 172 | 4.67E-01 | -0.02 | 0.03 | 0.95 |
| Brain_Hippocampus | ENSG00000143110.7 | 447590 | 123 | 63 | 4.67E-01 | 0.05 | 0.06 | 0.95 |
| Brain_Spinal_cord_cervical_c-1 | ENSG00000121933.13 | 357420 | 91 | 49 | 4.69E-01 | 0.07 | 0.10 | 0.95 |
| Liver | ENSG00000162777.12 | 716847 | 175 | 88 | 4.69E-01 | -0.08 | 0.11 | 0.95 |
| Lung | ENSG00000225075.1 | -775239 | 427 | 222 | 4.70E-01 | 0.05 | 0.06 | 0.95 |
| Colon_Transverse | ENSG00000116455.9 | 472519 | 274 | 119 | 4.70E-01 | 0.03 | 0.04 | 0.95 |
| Spleen | ENSG00000231246.1 | -439146 | 162 | 55 | 4.72E-01 | 0.09 | 0.13 | 0.95 |
| Colon_Transverse | ENSG00000173947.9 | 575094 | 274 | 119 | 4.72E-01 | -0.03 | 0.04 | 0.95 |
| Muscle_Skeletal | ENSG00000171385.5 | -67773 | 564 | 276 | 4.73E-01 | -0.03 | 0.04 | 0.95 |
| Skin_Not_Sun_Exposed_Suprapubic | ENSG00000273483.1 | -597059 | 387 | 200 | 4.73E-01 | 0.06 | 0.08 | 0.95 |

| | | | | | | | | |
|---|---|---|---|---|---|---|---|---|
| Colon_Sigmoid | ENSG00000273483.1 | -597059 | 233 | 107 | 4.73E-01 | 0.04 | 0.06 | 0.95 |
| Brain_Cerebellum | ENSG00000184599.9 | -799037 | 173 | 96 | 4.74E-01 | 0.07 | 0.10 | 0.95 |
| Brain_Nucleus_accumbens_basal_ganglia | ENSG00000156171.10 | 781166 | 147 | 77 | 4.74E-01 | -0.08 | 0.11 | 0.95 |
| Brain_Hippocampus | ENSG00000227811.2 | 181541 | 123 | 63 | 4.76E-01 | -0.09 | 0.13 | 0.95 |
| Cells_Transformed_fibroblasts | ENSG00000273483.1 | -597059 | 343 | 175 | 4.77E-01 | -0.05 | 0.07 | 0.95 |
| Esophagus_Gastroesophageal_Junction | ENSG00000155367.11 | -794095 | 244 | 110 | 4.77E-01 | -0.05 | 0.07 | 0.95 |
| Skin_Sun_Exposed_Lower_leg | ENSG00000155366.12 | -786052 | 473 | 233 | 4.78E-01 | 0.02 | 0.03 | 0.95 |
| Small_Intestine_Terminal_Ileum | ENSG00000232811.1 | 977915 | 137 | 54 | 4.78E-01 | -0.09 | 0.12 | 0.95 |
| Prostate | ENSG00000225075.1 | -775239 | 152 | 75 | 4.78E-01 | -0.06 | 0.09 | 0.95 |
| Spleen | ENSG00000155363.14 | -751759 | 162 | 55 | 4.78E-01 | -0.05 | 0.07 | 0.95 |
| Brain_Spinal_cord_cervical_c-1 | ENSG00000273483.1 | -597059 | 91 | 49 | 4.78E-01 | -0.11 | 0.15 | 0.95 |
| Small_Intestine_Terminal_Ileum | ENSG00000143079.10 | -474799 | 137 | 54 | 4.79E-01 | -0.05 | 0.07 | 0.95 |
| Liver | ENSG00000155367.11 | -794095 | 175 | 88 | 4.80E-01 | 0.07 | 0.11 | 0.95 |
| Cells_Transformed_fibroblasts | ENSG00000156171.10 | 781166 | 343 | 175 | 4.80E-01 | 0.02 | 0.03 | 0.95 |
| Colon_Transverse | ENSG00000261654.1 | 979029 | 274 | 119 | 4.82E-01 | -0.06 | 0.08 | 0.95 |
| Artery_Aorta | ENSG00000085465.11 | 493605 | 299 | 152 | 4.82E-01 | -0.03 | 0.04 | 0.95 |
| Brain_Hippocampus | ENSG00000116455.9 | 472519 | 123 | 63 | 4.82E-01 | -0.06 | 0.08 | 0.95 |
| Artery_Aorta | ENSG00000116459.6 | 472490 | 299 | 152 | 4.84E-01 | 0.02 | 0.03 | 0.95 |
| Liver | ENSG00000007341.14 | -699443 | 175 | 88 | 4.84E-01 | -0.09 | 0.13 | 0.95 |
| Brain_Cerebellar_Hemisphere | ENSG00000162777.12 | 716847 | 136 | 74 | 4.84E-01 | 0.07 | 0.10 | 0.95 |
| Vagina | ENSG00000064886.9 | 720611 | 115 | 59 | 4.84E-01 | -0.09 | 0.13 | 0.95 |
| Spleen | ENSG00000225075.1 | -775239 | 162 | 55 | 4.85E-01 | 0.08 | 0.12 | 0.95 |
| Brain_Hippocampus | ENSG00000156171.10 | 781166 | 123 | 63 | 4.85E-01 | -0.06 | 0.09 | 0.95 |
| Brain_Spinal_cord_cervical_c-1 | ENSG00000197852.8 | 168128 | 91 | 49 | 4.85E-01 | -0.11 | 0.16 | 0.95 |
| Brain_Substantia_nigra | ENSG00000197852.8 | 168128 | 88 | 49 | 4.86E-01 | 0.09 | 0.13 | 0.95 |
| Brain_Frontal_Cortex_BA9 | ENSG00000173947.9 | 575094 | 129 | 73 | 4.86E-01 | 0.06 | 0.09 | 0.95 |
| Esophagus_Gastroesophageal_Junction | ENSG00000232811.1 | 977915 | 244 | 110 | 4.87E-01 | 0.08 | 0.11 | 0.95 |
| Muscle_Skeletal | ENSG00000064886.9 | 720611 | 564 | 276 | 4.88E-01 | 0.03 | 0.05 | 0.95 |
| Stomach | ENSG00000116473.10 | 379164 | 262 | 107 | 4.89E-01 | -0.03 | 0.04 | 0.95 |
| Brain_Frontal_Cortex_BA9 | ENSG00000215866.3 | -929261 | 129 | 73 | 4.89E-01 | 0.11 | 0.15 | 0.95 |
| Colon_Transverse | ENSG00000064703.7 | 166137 | 274 | 119 | 4.89E-01 | 0.04 | 0.06 | 0.95 |
| Adipose_Subcutaneous | ENSG00000155367.11 | -794095 | 442 | 214 | 4.90E-01 | 0.03 | 0.04 | 0.95 |
| Nerve_Tibial | ENSG00000156171.10 | 781166 | 414 | 202 | 4.91E-01 | 0.04 | 0.05 | 0.95 |
| Vagina | ENSG00000134245.13 | -545159 | 115 | 59 | 4.91E-01 | 0.07 | 0.11 | 0.95 |
| Esophagus_Muscularis | ENSG00000143110.7 | 447590 | 370 | 188 | 4.93E-01 | -0.03 | 0.04 | 0.95 |
| Cells_Transformed_fibroblasts | ENSG00000233337.1 | 483868 | 343 | 175 | 4.94E-01 | -0.04 | 0.05 | 0.95 |
| Lung | ENSG00000231246.1 | -439146 | 427 | 222 | 4.94E-01 | -0.04 | 0.06 | 0.95 |
| Pituitary | ENSG00000121933.13 | 357420 | 183 | 95 | 4.95E-01 | 0.05 | 0.07 | 0.95 |
| Minor_Salivary_Gland | ENSG00000238975.1 | -731206 | 97 | 49 | 4.95E-01 | -0.10 | 0.14 | 0.95 |
| Brain_Cerebellar_Hemisphere | ENSG00000232811.1 | 977915 | 136 | 74 | 4.96E-01 | 0.09 | 0.14 | 0.95 |
| Vagina | ENSG00000085465.11 | 493605 | 115 | 59 | 4.96E-01 | 0.07 | 0.11 | 0.95 |
| Adipose_Subcutaneous | ENSG00000273010.1 | 958080 | 442 | 214 | 4.97E-01 | -0.04 | 0.07 | 0.95 |
| Colon_Sigmoid | ENSG00000197852.8 | 168128 | 233 | 107 | 4.97E-01 | 0.05 | 0.07 | 0.95 |
| Brain_Cortex | ENSG00000162777.12 | 716847 | 158 | 79 | 4.97E-01 | -0.08 | 0.11 | 0.95 |
| Adrenal_Gland | ENSG00000116455.9 | 472519 | 190 | 96 | 4.97E-01 | 0.04 | 0.06 | 0.95 |
| Brain_Frontal_Cortex_BA9 | ENSG00000232811.1 | 977915 | 129 | 73 | 4.99E-01 | -0.09 | 0.13 | 0.95 |
| Esophagus_Mucosa | ENSG00000121931.11 | 968421 | 407 | 208 | 4.99E-01 | -0.03 | 0.05 | 0.95 |

| | | | | | | | | |
|---|---|---|---|---|---|---|---|---|
| Stomach | ENSG00000155367.11 | -794095 | 262 | 107 | 5.01E-01 | 0.06 | 0.09 | 0.95 |
| Brain_Cortex | ENSG00000231346.1 | 312659 | 158 | 79 | 5.02E-01 | -0.09 | 0.14 | 0.95 |
| Thyroid | ENSG00000134245.13 | -545159 | 446 | 224 | 5.03E-01 | -0.03 | 0.04 | 0.95 |
| Brain_Substantia_nigra | ENSG00000273010.1 | 958080 | 88 | 49 | 5.04E-01 | -0.11 | 0.16 | 0.95 |
| Uterus | ENSG00000243960.1 | 482557 | 111 | 63 | 5.04E-01 | -0.09 | 0.14 | 0.95 |
| Minor_Salivary_Gland | ENSG00000143079.10 | -474799 | 97 | 49 | 5.05E-01 | -0.03 | 0.05 | 0.95 |
| Colon_Sigmoid | ENSG00000155367.11 | -794095 | 233 | 107 | 5.06E-01 | 0.04 | 0.07 | 0.95 |
| Muscle_Skeletal | ENSG00000156171.10 | 781166 | 564 | 276 | 5.06E-01 | -0.03 | 0.05 | 0.95 |
| Pituitary | ENSG00000155366.12 | -786052 | 183 | 95 | 5.06E-01 | 0.04 | 0.05 | 0.95 |
| Pituitary | ENSG00000225075.1 | -775239 | 183 | 95 | 5.07E-01 | -0.08 | 0.13 | 0.95 |
| Uterus | ENSG00000143079.10 | -474799 | 111 | 63 | 5.07E-01 | -0.05 | 0.07 | 0.95 |
| Vagina | ENSG00000243960.1 | 482557 | 115 | 59 | 5.07E-01 | 0.09 | 0.14 | 0.95 |
| Nerve_Tibial | ENSG00000227811.2 | 181541 | 414 | 202 | 5.07E-01 | 0.05 | 0.08 | 0.95 |
| Pancreas | ENSG00000134255.9 | 781156 | 248 | 108 | 5.08E-01 | -0.05 | 0.07 | 0.95 |
| Pancreas | ENSG00000085465.11 | 493605 | 248 | 108 | 5.09E-01 | 0.05 | 0.07 | 0.95 |
| Cells_EBV-transformed_lymphocytes | ENSG00000156171.10 | 781166 | 130 | 68 | 5.10E-01 | 0.04 | 0.06 | 0.95 |
| Brain_Spinal_cord_cervical_c-1 | ENSG00000243960.1 | 482557 | 91 | 49 | 5.12E-01 | -0.12 | 0.18 | 0.95 |
| Esophagus_Gastroesophageal_Junction | ENSG00000171385.5 | -67773 | 244 | 110 | 5.13E-01 | -0.03 | 0.04 | 0.95 |
| Brain_Cerebellum | ENSG00000064886.9 | 720611 | 173 | 96 | 5.14E-01 | -0.06 | 0.10 | 0.95 |
| Brain_Caudate_basal_ganglia | ENSG00000162777.12 | 716847 | 160 | 88 | 5.14E-01 | 0.06 | 0.09 | 0.95 |
| Cells_EBV-transformed_lymphocytes | ENSG00000121931.11 | 968421 | 130 | 68 | 5.15E-01 | -0.06 | 0.09 | 0.95 |
| Brain_Amygdala | ENSG00000173947.9 | 575094 | 100 | 47 | 5.15E-01 | 0.07 | 0.10 | 0.95 |
| Cells_EBV-transformed_lymphocytes | ENSG00000134255.9 | 781156 | 130 | 68 | 5.16E-01 | 0.06 | 0.10 | 0.95 |
| Cells_Transformed_fibroblasts | ENSG00000085465.11 | 493605 | 343 | 175 | 5.16E-01 | 0.02 | 0.03 | 0.95 |
| Brain_Anterior_cingulate_cortex_BA24 | ENSG00000162777.12 | 716847 | 121 | 65 | 5.17E-01 | 0.07 | 0.10 | 0.95 |
| Brain_Anterior_cingulate_cortex_BA24 | ENSG00000155366.12 | -786052 | 121 | 65 | 5.17E-01 | -0.02 | 0.04 | 0.95 |
| Lung | ENSG00000116473.10 | 379164 | 427 | 222 | 5.17E-01 | 0.02 | 0.03 | 0.95 |
| Brain_Cerebellum | ENSG00000237556.1 | 12046 | 173 | 96 | 5.17E-01 | -0.08 | 0.12 | 0.95 |
| Brain_Frontal_Cortex_BA9 | ENSG00000143079.10 | -474799 | 129 | 73 | 5.18E-01 | 0.05 | 0.08 | 0.95 |
| Brain_Frontal_Cortex_BA9 | ENSG00000243960.1 | 482557 | 129 | 73 | 5.18E-01 | 0.08 | 0.13 | 0.95 |
| Small_Intestine_Terminal_Ileum | ENSG00000134255.9 | 781156 | 137 | 54 | 5.19E-01 | -0.03 | 0.05 | 0.95 |
| Testis | ENSG00000155366.12 | -786052 | 259 | 128 | 5.19E-01 | -0.02 | 0.03 | 0.95 |
| Brain_Frontal_Cortex_BA9 | ENSG00000155367.11 | -794095 | 129 | 73 | 5.19E-01 | 0.05 | 0.07 | 0.95 |
| Testis | ENSG00000085465.11 | 493605 | 259 | 128 | 5.20E-01 | 0.03 | 0.04 | 0.95 |
| Brain_Amygdala | ENSG00000064886.9 | 720611 | 100 | 47 | 5.21E-01 | -0.08 | 0.13 | 0.95 |
| Artery_Aorta | ENSG00000232811.1 | 977915 | 299 | 152 | 5.22E-01 | -0.06 | 0.09 | 0.95 |
| Spleen | ENSG00000121933.13 | 357420 | 162 | 55 | 5.22E-01 | 0.06 | 0.10 | 0.95 |
| Artery_Coronary | ENSG00000162777.12 | 716847 | 173 | 81 | 5.22E-01 | -0.03 | 0.05 | 0.95 |
| Artery_Coronary | ENSG00000134245.13 | -545159 | 173 | 81 | 5.22E-01 | 0.04 | 0.07 | 0.95 |
| Prostate | ENSG00000121933.13 | 357420 | 152 | 75 | 5.23E-01 | -0.05 | 0.07 | 0.95 |
| Brain_Cerebellum | ENSG00000273010.1 | 958080 | 173 | 96 | 5.23E-01 | -0.07 | 0.12 | 0.95 |
| Heart_Left_Ventricle | ENSG00000064886.9 | 720611 | 303 | 154 | 5.24E-01 | 0.05 | 0.07 | 0.95 |
| Spleen | ENSG00000173947.9 | 575094 | 162 | 55 | 5.24E-01 | -0.10 | 0.16 | 0.95 |
| Brain_Putamen_basal_ganglia | ENSG00000260948.1 | 488314 | 124 | 71 | 5.24E-01 | -0.09 | 0.13 | 0.95 |
| Spleen | ENSG00000215866.3 | -929261 | 162 | 55 | 5.24E-01 | -0.09 | 0.14 | 0.95 |
| Ovary | ENSG00000155366.12 | -786052 | 133 | 71 | 5.24E-01 | -0.05 | 0.08 | 0.95 |
| Whole_Blood | ENSG00000232811.1 | 977915 | 407 | 195 | 5.25E-01 | 0.03 | 0.05 | 0.95 |

| | | | | | | | | |
|---|---|---|---|---|---|---|---|---|
| Small_Intestine_Terminal_Ileum | ENSG00000134245.13 | -545159 | 137 | 54 | 5.26E-01 | -0.05 | 0.08 | 0.95 |
| Vagina | ENSG00000007341.14 | -699443 | 115 | 59 | 5.27E-01 | -0.07 | 0.11 | 0.95 |
| Brain_Cortex | ENSG00000155366.12 | -786052 | 158 | 79 | 5.28E-01 | -0.03 | 0.04 | 0.95 |
| Brain_Putamen_basal_ganglia | ENSG00000143110.7 | 447590 | 124 | 71 | 5.28E-01 | -0.06 | 0.09 | 0.95 |
| Brain_Nucleus_accumbens_basal_ganglia | ENSG00000134255.9 | 781156 | 147 | 77 | 5.28E-01 | 0.03 | 0.06 | 0.95 |
| Cells_EBV-transformed_lymphocytes | ENSG00000273483.1 | -597059 | 130 | 68 | 5.30E-01 | -0.08 | 0.12 | 0.95 |
| Cells_EBV-transformed_lymphocytes | ENSG00000116455.9 | 472519 | 130 | 68 | 5.30E-01 | 0.03 | 0.05 | 0.95 |
| Esophagus_Gastroesophageal_Junction | ENSG00000064886.9 | 720611 | 244 | 110 | 5.30E-01 | -0.04 | 0.07 | 0.95 |
| Esophagus_Mucosa | ENSG00000116473.10 | 379164 | 407 | 208 | 5.32E-01 | -0.02 | 0.03 | 0.95 |
| Skin_Not_Sun_Exposed_Suprapubic | ENSG00000215866.3 | -929261 | 387 | 200 | 5.32E-01 | -0.05 | 0.08 | 0.95 |
| Brain_Spinal_cord_cervical_c-1 | ENSG00000155366.12 | -786052 | 91 | 49 | 5.33E-01 | 0.06 | 0.09 | 0.95 |
| Pancreas | ENSG00000260948.1 | 488314 | 248 | 108 | 5.33E-01 | -0.06 | 0.09 | 0.95 |
| Muscle_Skeletal | ENSG00000273483.1 | -597059 | 564 | 276 | 5.33E-01 | 0.03 | 0.05 | 0.95 |
| Cells_Transformed_fibroblasts | ENSG00000232811.1 | 977915 | 343 | 175 | 5.33E-01 | -0.03 | 0.05 | 0.95 |
| Whole_Blood | ENSG00000064703.7 | 166137 | 407 | 195 | 5.33E-01 | 0.03 | 0.05 | 0.95 |
| Prostate | ENSG00000121931.11 | 968421 | 152 | 75 | 5.34E-01 | 0.07 | 0.10 | 0.95 |
| Heart_Atrial_Appendage | ENSG00000121933.13 | 357420 | 297 | 147 | 5.35E-01 | 0.03 | 0.04 | 0.95 |
| Brain_Nucleus_accumbens_basal_ganglia | ENSG00000155367.11 | -794095 | 147 | 77 | 5.36E-01 | 0.05 | 0.09 | 0.95 |
| Brain_Hippocampus | ENSG00000085465.11 | 493605 | 123 | 63 | 5.36E-01 | -0.05 | 0.08 | 0.95 |
| Nerve_Tibial | ENSG00000162777.12 | 716847 | 414 | 202 | 5.37E-01 | -0.02 | 0.04 | 0.95 |
| Spleen | ENSG00000155366.12 | -786052 | 162 | 55 | 5.37E-01 | 0.04 | 0.07 | 0.95 |
| Lung | ENSG00000134245.13 | -545159 | 427 | 222 | 5.37E-01 | -0.02 | 0.04 | 0.95 |
| Esophagus_Muscularis | ENSG00000260948.1 | 488314 | 370 | 188 | 5.37E-01 | 0.04 | 0.07 | 0.95 |
| Brain_Cerebellum | ENSG00000155367.11 | -794095 | 173 | 96 | 5.38E-01 | 0.05 | 0.08 | 0.95 |
| Colon_Sigmoid | ENSG00000215867.4 | 271838 | 233 | 107 | 5.38E-01 | -0.06 | 0.10 | 0.95 |
| Pituitary | ENSG00000273483.1 | -597059 | 183 | 95 | 5.40E-01 | 0.05 | 0.08 | 0.96 |
| Brain_Hypothalamus | ENSG00000007341.14 | -699443 | 121 | 61 | 5.41E-01 | -0.07 | 0.12 | 0.96 |
| Breast_Mammary_Tissue | ENSG00000225075.1 | -775239 | 290 | 148 | 5.42E-01 | 0.06 | 0.09 | 0.96 |
| Brain_Spinal_cord_cervical_c-1 | ENSG00000233337.1 | 483868 | 91 | 49 | 5.43E-01 | -0.11 | 0.18 | 0.96 |
| Esophagus_Mucosa | ENSG00000116489.8 | -698415 | 407 | 208 | 5.43E-01 | -0.02 | 0.03 | 0.96 |
| Brain_Substantia_nigra | ENSG00000231437.3 | -68388 | 88 | 49 | 5.44E-01 | 0.05 | 0.08 | 0.96 |
| Brain_Anterior_cingulate_cortex_BA24 | ENSG00000007341.14 | -699443 | 121 | 65 | 5.44E-01 | -0.08 | 0.13 | 0.96 |
| Brain_Spinal_cord_cervical_c-1 | ENSG00000085465.11 | 493605 | 91 | 49 | 5.44E-01 | 0.05 | 0.08 | 0.96 |
| Esophagus_Mucosa | ENSG00000197852.8 | 168128 | 407 | 208 | 5.44E-01 | -0.03 | 0.05 | 0.96 |
| Ovary | ENSG00000134255.9 | 781156 | 133 | 71 | 5.46E-01 | 0.06 | 0.10 | 0.96 |
| Pituitary | ENSG00000184599.9 | -799037 | 183 | 95 | 5.46E-01 | 0.07 | 0.12 | 0.96 |
| Skin_Not_Sun_Exposed_Suprapubic | ENSG00000273010.1 | 958080 | 387 | 200 | 5.47E-01 | 0.05 | 0.08 | 0.96 |
| Vagina | ENSG00000197852.8 | 168128 | 115 | 59 | 5.47E-01 | -0.05 | 0.09 | 0.96 |
| Colon_Sigmoid | ENSG00000143110.7 | 447590 | 233 | 107 | 5.48E-01 | -0.04 | 0.06 | 0.96 |
| Brain_Cerebellar_Hemisphere | ENSG00000155363.14 | -751759 | 136 | 74 | 5.49E-01 | -0.04 | 0.07 | 0.96 |
| Artery_Coronary | ENSG00000231346.1 | 312659 | 173 | 81 | 5.50E-01 | -0.05 | 0.08 | 0.96 |
| Spleen | ENSG00000143110.7 | 447590 | 162 | 55 | 5.50E-01 | -0.05 | 0.09 | 0.96 |
| Artery_Aorta | ENSG00000227811.2 | 181541 | 299 | 152 | 5.51E-01 | -0.06 | 0.10 | 0.96 |
| Brain_Caudate_basal_ganglia | ENSG00000197852.8 | 168128 | 160 | 88 | 5.52E-01 | -0.06 | 0.10 | 0.96 |
| Brain_Caudate_basal_ganglia | ENSG00000134255.9 | 781156 | 160 | 88 | 5.52E-01 | 0.04 | 0.07 | 0.96 |
| Brain_Amygdala | ENSG00000273010.1 | 958080 | 100 | 47 | 5.53E-01 | 0.11 | 0.18 | 0.96 |

| | | | | | | | | |
|---|---|---|---|---|---|---|---|---|
| Esophagus_Muscularis | ENSG00000273483.1 | -597059 | 370 | 188 | 5.53E-01 | -0.03 | 0.05 | 0.96 |
| Brain_Spinal_cord_cervical_c-1 | ENSG00000116473.10 | 379164 | 91 | 49 | 5.53E-01 | -0.04 | 0.06 | 0.96 |
| Heart_Left_Ventricle | ENSG00000155363.14 | -751759 | 303 | 154 | 5.53E-01 | -0.02 | 0.03 | 0.96 |
| Brain_Hippocampus | ENSG00000134255.9 | 781156 | 123 | 63 | 5.55E-01 | 0.04 | 0.07 | 0.96 |
| Brain_Hippocampus | ENSG00000116459.6 | 472490 | 123 | 63 | 5.55E-01 | 0.03 | 0.05 | 0.96 |
| Brain_Hippocampus | ENSG00000273010.1 | 958080 | 123 | 63 | 5.55E-01 | -0.09 | 0.15 | 0.96 |
| Stomach | ENSG00000121933.13 | 357420 | 262 | 107 | 5.57E-01 | -0.04 | 0.06 | 0.96 |
| Brain_Hippocampus | ENSG00000260948.1 | 488314 | 123 | 63 | 5.59E-01 | 0.08 | 0.13 | 0.96 |
| Muscle_Skeletal | ENSG00000116473.10 | 379164 | 564 | 276 | 5.59E-01 | -0.01 | 0.02 | 0.96 |
| Colon_Sigmoid | ENSG00000260948.1 | 488314 | 233 | 107 | 5.62E-01 | 0.05 | 0.08 | 0.96 |
| Heart_Atrial_Appendage | ENSG00000116473.10 | 379164 | 297 | 147 | 5.63E-01 | -0.03 | 0.05 | 0.96 |
| Small_Intestine_Terminal_Ileum | ENSG00000238975.1 | -731206 | 137 | 54 | 5.63E-01 | 0.07 | 0.13 | 0.96 |
| Heart_Atrial_Appendage | ENSG00000134245.13 | -545159 | 297 | 147 | 5.65E-01 | 0.04 | 0.07 | 0.96 |
| Brain_Cerebellar_Hemisphere | ENSG00000116473.10 | 379164 | 136 | 74 | 5.66E-01 | -0.03 | 0.06 | 0.96 |
| Brain_Cerebellar_Hemisphere | ENSG00000184599.9 | -799037 | 136 | 74 | 5.66E-01 | -0.06 | 0.10 | 0.96 |
| Whole_Blood | ENSG00000155366.12 | -786052 | 407 | 195 | 5.66E-01 | 0.01 | 0.03 | 0.96 |
| Muscle_Skeletal | ENSG00000227811.2 | 181541 | 564 | 276 | 5.68E-01 | -0.03 | 0.06 | 0.96 |
| Thyroid | ENSG00000134216.14 | 630520 | 446 | 224 | 5.69E-01 | 0.03 | 0.06 | 0.96 |
| Prostate | ENSG00000155367.11 | -794095 | 152 | 75 | 5.70E-01 | -0.06 | 0.10 | 0.96 |
| Pancreas | ENSG00000121931.11 | 968421 | 248 | 108 | 5.70E-01 | -0.04 | 0.08 | 0.96 |
| Heart_Left_Ventricle | ENSG00000225075.1 | -775239 | 303 | 154 | 5.70E-01 | 0.05 | 0.10 | 0.96 |
| Adipose_Visceral_Omentum | ENSG00000261654.1 | 979029 | 355 | 172 | 5.72E-01 | 0.04 | 0.06 | 0.96 |
| Thyroid | ENSG00000171385.5 | -67773 | 446 | 224 | 5.72E-01 | -0.02 | 0.04 | 0.96 |
| Ovary | ENSG00000227811.2 | 181541 | 133 | 71 | 5.73E-01 | -0.06 | 0.11 | 0.96 |
| Breast_Mammary_Tissue | ENSG00000261654.1 | 979029 | 290 | 148 | 5.73E-01 | -0.04 | 0.07 | 0.96 |
| Thyroid | ENSG00000231246.1 | -439146 | 446 | 224 | 5.74E-01 | 0.03 | 0.06 | 0.96 |
| Whole_Blood | ENSG00000171385.5 | -67773 | 407 | 195 | 5.74E-01 | 0.04 | 0.06 | 0.96 |
| Artery_Aorta | ENSG00000064703.7 | 166137 | 299 | 152 | 5.76E-01 | 0.02 | 0.03 | 0.96 |
| Adrenal_Gland | ENSG00000231346.1 | 312659 | 190 | 96 | 5.77E-01 | 0.04 | 0.07 | 0.96 |
| Brain_Substantia_nigra | ENSG00000224167.1 | -955558 | 88 | 49 | 5.78E-01 | -0.08 | 0.14 | 0.96 |
| Whole_Blood | ENSG00000215867.4 | 271838 | 407 | 195 | 5.78E-01 | -0.04 | 0.07 | 0.96 |
| Pituitary | ENSG00000243960.1 | 482557 | 183 | 95 | 5.78E-01 | 0.08 | 0.14 | 0.96 |
| Small_Intestine_Terminal_Ileum | ENSG00000121931.11 | 968421 | 137 | 54 | 5.79E-01 | 0.06 | 0.11 | 0.96 |
| Heart_Left_Ventricle | ENSG00000143079.10 | -474799 | 303 | 154 | 5.83E-01 | 0.02 | 0.04 | 0.96 |
| Muscle_Skeletal | ENSG00000233337.1 | 483868 | 564 | 276 | 5.83E-01 | -0.03 | 0.06 | 0.96 |
| Brain_Hippocampus | ENSG00000116489.8 | -698415 | 123 | 63 | 5.84E-01 | 0.05 | 0.08 | 0.96 |
| Vagina | ENSG00000231246.1 | -439146 | 115 | 59 | 5.85E-01 | -0.05 | 0.10 | 0.96 |
| Colon_Transverse | ENSG00000143079.10 | -474799 | 274 | 119 | 5.85E-01 | 0.02 | 0.04 | 0.96 |
| Skin_Not_Sun_Exposed_Suprapubic | ENSG00000156171.10 | 781166 | 387 | 200 | 5.85E-01 | -0.02 | 0.04 | 0.96 |
| Heart_Atrial_Appendage | ENSG00000085465.11 | 493605 | 297 | 147 | 5.86E-01 | 0.02 | 0.04 | 0.96 |
| Stomach | ENSG00000232811.1 | 977915 | 262 | 107 | 5.86E-01 | -0.05 | 0.08 | 0.96 |
| Artery_Aorta | ENSG00000197852.8 | 168128 | 299 | 152 | 5.86E-01 | 0.02 | 0.04 | 0.96 |
| Adipose_Subcutaneous | ENSG00000155363.14 | -751759 | 442 | 214 | 5.87E-01 | -0.02 | 0.03 | 0.96 |
| Brain_Caudate_basal_ganglia | ENSG00000173947.9 | 575094 | 160 | 88 | 5.88E-01 | 0.03 | 0.06 | 0.96 |
| Skin_Not_Sun_Exposed_Suprapubic | ENSG00000121931.11 | 968421 | 387 | 200 | 5.88E-01 | 0.03 | 0.05 | 0.96 |
| Uterus | ENSG00000197852.8 | 168128 | 111 | 63 | 5.88E-01 | 0.05 | 0.09 | 0.96 |
| Nerve_Tibial | ENSG00000215866.3 | -929261 | 414 | 202 | 5.89E-01 | -0.03 | 0.06 | 0.96 |

| | | | | | | | | |
|---|---|---|---|---|---|---|---|---|
| Nerve_Tibial | ENSG00000273010.1 | 958080 | 414 | 202 | 5.89E-01 | -0.04 | 0.07 | 0.96 |
| Artery_Tibial | ENSG00000173947.9 | 575094 | 441 | 219 | 5.92E-01 | -0.03 | 0.05 | 0.96 |
| Skin_Sun_Exposed_Lower_leg | ENSG00000231346.1 | 312659 | 473 | 233 | 5.94E-01 | -0.03 | 0.05 | 0.96 |
| Brain_Cerebellar_Hemisphere | ENSG00000261654.1 | 979029 | 136 | 74 | 5.94E-01 | -0.06 | 0.11 | 0.96 |
| Adrenal_Gland | ENSG00000155363.14 | -751759 | 190 | 96 | 5.94E-01 | 0.03 | 0.06 | 0.96 |
| Whole_Blood | ENSG00000162777.12 | 716847 | 407 | 195 | 5.94E-01 | 0.01 | 0.03 | 0.96 |
| Whole_Blood | ENSG00000143079.10 | -474799 | 407 | 195 | 5.94E-01 | -0.02 | 0.04 | 0.96 |
| Brain_Putamen_basal_ganglia | ENSG00000273010.1 | 958080 | 124 | 71 | 5.95E-01 | 0.07 | 0.14 | 0.96 |
| Brain_Anterior_cingulate_cortex_BA24 | ENSG00000231346.1 | 312659 | 121 | 65 | 5.95E-01 | -0.07 | 0.13 | 0.96 |
| Colon_Transverse | ENSG00000121931.11 | 968421 | 274 | 119 | 5.96E-01 | -0.04 | 0.07 | 0.96 |
| Esophagus_Gastroesophageal_Junction | ENSG00000116455.9 | 472519 | 244 | 110 | 5.96E-01 | -0.03 | 0.05 | 0.96 |
| Esophagus_Muscularis | ENSG00000116489.8 | -698415 | 370 | 188 | 5.96E-01 | -0.02 | 0.04 | 0.96 |
| Stomach | ENSG00000155366.12 | -786052 | 262 | 107 | 5.98E-01 | -0.02 | 0.03 | 0.96 |
| Brain_Cerebellum | ENSG00000173947.9 | 575094 | 173 | 96 | 5.98E-01 | 0.05 | 0.09 | 0.96 |
| Muscle_Skeletal | ENSG00000232811.1 | 977915 | 564 | 276 | 5.99E-01 | -0.03 | 0.06 | 0.96 |
| Heart_Atrial_Appendage | ENSG00000116459.6 | 472490 | 297 | 147 | 5.99E-01 | 0.01 | 0.02 | 0.96 |
| Brain_Spinal_cord_cervical_c-1 | ENSG00000155363.14 | -751759 | 91 | 49 | 5.99E-01 | -0.05 | 0.09 | 0.96 |
| Esophagus_Muscularis | ENSG00000155366.12 | -786052 | 370 | 188 | 6.01E-01 | -0.02 | 0.04 | 0.96 |
| Brain_Substantia_nigra | ENSG00000260948.1 | 488314 | 88 | 49 | 6.02E-01 | 0.09 | 0.16 | 0.96 |
| Adipose_Visceral_Omentum | ENSG00000273483.1 | -597059 | 355 | 172 | 6.03E-01 | 0.04 | 0.07 | 0.96 |
| Esophagus_Muscularis | ENSG00000233337.1 | 483868 | 370 | 188 | 6.03E-01 | 0.04 | 0.08 | 0.96 |
| Skin_Not_Sun_Exposed_Suprapubic | ENSG00000233337.1 | 483868 | 387 | 200 | 6.03E-01 | 0.03 | 0.06 | 0.96 |
| Adipose_Subcutaneous | ENSG00000085465.11 | 493605 | 442 | 214 | 6.04E-01 | 0.02 | 0.03 | 0.96 |
| Esophagus_Mucosa | ENSG00000231346.1 | 312659 | 407 | 208 | 6.04E-01 | 0.03 | 0.05 | 0.96 |
| Brain_Nucleus_accumbens_basal_ganglia | ENSG00000116459.6 | 472490 | 147 | 77 | 6.04E-01 | -0.03 | 0.07 | 0.96 |
| Brain_Hippocampus | ENSG00000007341.14 | -699443 | 123 | 63 | 6.05E-01 | -0.06 | 0.12 | 0.96 |
| Brain_Hippocampus | ENSG00000121931.11 | 968421 | 123 | 63 | 6.06E-01 | 0.06 | 0.12 | 0.96 |
| Spleen | ENSG00000224167.1 | -955558 | 162 | 55 | 6.06E-01 | 0.07 | 0.13 | 0.96 |
| Breast_Mammary_Tissue | ENSG00000155363.14 | -751759 | 290 | 148 | 6.06E-01 | 0.02 | 0.03 | 0.96 |
| Small_Intestine_Terminal_Ileum | ENSG00000064703.7 | 166137 | 137 | 54 | 6.07E-01 | 0.04 | 0.08 | 0.96 |
| Brain_Putamen_basal_ganglia | ENSG00000116455.9 | 472519 | 124 | 71 | 6.08E-01 | -0.06 | 0.11 | 0.96 |
| Thyroid | ENSG00000121933.13 | 357420 | 446 | 224 | 6.08E-01 | -0.02 | 0.03 | 0.96 |
| Heart_Left_Ventricle | ENSG00000064703.7 | 166137 | 303 | 154 | 6.08E-01 | 0.02 | 0.05 | 0.96 |
| Brain_Caudate_basal_ganglia | ENSG00000215866.3 | -929261 | 160 | 88 | 6.10E-01 | 0.06 | 0.12 | 0.96 |
| Vagina | ENSG00000116473.10 | 379164 | 115 | 59 | 6.10E-01 | -0.04 | 0.08 | 0.96 |
| Heart_Left_Ventricle | ENSG00000173947.9 | 575094 | 303 | 154 | 6.11E-01 | -0.02 | 0.04 | 0.96 |
| Brain_Spinal_cord_cervical_c-1 | ENSG00000273010.1 | 958080 | 91 | 49 | 6.11E-01 | -0.10 | 0.20 | 0.96 |
| Nerve_Tibial | ENSG00000238975.1 | -731206 | 414 | 202 | 6.12E-01 | 0.04 | 0.07 | 0.96 |
| Lung | ENSG00000231346.1 | 312659 | 427 | 222 | 6.12E-01 | 0.03 | 0.05 | 0.96 |
| Stomach | ENSG00000116459.6 | 472490 | 262 | 107 | 6.13E-01 | -0.02 | 0.03 | 0.96 |
| Minor_Salivary_Gland | ENSG00000155367.11 | -794095 | 97 | 49 | 6.13E-01 | -0.06 | 0.11 | 0.96 |
| Brain_Cerebellum | ENSG00000134245.13 | -545159 | 173 | 96 | 6.13E-01 | 0.05 | 0.09 | 0.96 |
| Prostate | ENSG00000085465.11 | 493605 | 152 | 75 | 6.14E-01 | 0.04 | 0.08 | 0.96 |
| Ovary | ENSG00000173947.9 | 575094 | 133 | 71 | 6.15E-01 | 0.04 | 0.09 | 0.96 |
| Ovary | ENSG00000233337.1 | 483868 | 133 | 71 | 6.15E-01 | -0.05 | 0.11 | 0.96 |
| Lung | ENSG00000116459.6 | 472490 | 427 | 222 | 6.15E-01 | 0.01 | 0.02 | 0.96 |
| Adipose_Subcutaneous | ENSG00000121931.11 | 968421 | 442 | 214 | 6.15E-01 | -0.02 | 0.04 | 0.96 |

| | | | | | | | | |
|---|---|---|---|---|---|---|---|---|
| Nerve_Tibial | ENSG00000243960.1 | 482557 | 414 | 202 | 6.15E-01 | 0.04 | 0.08 | 0.96 |
| Brain_Cerebellar_Hemisphere | ENSG00000231437.3 | -68388 | 136 | 74 | 6.16E-01 | 0.04 | 0.09 | 0.96 |
| Colon_Transverse | ENSG00000243960.1 | 482557 | 274 | 119 | 6.17E-01 | -0.05 | 0.09 | 0.96 |
| Esophagus_Gastroesophageal_Junction | ENSG00000156171.10 | 781166 | 244 | 110 | 6.18E-01 | -0.03 | 0.07 | 0.96 |
| Nerve_Tibial | ENSG00000134255.9 | 781156 | 414 | 202 | 6.18E-01 | 0.02 | 0.04 | 0.96 |
| Brain_Putamen_basal_ganglia | ENSG00000134255.9 | 781156 | 124 | 71 | 6.19E-01 | 0.03 | 0.06 | 0.96 |
| Brain_Frontal_Cortex_BA9 | ENSG00000116489.8 | -698415 | 129 | 73 | 6.19E-01 | 0.05 | 0.09 | 0.96 |
| Colon_Transverse | ENSG00000155363.14 | -751759 | 274 | 119 | 6.19E-01 | -0.02 | 0.05 | 0.96 |
| Heart_Left_Ventricle | ENSG00000232811.1 | 977915 | 303 | 154 | 6.19E-01 | -0.05 | 0.10 | 0.96 |
| Small_Intestine_Terminal_Ileum | ENSG00000227811.2 | 181541 | 137 | 54 | 6.20E-01 | 0.06 | 0.13 | 0.96 |
| Brain_Hypothalamus | ENSG00000134245.13 | -545159 | 121 | 61 | 6.21E-01 | -0.05 | 0.11 | 0.96 |
| Artery_Tibial | ENSG00000243960.1 | 482557 | 441 | 219 | 6.21E-01 | 0.04 | 0.08 | 0.96 |
| Brain_Cerebellum | ENSG00000232811.1 | 977915 | 173 | 96 | 6.22E-01 | -0.05 | 0.10 | 0.96 |
| Ovary | ENSG00000162777.12 | 716847 | 133 | 71 | 6.23E-01 | -0.06 | 0.11 | 0.96 |
| Brain_Nucleus_accumbens_basal_ganglia | ENSG00000232811.1 | 977915 | 147 | 77 | 6.23E-01 | 0.06 | 0.13 | 0.96 |
| Cells_EBV-transformed_lymphocytes | ENSG00000231346.1 | 312659 | 130 | 68 | 6.23E-01 | 0.05 | 0.09 | 0.96 |
| Brain_Cerebellar_Hemisphere | ENSG00000085465.11 | 493605 | 136 | 74 | 6.24E-01 | 0.03 | 0.06 | 0.96 |
| Uterus | ENSG00000232811.1 | 977915 | 111 | 63 | 6.24E-01 | -0.07 | 0.15 | 0.96 |
| Artery_Aorta | ENSG00000156171.10 | 781166 | 299 | 152 | 6.24E-01 | -0.03 | 0.06 | 0.96 |
| Stomach | ENSG00000227179.2 | 529563 | 262 | 107 | 6.25E-01 | -0.04 | 0.08 | 0.96 |
| Minor_Salivary_Gland | ENSG00000116459.6 | 472490 | 97 | 49 | 6.26E-01 | 0.03 | 0.06 | 0.96 |
| Esophagus_Mucosa | ENSG00000143110.7 | 447590 | 407 | 208 | 6.27E-01 | -0.02 | 0.04 | 0.96 |
| Muscle_Skeletal | ENSG00000155366.12 | -786052 | 564 | 276 | 6.27E-01 | -0.02 | 0.03 | 0.96 |
| Ovary | ENSG00000197852.8 | 168128 | 133 | 71 | 6.27E-01 | -0.04 | 0.08 | 0.96 |
| Liver | ENSG00000273010.1 | 958080 | 175 | 88 | 6.27E-01 | 0.06 | 0.13 | 0.96 |
| Testis | ENSG00000273483.1 | -597059 | 259 | 128 | 6.28E-01 | -0.03 | 0.06 | 0.96 |
| Esophagus_Mucosa | ENSG00000134255.9 | 781156 | 407 | 208 | 6.28E-01 | -0.02 | 0.04 | 0.96 |
| Esophagus_Mucosa | ENSG00000155367.11 | -794095 | 407 | 208 | 6.28E-01 | -0.02 | 0.04 | 0.96 |
| Prostate | ENSG00000064703.7 | 166137 | 152 | 75 | 6.28E-01 | 0.03 | 0.07 | 0.96 |
| Spleen | ENSG00000162777.12 | 716847 | 162 | 55 | 6.29E-01 | 0.04 | 0.08 | 0.96 |
| Vagina | ENSG00000225075.1 | -775239 | 115 | 59 | 6.29E-01 | 0.08 | 0.16 | 0.96 |
| Brain_Hypothalamus | ENSG00000116473.10 | 379164 | 121 | 61 | 6.30E-01 | 0.02 | 0.05 | 0.96 |
| Brain_Anterior_cingulate_cortex_BA24 | ENSG00000116489.8 | -698415 | 121 | 65 | 6.30E-01 | 0.04 | 0.08 | 0.96 |
| Brain_Substantia_nigra | ENSG00000085465.11 | 493605 | 88 | 49 | 6.30E-01 | 0.04 | 0.08 | 0.96 |
| Heart_Atrial_Appendage | ENSG00000116455.9 | 472519 | 297 | 147 | 6.31E-01 | -0.02 | 0.03 | 0.96 |
| Brain_Cerebellar_Hemisphere | ENSG00000215866.3 | -929261 | 136 | 74 | 6.31E-01 | -0.06 | 0.13 | 0.96 |
| Adrenal_Gland | ENSG00000231246.1 | -439146 | 190 | 96 | 6.31E-01 | -0.05 | 0.10 | 0.96 |
| Brain_Nucleus_accumbens_basal_ganglia | ENSG00000064886.9 | 720611 | 147 | 77 | 6.33E-01 | -0.05 | 0.10 | 0.96 |
| Skin_Sun_Exposed_Lower_leg | ENSG00000134245.13 | -545159 | 473 | 233 | 6.33E-01 | -0.01 | 0.03 | 0.96 |
| Cells_Transformed_fibroblasts | ENSG00000134245.13 | -545159 | 343 | 175 | 6.34E-01 | -0.02 | 0.05 | 0.96 |
| Heart_Atrial_Appendage | ENSG00000273010.1 | 958080 | 297 | 147 | 6.34E-01 | -0.04 | 0.08 | 0.96 |
| Adrenal_Gland | ENSG00000116459.6 | 472490 | 190 | 96 | 6.35E-01 | -0.02 | 0.04 | 0.96 |
| Lung | ENSG00000064703.7 | 166137 | 427 | 222 | 6.35E-01 | 0.02 | 0.04 | 0.96 |
| Pituitary | ENSG00000215866.3 | -929261 | 183 | 95 | 6.38E-01 | -0.07 | 0.14 | 0.96 |
| Vagina | ENSG00000232811.1 | 977915 | 115 | 59 | 6.38E-01 | -0.07 | 0.14 | 0.96 |
| Colon_Sigmoid | ENSG00000085465.11 | 493605 | 233 | 107 | 6.38E-01 | -0.03 | 0.05 | 0.96 |

| | | | | | | | | |
|---|---|---|---|---|---|---|---|---|
| Brain_Cortex | ENSG00000184599.9 | -799037 | 158 | 79 | 6.38E-01 | -0.06 | 0.13 | 0.96 |
| Brain_Amygdala | ENSG00000064703.7 | 166137 | 100 | 47 | 6.40E-01 | 0.07 | 0.14 | 0.96 |
| Adrenal_Gland | ENSG00000116489.8 | -698415 | 190 | 96 | 6.41E-01 | 0.02 | 0.05 | 0.96 |
| Colon_Sigmoid | ENSG00000231346.1 | 312659 | 233 | 107 | 6.41E-01 | -0.03 | 0.06 | 0.96 |
| Adrenal_Gland | ENSG00000225075.1 | -775239 | 190 | 96 | 6.43E-01 | 0.05 | 0.10 | 0.96 |
| Artery_Tibial | ENSG00000231437.3 | -68388 | 441 | 219 | 6.43E-01 | -0.02 | 0.05 | 0.96 |
| Muscle_Skeletal | ENSG00000261654.1 | 979029 | 564 | 276 | 6.43E-01 | 0.02 | 0.05 | 0.96 |
| Brain_Caudate_basal_ganglia | ENSG00000116459.6 | 472490 | 160 | 88 | 6.44E-01 | -0.03 | 0.06 | 0.96 |
| Brain_Putamen_basal_ganglia | ENSG00000243960.1 | 482557 | 124 | 71 | 6.46E-01 | -0.07 | 0.15 | 0.96 |
| Muscle_Skeletal | ENSG00000155367.11 | -794095 | 564 | 276 | 6.48E-01 | -0.01 | 0.02 | 0.96 |
| Brain_Hippocampus | ENSG00000143079.10 | -474799 | 123 | 63 | 6.48E-01 | -0.03 | 0.07 | 0.96 |
| Thyroid | ENSG00000231346.1 | 312659 | 446 | 224 | 6.48E-01 | -0.02 | 0.05 | 0.96 |
| Whole_Blood | ENSG00000273483.1 | -597059 | 407 | 195 | 6.49E-01 | 0.03 | 0.06 | 0.96 |
| Brain_Amygdala | ENSG00000116473.10 | 379164 | 100 | 47 | 6.49E-01 | 0.03 | 0.06 | 0.96 |
| Artery_Tibial | ENSG00000143110.7 | 447590 | 441 | 219 | 6.50E-01 | -0.02 | 0.03 | 0.96 |
| Cells_EBV-transformed_lymphocytes | ENSG00000173947.9 | 575094 | 130 | 68 | 6.50E-01 | -0.06 | 0.12 | 0.96 |
| Pituitary | ENSG00000116459.6 | 472490 | 183 | 95 | 6.51E-01 | -0.02 | 0.05 | 0.96 |
| Brain_Cerebellum | ENSG00000155366.12 | -786052 | 173 | 96 | 6.52E-01 | 0.02 | 0.04 | 0.96 |
| Adipose_Subcutaneous | ENSG00000260948.1 | 488314 | 442 | 214 | 6.52E-01 | -0.02 | 0.05 | 0.96 |
| Adrenal_Gland | ENSG00000121931.11 | 968421 | 190 | 96 | 6.52E-01 | 0.03 | 0.08 | 0.96 |
| Colon_Transverse | ENSG00000231437.3 | -68388 | 274 | 119 | 6.53E-01 | -0.03 | 0.06 | 0.96 |
| Brain_Nucleus_accumbens_basal_ganglia | ENSG00000116455.9 | 472519 | 147 | 77 | 6.54E-01 | -0.05 | 0.11 | 0.96 |
| Brain_Putamen_basal_ganglia | ENSG00000171385.5 | -67773 | 124 | 71 | 6.54E-01 | 0.04 | 0.08 | 0.96 |
| Cells_Transformed_fibroblasts | ENSG00000197852.8 | 168128 | 343 | 175 | 6.54E-01 | 0.02 | 0.05 | 0.96 |
| Testis | ENSG00000162777.12 | 716847 | 259 | 128 | 6.54E-01 | 0.02 | 0.05 | 0.96 |
| Cells_Transformed_fibroblasts | ENSG00000162777.12 | 716847 | 343 | 175 | 6.58E-01 | -0.02 | 0.05 | 0.96 |
| Adrenal_Gland | ENSG00000260948.1 | 488314 | 190 | 96 | 6.59E-01 | 0.04 | 0.10 | 0.96 |
| Nerve_Tibial | ENSG00000064703.7 | 166137 | 414 | 202 | 6.60E-01 | -0.02 | 0.04 | 0.96 |
| Vagina | ENSG00000121931.11 | 968421 | 115 | 59 | 6.61E-01 | -0.05 | 0.11 | 0.96 |
| Minor_Salivary_Gland | ENSG00000064703.7 | 166137 | 97 | 49 | 6.61E-01 | 0.05 | 0.12 | 0.96 |
| Cells_EBV-transformed_lymphocytes | ENSG00000116473.10 | 379164 | 130 | 68 | 6.62E-01 | 0.04 | 0.08 | 0.96 |
| Small_Intestine_Terminal_Ileum | ENSG00000143110.7 | 447590 | 137 | 54 | 6.62E-01 | 0.05 | 0.10 | 0.96 |
| Stomach | ENSG00000225075.1 | -775239 | 262 | 107 | 6.64E-01 | -0.04 | 0.09 | 0.96 |
| Brain_Frontal_Cortex_BA9 | ENSG00000116455.9 | 472519 | 129 | 73 | 6.64E-01 | -0.04 | 0.09 | 0.96 |
| Brain_Substantia_nigra | ENSG00000134245.13 | -545159 | 88 | 49 | 6.65E-01 | -0.05 | 0.12 | 0.96 |
| Lung | ENSG00000143079.10 | -474799 | 427 | 222 | 6.66E-01 | 0.01 | 0.03 | 0.96 |
| Brain_Substantia_nigra | ENSG00000273483.1 | -597059 | 88 | 49 | 6.66E-01 | 0.06 | 0.15 | 0.96 |
| Pituitary | ENSG00000064703.7 | 166137 | 183 | 95 | 6.67E-01 | -0.03 | 0.07 | 0.96 |
| Artery_Aorta | ENSG00000134255.9 | 781156 | 299 | 152 | 6.67E-01 | 0.02 | 0.04 | 0.96 |
| Brain_Hypothalamus | ENSG00000116459.6 | 472490 | 121 | 61 | 6.68E-01 | -0.02 | 0.06 | 0.96 |
| Pituitary | ENSG00000116489.8 | -698415 | 183 | 95 | 6.69E-01 | 0.02 | 0.05 | 0.96 |
| Uterus | ENSG00000231346.1 | 312659 | 111 | 63 | 6.70E-01 | -0.04 | 0.10 | 0.96 |
| Artery_Coronary | ENSG00000116489.8 | -698415 | 173 | 81 | 6.71E-01 | -0.02 | 0.05 | 0.96 |
| Heart_Atrial_Appendage | ENSG00000143079.10 | -474799 | 297 | 147 | 6.71E-01 | 0.02 | 0.04 | 0.96 |
| Brain_Amygdala | ENSG00000273483.1 | -597059 | 100 | 47 | 6.72E-01 | 0.05 | 0.13 | 0.96 |
| Skin_Sun_Exposed_Lower_leg | ENSG00000224167.1 | -955558 | 473 | 233 | 6.72E-01 | 0.03 | 0.06 | 0.96 |
| Brain_Cerebellar_Hemisphere | ENSG00000156171.10 | 781166 | 136 | 74 | 6.73E-01 | -0.04 | 0.10 | 0.96 |

| | | | | | | | | |
|---|---|---|---|---|---|---|---|---|
| Stomach | ENSG00000231246.1 | -439146 | 262 | 107 | 6.74E-01 | -0.03 | 0.08 | 0.96 |
| Brain_Hippocampus | ENSG00000231346.1 | 312659 | 123 | 63 | 6.74E-01 | 0.05 | 0.13 | 0.96 |
| Brain_Amygdala | ENSG00000116489.8 | -698415 | 100 | 47 | 6.74E-01 | 0.05 | 0.11 | 0.96 |
| Esophagus_Muscularis | ENSG00000134255.9 | 781156 | 370 | 188 | 6.76E-01 | 0.02 | 0.05 | 0.96 |
| Artery_Tibial | ENSG00000116455.9 | 472519 | 441 | 219 | 6.76E-01 | 0.01 | 0.03 | 0.96 |
| Brain_Hypothalamus | ENSG00000155363.14 | -751759 | 121 | 61 | 6.77E-01 | -0.03 | 0.07 | 0.96 |
| Muscle_Skeletal | ENSG00000143079.10 | -474799 | 564 | 276 | 6.77E-01 | -0.01 | 0.03 | 0.96 |
| Brain_Putamen_basal_ganglia | ENSG00000116473.10 | 379164 | 124 | 71 | 6.77E-01 | 0.02 | 0.06 | 0.96 |
| Cells_EBV-transformed_lymphocytes | ENSG00000232811.1 | 977915 | 130 | 68 | 6.77E-01 | 0.04 | 0.11 | 0.96 |
| Brain_Cortex | ENSG00000116455.9 | 472519 | 158 | 79 | 6.77E-01 | -0.05 | 0.12 | 0.96 |
| Brain_Anterior_cingulate_cortex_BA24 | ENSG00000260948.1 | 488314 | 121 | 65 | 6.78E-01 | 0.05 | 0.12 | 0.96 |
| Testis | ENSG00000184599.9 | -799037 | 259 | 128 | 6.78E-01 | 0.02 | 0.05 | 0.96 |
| Brain_Hypothalamus | ENSG00000215866.3 | -929261 | 121 | 61 | 6.78E-01 | -0.05 | 0.12 | 0.96 |
| Prostate | ENSG00000197852.8 | 168128 | 152 | 75 | 6.78E-01 | 0.04 | 0.10 | 0.96 |
| Vagina | ENSG00000155363.14 | -751759 | 115 | 59 | 6.78E-01 | 0.03 | 0.07 | 0.96 |
| Thyroid | ENSG00000197852.8 | 168128 | 446 | 224 | 6.79E-01 | 0.02 | 0.04 | 0.96 |
| Prostate | ENSG00000116459.6 | 472490 | 152 | 75 | 6.79E-01 | -0.02 | 0.05 | 0.96 |
| Brain_Hypothalamus | ENSG00000143079.10 | -474799 | 121 | 61 | 6.79E-01 | -0.03 | 0.07 | 0.96 |
| Testis | ENSG00000233337.1 | 483868 | 259 | 128 | 6.80E-01 | 0.04 | 0.09 | 0.96 |
| Brain_Frontal_Cortex_BA9 | ENSG00000231346.1 | 312659 | 129 | 73 | 6.81E-01 | -0.06 | 0.14 | 0.96 |
| Skin_Sun_Exposed_Lower_leg | ENSG00000116459.6 | 472490 | 473 | 233 | 6.81E-01 | -0.01 | 0.02 | 0.96 |
| Brain_Hypothalamus | ENSG00000116489.8 | -698415 | 121 | 61 | 6.82E-01 | -0.04 | 0.09 | 0.96 |
| Brain_Amygdala | ENSG00000116455.9 | 472519 | 100 | 47 | 6.83E-01 | -0.05 | 0.13 | 0.96 |
| Prostate | ENSG00000007341.14 | -699443 | 152 | 75 | 6.83E-01 | 0.05 | 0.12 | 0.96 |
| Lung | ENSG00000116455.9 | 472519 | 427 | 222 | 6.84E-01 | -0.01 | 0.03 | 0.96 |
| Artery_Coronary | ENSG00000155363.14 | -751759 | 173 | 81 | 6.84E-01 | -0.02 | 0.04 | 0.96 |
| Pancreas | ENSG00000162777.12 | 716847 | 248 | 108 | 6.84E-01 | 0.02 | 0.04 | 0.96 |
| Whole_Blood | ENSG00000260948.1 | 488314 | 407 | 195 | 6.84E-01 | 0.02 | 0.06 | 0.96 |
| Stomach | ENSG00000134245.13 | -545159 | 262 | 107 | 6.84E-01 | -0.02 | 0.06 | 0.96 |
| Heart_Left_Ventricle | ENSG00000273483.1 | -597059 | 303 | 154 | 6.85E-01 | 0.03 | 0.06 | 0.96 |
| Brain_Cortex | ENSG00000227811.2 | 181541 | 158 | 79 | 6.85E-01 | -0.05 | 0.13 | 0.96 |
| Stomach | ENSG00000134216.14 | 630520 | 262 | 107 | 6.86E-01 | -0.02 | 0.06 | 0.96 |
| Cells_Transformed_fibroblasts | ENSG00000173947.9 | 575094 | 343 | 175 | 6.86E-01 | 0.02 | 0.04 | 0.96 |
| Esophagus_Muscularis | ENSG00000231346.1 | 312659 | 370 | 188 | 6.87E-01 | 0.01 | 0.03 | 0.96 |
| Brain_Cerebellar_Hemisphere | ENSG00000064886.9 | 720611 | 136 | 74 | 6.87E-01 | -0.05 | 0.12 | 0.96 |
| Brain_Cerebellar_Hemisphere | ENSG00000155366.12 | -786052 | 136 | 74 | 6.87E-01 | 0.02 | 0.06 | 0.96 |
| Brain_Amygdala | ENSG00000232811.1 | 977915 | 100 | 47 | 6.87E-01 | 0.09 | 0.21 | 0.96 |
| Skin_Not_Sun_Exposed_Suprapubic | ENSG00000134255.9 | 781156 | 387 | 200 | 6.88E-01 | -0.02 | 0.04 | 0.96 |
| Skin_Not_Sun_Exposed_Suprapubic | ENSG00000231346.1 | 312659 | 387 | 200 | 6.91E-01 | 0.02 | 0.05 | 0.96 |
| Thyroid | ENSG00000231437.3 | -68388 | 446 | 224 | 6.91E-01 | 0.02 | 0.05 | 0.96 |
| Skin_Sun_Exposed_Lower_leg | ENSG00000215866.3 | -929261 | 473 | 233 | 6.92E-01 | 0.02 | 0.06 | 0.96 |
| Stomach | ENSG00000173947.9 | 575094 | 262 | 107 | 6.92E-01 | -0.02 | 0.06 | 0.96 |
| Adipose_Subcutaneous | ENSG00000184599.9 | -799037 | 442 | 214 | 6.93E-01 | 0.03 | 0.06 | 0.96 |
| Nerve_Tibial | ENSG00000231246.1 | -439146 | 414 | 202 | 6.93E-01 | -0.02 | 0.06 | 0.96 |
| Heart_Atrial_Appendage | ENSG00000134255.9 | 781156 | 297 | 147 | 6.93E-01 | 0.02 | 0.05 | 0.96 |
| Vagina | ENSG00000260948.1 | 488314 | 115 | 59 | 6.93E-01 | -0.05 | 0.13 | 0.96 |
| Brain_Spinal_cord_cervical_c-1 | ENSG00000064886.9 | 720611 | 91 | 49 | 6.95E-01 | -0.05 | 0.13 | 0.96 |

| | | | | | | | | |
|---|---|---|---|---|---|---|---|---|
| Cells_EBV-transformed_lymphocytes | ENSG00000155366.12 | -786052 | 130 | 68 | 6.95E-01 | -0.03 | 0.08 | 0.96 |
| Cells_EBV-transformed_lymphocytes | ENSG00000085465.11 | 493605 | 130 | 68 | 6.96E-01 | -0.04 | 0.10 | 0.96 |
| Cells_Transformed_fibroblasts | ENSG00000116489.8 | -698415 | 343 | 175 | 6.96E-01 | 0.01 | 0.02 | 0.96 |
| Skin_Not_Sun_Exposed_Suprapubic | ENSG00000227811.2 | 181541 | 387 | 200 | 6.96E-01 | 0.03 | 0.08 | 0.96 |
| Skin_Not_Sun_Exposed_Suprapubic | ENSG00000064703.7 | 166137 | 387 | 200 | 6.97E-01 | 0.02 | 0.05 | 0.96 |
| Prostate | ENSG00000116489.8 | -698415 | 152 | 75 | 6.98E-01 | -0.03 | 0.09 | 0.96 |
| Brain_Cerebellum | ENSG00000162777.12 | 716847 | 173 | 96 | 6.98E-01 | -0.03 | 0.08 | 0.96 |
| Brain_Amygdala | ENSG00000134245.13 | -545159 | 100 | 47 | 6.98E-01 | 0.05 | 0.13 | 0.96 |
| Brain_Nucleus_accumbens_basal_ganglia | ENSG00000173947.9 | 575094 | 147 | 77 | 6.99E-01 | 0.03 | 0.09 | 0.96 |
| Cells_Transformed_fibroblasts | ENSG00000134255.9 | 781156 | 343 | 175 | 6.99E-01 | 0.02 | 0.05 | 0.96 |
| Liver | ENSG00000121933.13 | 357420 | 175 | 88 | 7.00E-01 | 0.03 | 0.07 | 0.96 |
| Uterus | ENSG00000156171.10 | 781166 | 111 | 63 | 7.01E-01 | -0.04 | 0.11 | 0.96 |
| Artery_Aorta | ENSG00000116455.9 | 472519 | 299 | 152 | 7.01E-01 | -0.01 | 0.03 | 0.96 |
| Stomach | ENSG00000171385.5 | -67773 | 262 | 107 | 7.02E-01 | 0.02 | 0.05 | 0.96 |
| Heart_Left_Ventricle | ENSG00000231246.1 | -439146 | 303 | 154 | 7.03E-01 | -0.03 | 0.08 | 0.96 |
| Whole_Blood | ENSG00000064886.9 | 720611 | 407 | 195 | 7.03E-01 | 0.02 | 0.06 | 0.96 |
| Ovary | ENSG00000156171.10 | 781166 | 133 | 71 | 7.04E-01 | -0.03 | 0.09 | 0.96 |
| Pituitary | ENSG00000134255.9 | 781156 | 183 | 95 | 7.05E-01 | -0.03 | 0.07 | 0.96 |
| Artery_Coronary | ENSG00000155366.12 | -786052 | 173 | 81 | 7.05E-01 | -0.02 | 0.05 | 0.96 |
| Stomach | ENSG00000227811.2 | 181541 | 262 | 107 | 7.05E-01 | 0.03 | 0.07 | 0.96 |
| Brain_Hippocampus | ENSG00000155366.12 | -786052 | 123 | 63 | 7.06E-01 | -0.02 | 0.05 | 0.96 |
| Muscle_Skeletal | ENSG00000243960.1 | 482557 | 564 | 276 | 7.06E-01 | -0.02 | 0.07 | 0.96 |
| Cells_EBV-transformed_lymphocytes | ENSG00000134216.14 | 630520 | 130 | 68 | 7.07E-01 | -0.05 | 0.13 | 0.96 |
| Cells_EBV-transformed_lymphocytes | ENSG00000134245.13 | -545159 | 130 | 68 | 7.08E-01 | -0.04 | 0.10 | 0.96 |
| Heart_Atrial_Appendage | ENSG00000231437.3 | -68388 | 297 | 147 | 7.09E-01 | 0.03 | 0.07 | 0.96 |
| Thyroid | ENSG00000162777.12 | 716847 | 446 | 224 | 7.10E-01 | 0.01 | 0.04 | 0.96 |
| Esophagus_Gastroesophageal_Junction | ENSG00000121933.13 | 357420 | 244 | 110 | 7.10E-01 | 0.02 | 0.06 | 0.96 |
| Muscle_Skeletal | ENSG00000231437.3 | -68388 | 564 | 276 | 7.11E-01 | -0.02 | 0.05 | 0.96 |
| Brain_Hypothalamus | ENSG00000224167.1 | -955558 | 121 | 61 | 7.12E-01 | 0.05 | 0.14 | 0.96 |
| Ovary | ENSG00000116489.8 | -698415 | 133 | 71 | 7.13E-01 | 0.03 | 0.09 | 0.96 |
| Whole_Blood | ENSG00000227811.2 | 181541 | 407 | 195 | 7.14E-01 | -0.02 | 0.06 | 0.96 |
| Adipose_Visceral_Omentum | ENSG00000156171.10 | 781166 | 355 | 172 | 7.14E-01 | -0.02 | 0.05 | 0.96 |
| Brain_Cortex | ENSG00000273010.1 | 958080 | 158 | 79 | 7.15E-01 | -0.05 | 0.13 | 0.96 |
| Brain_Caudate_basal_ganglia | ENSG00000227811.2 | 181541 | 160 | 88 | 7.15E-01 | 0.05 | 0.13 | 0.96 |
| Whole_Blood | ENSG00000273010.1 | 958080 | 407 | 195 | 7.15E-01 | -0.03 | 0.08 | 0.96 |
| Thyroid | ENSG00000121931.11 | 968421 | 446 | 224 | 7.15E-01 | -0.01 | 0.04 | 0.96 |
| Artery_Coronary | ENSG00000231437.3 | -68388 | 173 | 81 | 7.15E-01 | 0.03 | 0.07 | 0.96 |
| Artery_Aorta | ENSG00000155366.12 | -786052 | 299 | 152 | 7.16E-01 | 0.01 | 0.03 | 0.96 |
| Skin_Not_Sun_Exposed_Suprapubic | ENSG00000261654.1 | 979029 | 387 | 200 | 7.17E-01 | 0.02 | 0.07 | 0.96 |
| Ovary | ENSG00000143110.7 | 447590 | 133 | 71 | 7.17E-01 | -0.03 | 0.09 | 0.96 |
| Adipose_Visceral_Omentum | ENSG00000197852.8 | 168128 | 355 | 172 | 7.17E-01 | 0.01 | 0.03 | 0.96 |
| Brain_Caudate_basal_ganglia | ENSG00000116489.8 | -698415 | 160 | 88 | 7.17E-01 | -0.02 | 0.05 | 0.96 |
| Uterus | ENSG00000134245.13 | -545159 | 111 | 63 | 7.19E-01 | 0.04 | 0.12 | 0.96 |
| Adipose_Subcutaneous | ENSG00000143079.10 | -474799 | 442 | 214 | 7.19E-01 | 0.01 | 0.03 | 0.96 |
| Brain_Cerebellar_Hemisphere | ENSG00000233337.1 | 483868 | 136 | 74 | 7.19E-01 | 0.04 | 0.10 | 0.96 |
| Vagina | ENSG00000156171.10 | 781166 | 115 | 59 | 7.20E-01 | -0.04 | 0.10 | 0.96 |
| Whole_Blood | ENSG00000233337.1 | 483868 | 407 | 195 | 7.21E-01 | 0.03 | 0.08 | 0.96 |

| | | | | | | | | |
|---|---|---|---|---|---|---|---|---|
| Brain_Amygdala | ENSG00000155366.12 | -786052 | 100 | 47 | 7.21E-01 | 0.02 | 0.07 | 0.96 |
| Adipose_Subcutaneous | ENSG00000232811.1 | 977915 | 442 | 214 | 7.21E-01 | 0.03 | 0.08 | 0.96 |
| Spleen | ENSG00000134245.13 | -545159 | 162 | 55 | 7.22E-01 | -0.04 | 0.10 | 0.96 |
| Brain_Amygdala | ENSG00000085465.11 | 493605 | 100 | 47 | 7.22E-01 | 0.04 | 0.11 | 0.96 |
| Uterus | ENSG00000225075.1 | -775239 | 111 | 63 | 7.23E-01 | -0.05 | 0.13 | 0.96 |
| Lung | ENSG00000162777.12 | 716847 | 427 | 222 | 7.23E-01 | -0.01 | 0.03 | 0.96 |
| Breast_Mammary_Tissue | ENSG00000116473.10 | 379164 | 290 | 148 | 7.23E-01 | -0.01 | 0.03 | 0.96 |
| Testis | ENSG00000064886.9 | 720611 | 259 | 128 | 7.24E-01 | 0.03 | 0.08 | 0.96 |
| Pituitary | ENSG00000134245.13 | -545159 | 183 | 95 | 7.24E-01 | 0.04 | 0.11 | 0.96 |
| Brain_Putamen_basal_ganglia | ENSG00000173947.9 | 575094 | 124 | 71 | 7.25E-01 | 0.04 | 0.11 | 0.96 |
| Minor_Salivary_Gland | ENSG00000116455.9 | 472519 | 97 | 49 | 7.25E-01 | -0.03 | 0.08 | 0.96 |
| Thyroid | ENSG00000085465.11 | 493605 | 446 | 224 | 7.26E-01 | 0.01 | 0.03 | 0.96 |
| Colon_Transverse | ENSG00000155366.12 | -786052 | 274 | 119 | 7.26E-01 | -0.01 | 0.04 | 0.96 |
| Cells_EBV-transformed_lymphocytes | ENSG00000227811.2 | 181541 | 130 | 68 | 7.26E-01 | 0.05 | 0.14 | 0.96 |
| Skin_Not_Sun_Exposed_Suprapubic | ENSG00000231437.3 | -68388 | 387 | 200 | 7.27E-01 | -0.02 | 0.05 | 0.96 |
| Esophagus_Gastroesophageal_Junction | ENSG00000260948.1 | 488314 | 244 | 110 | 7.27E-01 | -0.03 | 0.09 | 0.96 |
| Brain_Nucleus_accumbens_basal_ganglia | ENSG00000231437.3 | -68388 | 147 | 77 | 7.28E-01 | 0.03 | 0.08 | 0.96 |
| Adipose_Visceral_Omentum | ENSG00000173947.9 | 575094 | 355 | 172 | 7.28E-01 | -0.02 | 0.05 | 0.96 |
| Brain_Amygdala | ENSG00000260948.1 | 488314 | 100 | 47 | 7.29E-01 | -0.06 | 0.17 | 0.96 |
| Prostate | ENSG00000134245.13 | -545159 | 152 | 75 | 7.30E-01 | 0.03 | 0.09 | 0.96 |
| Artery_Coronary | ENSG00000007341.14 | -699443 | 173 | 81 | 7.31E-01 | -0.03 | 0.09 | 0.96 |
| Vagina | ENSG00000261654.1 | 979029 | 115 | 59 | 7.32E-01 | -0.04 | 0.13 | 0.96 |
| Vagina | ENSG00000273483.1 | -597059 | 115 | 59 | 7.32E-01 | -0.04 | 0.11 | 0.96 |
| Nerve_Tibial | ENSG00000197852.8 | 168128 | 414 | 202 | 7.33E-01 | 0.01 | 0.03 | 0.96 |
| Prostate | ENSG00000134255.9 | 781156 | 152 | 75 | 7.33E-01 | -0.03 | 0.07 | 0.96 |
| Esophagus_Mucosa | ENSG00000155363.14 | -751759 | 407 | 208 | 7.34E-01 | -0.01 | 0.03 | 0.96 |
| Cells_EBV-transformed_lymphocytes | ENSG00000260948.1 | 488314 | 130 | 68 | 7.34E-01 | 0.04 | 0.10 | 0.96 |
| Testis | ENSG00000225075.1 | -775239 | 259 | 128 | 7.36E-01 | 0.03 | 0.08 | 0.96 |
| Brain_Anterior_cingulate_cortex_BA24 | ENSG00000116473.10 | 379164 | 121 | 65 | 7.36E-01 | -0.02 | 0.05 | 0.96 |
| Esophagus_Muscularis | ENSG00000173947.9 | 575094 | 370 | 188 | 7.37E-01 | 0.01 | 0.04 | 0.96 |
| Vagina | ENSG00000171385.5 | -67773 | 115 | 59 | 7.37E-01 | 0.04 | 0.13 | 0.96 |
| Lung | ENSG00000231437.3 | -68388 | 427 | 222 | 7.38E-01 | 0.02 | 0.05 | 0.96 |
| Adipose_Subcutaneous | ENSG00000064703.7 | 166137 | 442 | 214 | 7.38E-01 | -0.01 | 0.04 | 0.96 |
| Colon_Sigmoid | ENSG00000116459.6 | 472490 | 233 | 107 | 7.39E-01 | -0.01 | 0.03 | 0.96 |
| Uterus | ENSG00000231246.1 | -439146 | 111 | 63 | 7.39E-01 | 0.04 | 0.11 | 0.96 |
| Minor_Salivary_Gland | ENSG00000116473.10 | 379164 | 97 | 49 | 7.39E-01 | -0.04 | 0.10 | 0.96 |
| Skin_Sun_Exposed_Lower_leg | ENSG00000162777.12 | 716847 | 473 | 233 | 7.39E-01 | -0.01 | 0.02 | 0.96 |
| Cells_EBV-transformed_lymphocytes | ENSG00000155367.11 | -794095 | 130 | 68 | 7.40E-01 | -0.03 | 0.09 | 0.96 |
| Colon_Transverse | ENSG00000064886.9 | 720611 | 274 | 119 | 7.41E-01 | -0.02 | 0.06 | 0.96 |
| Artery_Tibial | ENSG00000231246.1 | -439146 | 441 | 219 | 7.41E-01 | -0.02 | 0.06 | 0.96 |
| Brain_Caudate_basal_ganglia | ENSG00000116455.9 | 472519 | 160 | 88 | 7.41E-01 | 0.03 | 0.10 | 0.96 |
| Cells_Transformed_fibroblasts | ENSG00000171385.5 | -67773 | 343 | 175 | 7.42E-01 | 0.01 | 0.04 | 0.96 |
| Brain_Putamen_basal_ganglia | ENSG00000184599.9 | -799037 | 124 | 71 | 7.42E-01 | -0.05 | 0.15 | 0.96 |
| Vagina | ENSG00000121933.13 | 357420 | 115 | 59 | 7.42E-01 | -0.03 | 0.09 | 0.96 |
| Brain_Substantia_nigra | ENSG00000064886.9 | 720611 | 88 | 49 | 7.43E-01 | 0.04 | 0.12 | 0.96 |
| Esophagus_Muscularis | ENSG00000261654.1 | 979029 | 370 | 188 | 7.44E-01 | -0.02 | 0.07 | 0.96 |
| Skin_Not_Sun_Exposed_Suprapubic | ENSG00000121933.13 | 357420 | 387 | 200 | 7.44E-01 | 0.01 | 0.04 | 0.96 |

| | | | | | | | | |
|---|---|---|---|---|---|---|---|---|
| Ovary | ENSG00000121933.13 | 357420 | 133 | 71 | 7.44E-01 | -0.02 | 0.07 | 0.96 |
| Liver | ENSG00000116459.6 | 472490 | 175 | 88 | 7.45E-01 | 0.01 | 0.04 | 0.96 |
| Lung | ENSG00000203878.7 | 641323 | 427 | 222 | 7.46E-01 | 0.02 | 0.06 | 0.96 |
| Artery_Coronary | ENSG00000155367.11 | -794095 | 173 | 81 | 7.48E-01 | 0.02 | 0.06 | 0.96 |
| Artery_Aorta | ENSG00000233337.1 | 483868 | 299 | 152 | 7.48E-01 | 0.03 | 0.09 | 0.96 |
| Colon_Transverse | ENSG00000155367.11 | -794095 | 274 | 119 | 7.48E-01 | 0.02 | 0.07 | 0.96 |
| Heart_Left_Ventricle | ENSG00000231346.1 | 312659 | 303 | 154 | 7.51E-01 | 0.02 | 0.06 | 0.96 |
| Muscle_Skeletal | ENSG00000143110.7 | 447590 | 564 | 276 | 7.51E-01 | -0.01 | 0.04 | 0.96 |
| Whole_Blood | ENSG00000231437.3 | -68388 | 407 | 195 | 7.52E-01 | 0.02 | 0.07 | 0.96 |
| Brain_Cortex | ENSG00000085465.11 | 493605 | 158 | 79 | 7.53E-01 | 0.02 | 0.07 | 0.96 |
| Artery_Coronary | ENSG00000085465.11 | 493605 | 173 | 81 | 7.53E-01 | -0.02 | 0.06 | 0.96 |
| Nerve_Tibial | ENSG00000155363.14 | -751759 | 414 | 202 | 7.53E-01 | -0.01 | 0.03 | 0.96 |
| Whole_Blood | ENSG00000085465.11 | 493605 | 407 | 195 | 7.54E-01 | -0.01 | 0.03 | 0.96 |
| Stomach | ENSG00000064886.9 | 720611 | 262 | 107 | 7.54E-01 | -0.02 | 0.06 | 0.96 |
| Skin_Sun_Exposed_Lower_leg | ENSG00000260948.1 | 488314 | 473 | 233 | 7.55E-01 | 0.01 | 0.03 | 0.96 |
| Esophagus_Muscularis | ENSG00000121931.11 | 968421 | 370 | 188 | 7.58E-01 | -0.01 | 0.05 | 0.96 |
| Brain_Spinal_cord_cervical_c-1 | ENSG00000116489.8 | -698415 | 91 | 49 | 7.58E-01 | 0.03 | 0.10 | 0.96 |
| Brain_Hypothalamus | ENSG00000232811.1 | 977915 | 121 | 61 | 7.58E-01 | -0.04 | 0.14 | 0.96 |
| Whole_Blood | ENSG00000155367.11 | -794095 | 407 | 195 | 7.59E-01 | -0.02 | 0.06 | 0.96 |
| Cells_EBV-transformed_lymphocytes | ENSG00000162777.12 | 716847 | 130 | 68 | 7.60E-01 | 0.03 | 0.10 | 0.96 |
| Heart_Left_Ventricle | ENSG00000231437.3 | -68388 | 303 | 154 | 7.60E-01 | 0.02 | 0.06 | 0.96 |
| Small_Intestine_Terminal_Ileum | ENSG00000231246.1 | -439146 | 137 | 54 | 7.60E-01 | -0.04 | 0.13 | 0.96 |
| Small_Intestine_Terminal_Ileum | ENSG00000260948.1 | 488314 | 137 | 54 | 7.61E-01 | -0.02 | 0.08 | 0.96 |
| Muscle_Skeletal | ENSG00000064703.7 | 166137 | 564 | 276 | 7.61E-01 | -0.01 | 0.03 | 0.96 |
| Heart_Left_Ventricle | ENSG00000121931.11 | 968421 | 303 | 154 | 7.62E-01 | -0.01 | 0.05 | 0.96 |
| Brain_Spinal_cord_cervical_c-1 | ENSG00000116455.9 | 472519 | 91 | 49 | 7.63E-01 | -0.04 | 0.15 | 0.96 |
| Ovary | ENSG00000231346.1 | 312659 | 133 | 71 | 7.64E-01 | -0.03 | 0.09 | 0.96 |
| Brain_Substantia_nigra | ENSG00000143079.10 | -474799 | 88 | 49 | 7.64E-01 | 0.03 | 0.10 | 0.96 |
| Esophagus_Mucosa | ENSG00000162777.12 | 716847 | 407 | 208 | 7.64E-01 | 0.01 | 0.03 | 0.96 |
| Brain_Hypothalamus | ENSG00000273010.1 | 958080 | 121 | 61 | 7.65E-01 | 0.04 | 0.14 | 0.96 |
| Heart_Atrial_Appendage | ENSG00000232811.1 | 977915 | 297 | 147 | 7.65E-01 | -0.03 | 0.09 | 0.96 |
| Nerve_Tibial | ENSG00000085465.11 | 493605 | 414 | 202 | 7.66E-01 | -0.01 | 0.03 | 0.96 |
| Artery_Coronary | ENSG00000232811.1 | 977915 | 173 | 81 | 7.66E-01 | 0.03 | 0.12 | 0.96 |
| Uterus | ENSG00000064886.9 | 720611 | 111 | 63 | 7.68E-01 | -0.04 | 0.14 | 0.96 |
| Vagina | ENSG00000184599.9 | -799037 | 115 | 59 | 7.68E-01 | -0.04 | 0.13 | 0.96 |
| Stomach | ENSG00000134255.9 | 781156 | 262 | 107 | 7.69E-01 | -0.01 | 0.04 | 0.96 |
| Artery_Coronary | ENSG00000243960.1 | 482557 | 173 | 81 | 7.70E-01 | -0.04 | 0.13 | 0.96 |
| Lung | ENSG00000233337.1 | 483868 | 427 | 222 | 7.70E-01 | -0.02 | 0.06 | 0.96 |
| Brain_Putamen_basal_ganglia | ENSG00000121931.11 | 968421 | 124 | 71 | 7.70E-01 | 0.03 | 0.11 | 0.96 |
| Cells_Transformed_fibroblasts | ENSG00000155367.11 | -794095 | 343 | 175 | 7.71E-01 | 0.01 | 0.05 | 0.96 |
| Brain_Nucleus_accumbens_basal_ganglia | ENSG00000273010.1 | 958080 | 147 | 77 | 7.71E-01 | 0.04 | 0.13 | 0.96 |
| Heart_Atrial_Appendage | ENSG00000231246.1 | -439146 | 297 | 147 | 7.72E-01 | -0.02 | 0.07 | 0.96 |
| Brain_Anterior_cingulate_cortex_BA24 | ENSG00000116455.9 | 472519 | 121 | 65 | 7.74E-01 | -0.03 | 0.10 | 0.96 |
| Adipose_Visceral_Omentum | ENSG00000085465.11 | 493605 | 355 | 172 | 7.74E-01 | 0.01 | 0.04 | 0.96 |
| Brain_Cerebellar_Hemisphere | ENSG00000173947.9 | 575094 | 136 | 74 | 7.74E-01 | -0.03 | 0.10 | 0.96 |
| Artery_Aorta | ENSG00000162777.12 | 716847 | 299 | 152 | 7.74E-01 | 0.01 | 0.04 | 0.96 |
| Esophagus_Muscularis | ENSG00000121933.13 | 357420 | 370 | 188 | 7.75E-01 | -0.01 | 0.04 | 0.96 |

| | | | | | | | | |
|---|---|---|---|---|---|---|---|---|
| Brain_Anterior_cingulate_cortex_BA24 | ENSG00000243960.1 | 482557 | 121 | 65 | 7.75E-01 | -0.04 | 0.14 | 0.96 |
| Brain_Putamen_basal_ganglia | ENSG00000156171.10 | 781166 | 124 | 71 | 7.75E-01 | -0.04 | 0.12 | 0.96 |
| Adrenal_Gland | ENSG00000134255.9 | 781156 | 190 | 96 | 7.76E-01 | 0.02 | 0.06 | 0.96 |
| Liver | ENSG00000155366.12 | -786052 | 175 | 88 | 7.76E-01 | 0.02 | 0.06 | 0.96 |
| Brain_Caudate_basal_ganglia | ENSG00000233337.1 | 483868 | 160 | 88 | 7.77E-01 | 0.03 | 0.11 | 0.96 |
| Pancreas | ENSG00000064886.9 | 720611 | 248 | 108 | 7.79E-01 | -0.03 | 0.09 | 0.96 |
| Skin_Sun_Exposed_Lower_leg | ENSG00000134255.9 | 781156 | 473 | 233 | 7.79E-01 | 0.01 | 0.04 | 0.96 |
| Brain_Substantia_nigra | ENSG00000007341.14 | -699443 | 88 | 49 | 7.79E-01 | -0.04 | 0.14 | 0.96 |
| Spleen | ENSG00000064886.9 | 720611 | 162 | 55 | 7.79E-01 | -0.04 | 0.14 | 0.96 |
| Spleen | ENSG00000134255.9 | 781156 | 162 | 55 | 7.79E-01 | -0.02 | 0.09 | 0.96 |
| Esophagus_Muscularis | ENSG00000116459.6 | 472490 | 370 | 188 | 7.80E-01 | 0.01 | 0.02 | 0.96 |
| Artery_Aorta | ENSG00000260948.1 | 488314 | 299 | 152 | 7.80E-01 | -0.02 | 0.07 | 0.96 |
| Brain_Cortex | ENSG00000215866.3 | -929261 | 158 | 79 | 7.81E-01 | 0.04 | 0.14 | 0.96 |
| Brain_Cortex | ENSG00000121931.11 | 968421 | 158 | 79 | 7.81E-01 | 0.03 | 0.10 | 0.96 |
| Colon_Transverse | ENSG00000231246.1 | -439146 | 274 | 119 | 7.82E-01 | -0.02 | 0.08 | 0.96 |
| Colon_Sigmoid | ENSG00000227811.2 | 181541 | 233 | 107 | 7.82E-01 | 0.03 | 0.12 | 0.96 |
| Brain_Cortex | ENSG00000224167.1 | -955558 | 158 | 79 | 7.82E-01 | -0.04 | 0.14 | 0.96 |
| Nerve_Tibial | ENSG00000116459.6 | 472490 | 414 | 202 | 7.83E-01 | -0.01 | 0.02 | 0.96 |
| Brain_Frontal_Cortex_BA9 | ENSG00000261654.1 | 979029 | 129 | 73 | 7.83E-01 | 0.03 | 0.11 | 0.96 |
| Thyroid | ENSG00000243960.1 | 482557 | 446 | 224 | 7.83E-01 | 0.02 | 0.07 | 0.96 |
| Stomach | ENSG00000116455.9 | 472519 | 262 | 107 | 7.83E-01 | 0.01 | 0.05 | 0.96 |
| Brain_Substantia_nigra | ENSG00000064703.7 | 166137 | 88 | 49 | 7.84E-01 | 0.03 | 0.12 | 0.96 |
| Brain_Hypothalamus | ENSG00000197852.8 | 168128 | 121 | 61 | 7.84E-01 | 0.03 | 0.11 | 0.96 |
| Adrenal_Gland | ENSG00000273010.1 | 958080 | 190 | 96 | 7.85E-01 | -0.03 | 0.11 | 0.96 |
| Esophagus_Muscularis | ENSG00000231246.1 | -439146 | 370 | 188 | 7.87E-01 | -0.02 | 0.07 | 0.96 |
| Brain_Cerebellum | ENSG00000085465.11 | 493605 | 173 | 96 | 7.87E-01 | -0.02 | 0.06 | 0.96 |
| Nerve_Tibial | ENSG00000184599.9 | -799037 | 414 | 202 | 7.87E-01 | 0.02 | 0.06 | 0.96 |
| Artery_Coronary | ENSG00000231246.1 | -439146 | 173 | 81 | 7.87E-01 | 0.03 | 0.10 | 0.96 |
| Esophagus_Mucosa | ENSG00000231246.1 | -439146 | 407 | 208 | 7.87E-01 | -0.02 | 0.07 | 0.96 |
| Spleen | ENSG00000085465.11 | 493605 | 162 | 55 | 7.88E-01 | 0.03 | 0.09 | 0.97 |
| Brain_Caudate_basal_ganglia | ENSG00000243960.1 | 482557 | 160 | 88 | 7.89E-01 | -0.04 | 0.13 | 0.97 |
| Artery_Tibial | ENSG00000134245.13 | -545159 | 441 | 219 | 7.91E-01 | 0.01 | 0.04 | 0.97 |
| Small_Intestine_Terminal_Ileum | ENSG00000233337.1 | 483868 | 137 | 54 | 7.91E-01 | -0.03 | 0.11 | 0.97 |
| Colon_Transverse | ENSG00000225075.1 | -775239 | 274 | 119 | 7.93E-01 | 0.02 | 0.09 | 0.97 |
| Lung | ENSG00000007341.14 | -699443 | 427 | 222 | 7.94E-01 | -0.01 | 0.06 | 0.97 |
| Brain_Anterior_cingulate_cortex_BA24 | ENSG00000224167.1 | -955558 | 121 | 65 | 7.94E-01 | 0.04 | 0.14 | 0.97 |
| Pancreas | ENSG00000155366.12 | -786052 | 248 | 108 | 7.94E-01 | -0.01 | 0.04 | 0.97 |
| Skin_Sun_Exposed_Lower_leg | ENSG00000155367.11 | -794095 | 473 | 233 | 7.95E-01 | 0.01 | 0.03 | 0.97 |
| Prostate | ENSG00000231246.1 | -439146 | 152 | 75 | 7.95E-01 | -0.03 | 0.10 | 0.97 |
| Artery_Aorta | ENSG00000143110.7 | 447590 | 299 | 152 | 7.99E-01 | -0.01 | 0.04 | 0.97 |
| Adrenal_Gland | ENSG00000162777.12 | 716847 | 190 | 96 | 8.00E-01 | -0.02 | 0.07 | 0.97 |
| Artery_Coronary | ENSG00000116459.6 | 472490 | 173 | 81 | 8.01E-01 | -0.01 | 0.03 | 0.97 |
| Brain_Caudate_basal_ganglia | ENSG00000260948.1 | 488314 | 160 | 88 | 8.03E-01 | -0.02 | 0.10 | 0.97 |
| Spleen | ENSG00000116455.9 | 472519 | 162 | 55 | 8.04E-01 | 0.02 | 0.09 | 0.97 |
| Prostate | ENSG00000231437.3 | -68388 | 152 | 75 | 8.05E-01 | 0.02 | 0.08 | 0.97 |
| Muscle_Skeletal | ENSG00000121931.11 | 968421 | 564 | 276 | 8.06E-01 | 0.01 | 0.04 | 0.97 |
| Brain_Nucleus_accumbens_basal_ganglia | ENSG00000171385.5 | -67773 | 147 | 77 | 8.06E-01 | -0.01 | 0.04 | 0.97 |

| | | | | | | | | |
|---|---|---|---|---|---|---|---|---|
| Brain_Frontal_Cortex_BA9 | ENSG00000171385.5 | -67773 | 129 | 73 | 8.06E-01 | 0.01 | 0.06 | 0.97 |
| Esophagus_Muscularis | ENSG00000227811.2 | 181541 | 370 | 188 | 8.08E-01 | 0.02 | 0.08 | 0.97 |
| Ovary | ENSG00000231246.1 | -439146 | 133 | 71 | 8.08E-01 | 0.03 | 0.10 | 0.97 |
| Prostate | ENSG00000227811.2 | 181541 | 152 | 75 | 8.09E-01 | 0.03 | 0.12 | 0.97 |
| Esophagus_Gastroesophageal_Junction | ENSG00000231437.3 | -68388 | 244 | 110 | 8.09E-01 | -0.02 | 0.07 | 0.97 |
| Skin_Sun_Exposed_Lower_leg | ENSG00000155363.14 | -751759 | 473 | 233 | 8.09E-01 | 0.01 | 0.03 | 0.97 |
| Testis | ENSG00000261654.1 | 979029 | 259 | 128 | 8.10E-01 | -0.02 | 0.08 | 0.97 |
| Lung | ENSG00000273483.1 | -597059 | 427 | 222 | 8.10E-01 | 0.01 | 0.06 | 0.97 |
| Brain_Putamen_basal_ganglia | ENSG00000116489.8 | -698415 | 124 | 71 | 8.11E-01 | 0.02 | 0.08 | 0.97 |
| Brain_Putamen_basal_ganglia | ENSG00000064703.7 | 166137 | 124 | 71 | 8.12E-01 | 0.02 | 0.10 | 0.97 |
| Brain_Caudate_basal_ganglia | ENSG00000224167.1 | -955558 | 160 | 88 | 8.13E-01 | -0.03 | 0.13 | 0.97 |
| Heart_Left_Ventricle | ENSG00000243960.1 | 482557 | 303 | 154 | 8.14E-01 | -0.02 | 0.10 | 0.97 |
| Brain_Cerebellar_Hemisphere | ENSG00000171385.5 | -67773 | 136 | 74 | 8.16E-01 | -0.02 | 0.07 | 0.97 |
| Skin_Sun_Exposed_Lower_leg | ENSG00000121933.13 | 357420 | 473 | 233 | 8.17E-01 | 0.01 | 0.04 | 0.97 |
| Pituitary | ENSG00000134216.14 | 630520 | 183 | 95 | 8.18E-01 | -0.02 | 0.09 | 0.97 |
| Adrenal_Gland | ENSG00000121933.13 | 357420 | 190 | 96 | 8.18E-01 | 0.01 | 0.06 | 0.97 |
| Vagina | ENSG00000155367.11 | -794095 | 115 | 59 | 8.19E-01 | 0.02 | 0.07 | 0.97 |
| Brain_Anterior_cingulate_cortex_BA24 | ENSG00000121931.11 | 968421 | 121 | 65 | 8.19E-01 | 0.02 | 0.10 | 0.97 |
| Brain_Anterior_cingulate_cortex_BA24 | ENSG00000215866.3 | -929261 | 121 | 65 | 8.20E-01 | -0.03 | 0.13 | 0.97 |
| Breast_Mammary_Tissue | ENSG00000121931.11 | 968421 | 290 | 148 | 8.20E-01 | 0.01 | 0.06 | 0.97 |
| Breast_Mammary_Tissue | ENSG00000155367.11 | -794095 | 290 | 148 | 8.21E-01 | 0.01 | 0.04 | 0.97 |
| Whole_Blood | ENSG00000121931.11 | 968421 | 407 | 195 | 8.21E-01 | 0.01 | 0.02 | 0.97 |
| Heart_Left_Ventricle | ENSG00000233337.1 | 483868 | 303 | 154 | 8.22E-01 | -0.02 | 0.10 | 0.97 |
| Muscle_Skeletal | ENSG00000155363.14 | -751759 | 564 | 276 | 8.22E-01 | -0.01 | 0.03 | 0.97 |
| Adipose_Subcutaneous | ENSG00000197852.8 | 168128 | 442 | 214 | 8.22E-01 | 0.01 | 0.04 | 0.97 |
| Skin_Sun_Exposed_Lower_leg | ENSG00000233337.1 | 483868 | 473 | 233 | 8.23E-01 | 0.01 | 0.05 | 0.97 |
| Stomach | ENSG00000273483.1 | -597059 | 262 | 107 | 8.23E-01 | 0.02 | 0.09 | 0.97 |
| Cells_EBV-transformed_lymphocytes | ENSG00000233337.1 | 483868 | 130 | 68 | 8.23E-01 | 0.03 | 0.12 | 0.97 |
| Vagina | ENSG00000143110.7 | 447590 | 115 | 59 | 8.24E-01 | 0.02 | 0.08 | 0.97 |
| Ovary | ENSG00000273483.1 | -597059 | 133 | 71 | 8.24E-01 | 0.02 | 0.08 | 0.97 |
| Artery_Coronary | ENSG00000173947.9 | 575094 | 173 | 81 | 8.24E-01 | 0.02 | 0.09 | 0.97 |
| Brain_Anterior_cingulate_cortex_BA24 | ENSG00000156171.10 | 781166 | 121 | 65 | 8.24E-01 | -0.02 | 0.09 | 0.97 |
| Artery_Coronary | ENSG00000171385.5 | -67773 | 173 | 81 | 8.24E-01 | 0.02 | 0.09 | 0.97 |
| Uterus | ENSG00000121933.13 | 357420 | 111 | 63 | 8.27E-01 | -0.02 | 0.10 | 0.98 |
| Brain_Anterior_cingulate_cortex_BA24 | ENSG00000143110.7 | 447590 | 121 | 65 | 8.28E-01 | -0.01 | 0.06 | 0.98 |
| Brain_Cerebellar_Hemisphere | ENSG00000231346.1 | 312659 | 136 | 74 | 8.29E-01 | -0.03 | 0.12 | 0.98 |
| Cells_Transformed_fibroblasts | ENSG00000064703.7 | 166137 | 343 | 175 | 8.30E-01 | -0.01 | 0.03 | 0.98 |
| Brain_Cerebellum | ENSG00000121933.13 | 357420 | 173 | 96 | 8.30E-01 | -0.02 | 0.09 | 0.98 |
| Prostate | ENSG00000243960.1 | 482557 | 152 | 75 | 8.31E-01 | 0.02 | 0.11 | 0.98 |
| Prostate | ENSG00000233337.1 | 483868 | 152 | 75 | 8.32E-01 | 0.02 | 0.09 | 0.98 |
| Small_Intestine_Terminal_Ileum | ENSG00000155366.12 | -786052 | 137 | 54 | 8.33E-01 | -0.01 | 0.03 | 0.98 |
| Lung | ENSG00000273010.1 | 958080 | 427 | 222 | 8.33E-01 | -0.02 | 0.07 | 0.98 |
| Breast_Mammary_Tissue | ENSG00000231246.1 | -439146 | 290 | 148 | 8.33E-01 | 0.01 | 0.06 | 0.98 |
| Brain_Frontal_Cortex_BA9 | ENSG00000155366.12 | -786052 | 129 | 73 | 8.33E-01 | 0.01 | 0.03 | 0.98 |
| Thyroid | ENSG00000260948.1 | 488314 | 446 | 224 | 8.34E-01 | 0.01 | 0.05 | 0.98 |
| Pituitary | ENSG00000085465.11 | 493605 | 183 | 95 | 8.34E-01 | 0.02 | 0.10 | 0.98 |
| Pancreas | ENSG00000007341.14 | -699443 | 248 | 108 | 8.38E-01 | 0.02 | 0.08 | 0.98 |

| | | | | | | | | |
|---|---|---|---|---|---|---|---|---|
| Skin_Not_Sun_Exposed_Suprapubic | ENSG00000184599.9 | -799037 | 387 | 200 | 8.39E-01 | 0.01 | 0.06 | 0.98 |
| Adipose_Subcutaneous | ENSG00000261654.1 | 979029 | 442 | 214 | 8.42E-01 | 0.01 | 0.07 | 0.98 |
| Brain_Nucleus_accumbens_basal_ganglia | ENSG00000233337.1 | 483868 | 147 | 77 | 8.44E-01 | -0.03 | 0.13 | 0.98 |
| Colon_Transverse | ENSG00000162777.12 | 716847 | 274 | 119 | 8.46E-01 | -0.01 | 0.03 | 0.98 |
| Brain_Putamen_basal_ganglia | ENSG00000155366.12 | -786052 | 124 | 71 | 8.46E-01 | -0.01 | 0.05 | 0.98 |
| Cells_EBV-transformed_lymphocytes | ENSG00000215867.4 | 271838 | 130 | 68 | 8.46E-01 | 0.02 | 0.12 | 0.98 |
| Whole_Blood | ENSG00000134255.9 | 781156 | 407 | 195 | 8.47E-01 | 0.00 | 0.02 | 0.98 |
| Prostate | ENSG00000215866.3 | -929261 | 152 | 75 | 8.47E-01 | -0.02 | 0.11 | 0.98 |
| Minor_Salivary_Gland | ENSG00000225075.1 | -775239 | 97 | 49 | 8.47E-01 | -0.03 | 0.15 | 0.98 |
| Adipose_Visceral_Omentum | ENSG00000232811.1 | 977915 | 355 | 172 | 8.47E-01 | -0.01 | 0.07 | 0.98 |
| Artery_Tibial | ENSG00000085465.11 | 493605 | 441 | 219 | 8.48E-01 | 0.01 | 0.03 | 0.98 |
| Adipose_Visceral_Omentum | ENSG00000227179.2 | 529563 | 355 | 172 | 8.48E-01 | -0.01 | 0.06 | 0.98 |
| Artery_Aorta | ENSG00000173947.9 | 575094 | 299 | 152 | 8.48E-01 | 0.01 | 0.08 | 0.98 |
| Brain_Hypothalamus | ENSG00000273483.1 | -597059 | 121 | 61 | 8.48E-01 | -0.02 | 0.09 | 0.98 |
| Adipose_Visceral_Omentum | ENSG00000233337.1 | 483868 | 355 | 172 | 8.48E-01 | -0.01 | 0.07 | 0.98 |
| Ovary | ENSG00000121931.11 | 968421 | 133 | 71 | 8.49E-01 | -0.02 | 0.08 | 0.98 |
| Breast_Mammary_Tissue | ENSG00000184599.9 | -799037 | 290 | 148 | 8.50E-01 | 0.01 | 0.06 | 0.98 |
| Liver | ENSG00000134255.9 | 781156 | 175 | 88 | 8.51E-01 | -0.02 | 0.08 | 0.98 |
| Brain_Anterior_cingulate_cortex_BA24 | ENSG00000231437.3 | -68388 | 121 | 65 | 8.51E-01 | 0.02 | 0.11 | 0.98 |
| Pancreas | ENSG00000116459.6 | 472490 | 248 | 108 | 8.53E-01 | 0.01 | 0.05 | 0.98 |
| Cells_Transformed_fibroblasts | ENSG00000227811.2 | 181541 | 343 | 175 | 8.54E-01 | 0.01 | 0.07 | 0.98 |
| Brain_Putamen_basal_ganglia | ENSG00000116459.6 | 472490 | 124 | 71 | 8.55E-01 | -0.01 | 0.07 | 0.98 |
| Nerve_Tibial | ENSG00000116455.9 | 472519 | 414 | 202 | 8.55E-01 | -0.01 | 0.03 | 0.98 |
| Uterus | ENSG00000238975.1 | -731206 | 111 | 63 | 8.55E-01 | -0.03 | 0.14 | 0.98 |
| Brain_Amygdala | ENSG00000007341.14 | -699443 | 100 | 47 | 8.57E-01 | 0.03 | 0.17 | 0.99 |
| Liver | ENSG00000233337.1 | 483868 | 175 | 88 | 8.57E-01 | -0.02 | 0.10 | 0.99 |
| Esophagus_Muscularis | ENSG00000231437.3 | -68388 | 370 | 188 | 8.58E-01 | -0.01 | 0.05 | 0.99 |
| Skin_Not_Sun_Exposed_Suprapubic | ENSG00000134245.13 | -545159 | 387 | 200 | 8.59E-01 | -0.01 | 0.03 | 0.99 |
| Adrenal_Gland | ENSG00000171385.5 | -67773 | 190 | 96 | 8.60E-01 | 0.01 | 0.08 | 0.99 |
| Spleen | ENSG00000261654.1 | 979029 | 162 | 55 | 8.61E-01 | -0.02 | 0.11 | 0.99 |
| Brain_Cortex | ENSG00000134255.9 | 781156 | 158 | 79 | 8.62E-01 | 0.02 | 0.09 | 0.99 |
| Brain_Frontal_Cortex_BA9 | ENSG00000156171.10 | 781166 | 129 | 73 | 8.65E-01 | 0.01 | 0.09 | 0.99 |
| Pituitary | ENSG00000233337.1 | 483868 | 183 | 95 | 8.65E-01 | -0.02 | 0.11 | 0.99 |
| Skin_Not_Sun_Exposed_Suprapubic | ENSG00000173947.9 | 575094 | 387 | 200 | 8.66E-01 | 0.01 | 0.06 | 0.99 |
| Liver | ENSG00000085465.11 | 493605 | 175 | 88 | 8.66E-01 | -0.01 | 0.07 | 0.99 |
| Brain_Cortex | ENSG00000064703.7 | 166137 | 158 | 79 | 8.66E-01 | -0.02 | 0.10 | 0.99 |
| Brain_Caudate_basal_ganglia | ENSG00000116473.10 | 379164 | 160 | 88 | 8.67E-01 | -0.01 | 0.04 | 0.99 |
| Colon_Transverse | ENSG00000231346.1 | 312659 | 274 | 119 | 8.68E-01 | -0.01 | 0.05 | 0.99 |
| Pituitary | ENSG00000155363.14 | -751759 | 183 | 95 | 8.69E-01 | -0.01 | 0.08 | 0.99 |
| Artery_Coronary | ENSG00000273483.1 | -597059 | 173 | 81 | 8.69E-01 | -0.02 | 0.10 | 0.99 |
| Brain_Hippocampus | ENSG00000215866.3 | -929261 | 123 | 63 | 8.70E-01 | -0.02 | 0.12 | 0.99 |
| Brain_Cerebellum | ENSG00000116489.8 | -698415 | 173 | 96 | 8.71E-01 | 0.01 | 0.06 | 0.99 |
| Brain_Spinal_cord_cervical_c-1 | ENSG00000162777.12 | 716847 | 91 | 49 | 8.71E-01 | 0.02 | 0.12 | 0.99 |
| Lung | ENSG00000116489.8 | -698415 | 427 | 222 | 8.73E-01 | 0.00 | 0.02 | 0.99 |
| Adipose_Visceral_Omentum | ENSG00000155366.12 | -786052 | 355 | 172 | 8.74E-01 | 0.01 | 0.03 | 0.99 |
| Testis | ENSG00000173947.9 | 575094 | 259 | 128 | 8.74E-01 | 0.00 | 0.03 | 0.99 |
| Ovary | ENSG00000116459.6 | 472490 | 133 | 71 | 8.75E-01 | -0.01 | 0.06 | 0.99 |

| | | | | | | | | |
|---|---|---|---|---|---|---|---|---|
| Pituitary | ENSG00000261654.1 | 979029 | 183 | 95 | 8.78E-01 | -0.02 | 0.13 | 0.99 |
| Esophagus_Muscularis | ENSG00000085465.11 | 493605 | 370 | 188 | 8.78E-01 | -0.01 | 0.04 | 0.99 |
| Prostate | ENSG00000231346.1 | 312659 | 152 | 75 | 8.79E-01 | 0.01 | 0.08 | 0.99 |
| Brain_Frontal_Cortex_BA9 | ENSG00000121931.11 | 968421 | 129 | 73 | 8.80E-01 | -0.01 | 0.07 | 0.99 |
| Adrenal_Gland | ENSG00000155367.11 | -794095 | 190 | 96 | 8.80E-01 | 0.01 | 0.07 | 0.99 |
| Breast_Mammary_Tissue | ENSG00000134245.13 | -545159 | 290 | 148 | 8.80E-01 | 0.01 | 0.06 | 0.99 |
| Whole_Blood | ENSG00000116455.9 | 472519 | 407 | 195 | 8.80E-01 | -0.01 | 0.04 | 0.99 |
| Thyroid | ENSG00000184599.9 | -799037 | 446 | 224 | 8.82E-01 | -0.01 | 0.05 | 0.99 |
| Artery_Coronary | ENSG00000227811.2 | 181541 | 173 | 81 | 8.82E-01 | -0.02 | 0.13 | 0.99 |
| Brain_Frontal_Cortex_BA9 | ENSG00000162777.12 | 716847 | 129 | 73 | 8.82E-01 | -0.02 | 0.11 | 0.99 |
| Stomach | ENSG00000156171.10 | 781166 | 262 | 107 | 8.82E-01 | 0.01 | 0.06 | 0.99 |
| Brain_Hypothalamus | ENSG00000260948.1 | 488314 | 121 | 61 | 8.83E-01 | 0.02 | 0.12 | 0.99 |
| Small_Intestine_Terminal_Ileum | ENSG00000007341.14 | -699443 | 137 | 54 | 8.83E-01 | -0.02 | 0.11 | 0.99 |
| Heart_Left_Ventricle | ENSG00000197852.8 | 168128 | 303 | 154 | 8.83E-01 | 0.01 | 0.04 | 0.99 |
| Lung | ENSG00000171385.5 | -67773 | 427 | 222 | 8.84E-01 | 0.01 | 0.04 | 0.99 |
| Skin_Not_Sun_Exposed_Suprapubic | ENSG00000007341.14 | -699443 | 387 | 200 | 8.84E-01 | 0.01 | 0.04 | 0.99 |
| Brain_Hippocampus | ENSG00000197852.8 | 168128 | 123 | 63 | 8.84E-01 | 0.01 | 0.06 | 0.99 |
| Brain_Nucleus_accumbens_basal_ganglia | ENSG00000184599.9 | -799037 | 147 | 77 | 8.84E-01 | 0.02 | 0.12 | 0.99 |
| Pituitary | ENSG00000260948.1 | 488314 | 183 | 95 | 8.86E-01 | -0.02 | 0.12 | 0.99 |
| Adrenal_Gland | ENSG00000064886.9 | 720611 | 190 | 96 | 8.87E-01 | -0.01 | 0.08 | 0.99 |
| Spleen | ENSG00000156171.10 | 781166 | 162 | 55 | 8.87E-01 | 0.02 | 0.11 | 0.99 |
| Brain_Cerebellum | ENSG00000260948.1 | 488314 | 173 | 96 | 8.88E-01 | -0.01 | 0.06 | 0.99 |
| Artery_Aorta | ENSG00000231246.1 | -439146 | 299 | 152 | 8.88E-01 | -0.01 | 0.08 | 0.99 |
| Skin_Not_Sun_Exposed_Suprapubic | ENSG00000171385.5 | -67773 | 387 | 200 | 8.88E-01 | -0.01 | 0.05 | 0.99 |
| Heart_Left_Ventricle | ENSG00000007341.14 | -699443 | 303 | 154 | 8.88E-01 | 0.01 | 0.07 | 0.99 |
| Artery_Tibial | ENSG00000121933.13 | 357420 | 441 | 219 | 8.88E-01 | -0.01 | 0.04 | 0.99 |
| Breast_Mammary_Tissue | ENSG00000260948.1 | 488314 | 290 | 148 | 8.90E-01 | -0.01 | 0.07 | 0.99 |
| Brain_Cerebellum | ENSG00000233337.1 | 483868 | 173 | 96 | 8.90E-01 | 0.01 | 0.10 | 0.99 |
| Minor_Salivary_Gland | ENSG00000231246.1 | -439146 | 97 | 49 | 8.90E-01 | -0.02 | 0.12 | 0.99 |
| Brain_Nucleus_accumbens_basal_ganglia | ENSG00000143079.10 | -474799 | 147 | 77 | 8.91E-01 | 0.01 | 0.08 | 0.99 |
| Heart_Left_Ventricle | ENSG00000116459.6 | 472490 | 303 | 154 | 8.91E-01 | 0.00 | 0.02 | 0.99 |
| Heart_Atrial_Appendage | ENSG00000143110.7 | 447590 | 297 | 147 | 8.92E-01 | 0.01 | 0.04 | 0.99 |
| Adipose_Subcutaneous | ENSG00000227811.2 | 181541 | 442 | 214 | 8.93E-01 | 0.01 | 0.07 | 0.99 |
| Adipose_Subcutaneous | ENSG00000231246.1 | -439146 | 442 | 214 | 8.94E-01 | 0.01 | 0.07 | 0.99 |
| Brain_Cortex | ENSG00000173947.9 | 575094 | 158 | 79 | 8.96E-01 | -0.01 | 0.07 | 0.99 |
| Brain_Cerebellum | ENSG00000116459.6 | 472490 | 173 | 96 | 8.96E-01 | -0.01 | 0.05 | 0.99 |
| Heart_Left_Ventricle | ENSG00000116455.9 | 472519 | 303 | 154 | 8.96E-01 | 0.00 | 0.03 | 0.99 |
| Brain_Cerebellar_Hemisphere | ENSG00000143079.10 | -474799 | 136 | 74 | 8.97E-01 | -0.01 | 0.09 | 0.99 |
| Adrenal_Gland | ENSG00000261654.1 | 979029 | 190 | 96 | 8.98E-01 | -0.01 | 0.12 | 0.99 |
| Brain_Anterior_cingulate_cortex_BA24 | ENSG00000173947.9 | 575094 | 121 | 65 | 8.98E-01 | 0.02 | 0.12 | 0.99 |
| Artery_Tibial | ENSG00000116459.6 | 472490 | 441 | 219 | 8.98E-01 | 0.00 | 0.02 | 0.99 |
| Stomach | ENSG00000203878.7 | 641323 | 262 | 107 | 9.01E-01 | 0.01 | 0.07 | 0.99 |
| Colon_Sigmoid | ENSG00000143079.10 | -474799 | 233 | 107 | 9.01E-01 | 0.01 | 0.06 | 0.99 |
| Heart_Atrial_Appendage | ENSG00000273483.1 | -597059 | 297 | 147 | 9.02E-01 | -0.01 | 0.07 | 0.99 |
| Adipose_Subcutaneous | ENSG00000233337.1 | 483868 | 442 | 214 | 9.02E-01 | -0.01 | 0.07 | 0.99 |
| Heart_Atrial_Appendage | ENSG00000155367.11 | -794095 | 297 | 147 | 9.04E-01 | 0.01 | 0.05 | 0.99 |

| | | | | | | | | |
|---|---|---|---|---|---|---|---|---|
| Esophagus_Mucosa | ENSG00000121933.13 | 357420 | 407 | 208 | 9.05E-01 | 0.00 | 0.04 | 0.99 |
| Brain_Amygdala | ENSG00000121931.11 | 968421 | 100 | 47 | 9.05E-01 | 0.02 | 0.14 | 0.99 |
| Brain_Substantia_nigra | ENSG00000143110.7 | 447590 | 88 | 49 | 9.05E-01 | -0.01 | 0.08 | 0.99 |
| Uterus | ENSG00000231437.3 | -68388 | 111 | 63 | 9.05E-01 | -0.02 | 0.14 | 0.99 |
| Cells_Transformed_fibroblasts | ENSG00000116455.9 | 472519 | 343 | 175 | 9.07E-01 | 0.00 | 0.02 | 0.99 |
| Breast_Mammary_Tissue | ENSG00000085465.11 | 493605 | 290 | 148 | 9.07E-01 | -0.01 | 0.05 | 0.99 |
| Esophagus_Mucosa | ENSG00000238975.1 | -731206 | 407 | 208 | 9.07E-01 | -0.01 | 0.07 | 0.99 |
| Brain_Cortex | ENSG00000243960.1 | 482557 | 158 | 79 | 9.07E-01 | -0.02 | 0.14 | 0.99 |
| Thyroid | ENSG00000232811.1 | 977915 | 446 | 224 | 9.08E-01 | 0.01 | 0.06 | 0.99 |
| Spleen | ENSG00000143079.10 | -474799 | 162 | 55 | 9.08E-01 | -0.01 | 0.08 | 0.99 |
| Brain_Putamen_basal_ganglia | ENSG00000197852.8 | 168128 | 124 | 71 | 9.08E-01 | 0.01 | 0.13 | 0.99 |
| Adipose_Visceral_Omentum | ENSG00000273010.1 | 958080 | 355 | 172 | 9.09E-01 | -0.01 | 0.07 | 0.99 |
| Testis | ENSG00000121931.11 | 968421 | 259 | 128 | 9.10E-01 | 0.00 | 0.03 | 0.99 |
| Brain_Spinal_cord_cervical_c-1 | ENSG00000155367.11 | -794095 | 91 | 49 | 9.10E-01 | 0.01 | 0.10 | 0.99 |
| Brain_Caudate_basal_ganglia | ENSG00000155363.14 | -751759 | 160 | 88 | 9.10E-01 | 0.01 | 0.06 | 0.99 |
| Testis | ENSG00000224167.1 | -955558 | 259 | 128 | 9.10E-01 | 0.01 | 0.08 | 0.99 |
| Brain_Anterior_cingulate_cortex_BA24 | ENSG00000231246.1 | -439146 | 121 | 65 | 9.11E-01 | 0.01 | 0.13 | 0.99 |
| Adipose_Subcutaneous | ENSG00000273483.1 | -597059 | 442 | 214 | 9.12E-01 | 0.01 | 0.07 | 0.99 |
| Skin_Not_Sun_Exposed_Suprapubic | ENSG00000232811.1 | 977915 | 387 | 200 | 9.12E-01 | -0.01 | 0.07 | 0.99 |
| Esophagus_Gastroesophageal_Junction | ENSG00000273483.1 | -597059 | 244 | 110 | 9.12E-01 | 0.01 | 0.07 | 0.99 |
| Ovary | ENSG00000238975.1 | -731206 | 133 | 71 | 9.12E-01 | 0.01 | 0.12 | 0.99 |
| Colon_Transverse | ENSG00000260948.1 | 488314 | 274 | 119 | 9.13E-01 | -0.01 | 0.07 | 0.99 |
| Thyroid | ENSG00000233337.1 | 483868 | 446 | 224 | 9.14E-01 | -0.01 | 0.06 | 0.99 |
| Adipose_Subcutaneous | ENSG00000238975.1 | -731206 | 442 | 214 | 9.14E-01 | -0.01 | 0.07 | 0.99 |
| Breast_Mammary_Tissue | ENSG00000197852.8 | 168128 | 290 | 148 | 9.15E-01 | 0.00 | 0.04 | 0.99 |
| Esophagus_Mucosa | ENSG00000116459.6 | 472490 | 407 | 208 | 9.16E-01 | 0.00 | 0.02 | 0.99 |
| Esophagus_Mucosa | ENSG00000233337.1 | 483868 | 407 | 208 | 9.16E-01 | 0.01 | 0.08 | 0.99 |
| Whole_Blood | ENSG00000121933.13 | 357420 | 407 | 195 | 9.17E-01 | 0.00 | 0.04 | 0.99 |
| Skin_Sun_Exposed_Lower_leg | ENSG00000085465.11 | 493605 | 473 | 233 | 9.18E-01 | 0.00 | 0.02 | 0.99 |
| Skin_Not_Sun_Exposed_Suprapubic | ENSG00000064886.9 | 720611 | 387 | 200 | 9.19E-01 | 0.00 | 0.05 | 0.99 |
| Brain_Hypothalamus | ENSG00000155366.12 | -786052 | 121 | 61 | 9.19E-01 | -0.01 | 0.05 | 0.99 |
| Heart_Atrial_Appendage | ENSG00000231346.1 | 312659 | 297 | 147 | 9.21E-01 | -0.01 | 0.06 | 0.99 |
| Colon_Sigmoid | ENSG00000231437.3 | -68388 | 233 | 107 | 9.23E-01 | 0.01 | 0.06 | 0.99 |
| Minor_Salivary_Gland | ENSG00000121933.13 | 357420 | 97 | 49 | 9.23E-01 | -0.01 | 0.09 | 0.99 |
| Brain_Cerebellum | ENSG00000007341.14 | -699443 | 173 | 96 | 9.24E-01 | -0.01 | 0.09 | 0.99 |
| Adipose_Subcutaneous | ENSG00000143110.7 | 447590 | 442 | 214 | 9.24E-01 | 0.00 | 0.02 | 0.99 |
| Thyroid | ENSG00000225075.1 | -775239 | 446 | 224 | 9.24E-01 | -0.01 | 0.06 | 0.99 |
| Liver | ENSG00000225075.1 | -775239 | 175 | 88 | 9.25E-01 | -0.01 | 0.12 | 0.99 |
| Whole_Blood | ENSG00000238975.1 | -731206 | 407 | 195 | 9.25E-01 | 0.00 | 0.05 | 0.99 |
| Brain_Amygdala | ENSG00000143110.7 | 447590 | 100 | 47 | 9.28E-01 | 0.01 | 0.10 | 0.99 |
| Pancreas | ENSG00000116489.8 | -698415 | 248 | 108 | 9.28E-01 | 0.00 | 0.05 | 0.99 |
| Testis | ENSG00000227179.2 | 529563 | 259 | 128 | 9.28E-01 | 0.01 | 0.08 | 0.99 |
| Colon_Transverse | ENSG00000085465.11 | 493605 | 274 | 119 | 9.29E-01 | 0.00 | 0.05 | 0.99 |
| Thyroid | ENSG00000134255.9 | 781156 | 446 | 224 | 9.29E-01 | 0.00 | 0.03 | 0.99 |
| Brain_Hypothalamus | ENSG00000173947.9 | 575094 | 121 | 61 | 9.30E-01 | -0.01 | 0.07 | 0.99 |
| Spleen | ENSG00000233337.1 | 483868 | 162 | 55 | 9.31E-01 | 0.01 | 0.13 | 0.99 |
| Lung | ENSG00000155363.14 | -751759 | 427 | 222 | 9.31E-01 | 0.00 | 0.02 | 0.99 |

| | | | | | | | | |
|---|---|---|---|---|---|---|---|---|
| Artery_Coronary | ENSG00000225075.1 | -775239 | 173 | 81 | 9.31E-01 | 0.01 | 0.12 | 0.99 |
| Brain_Cortex | ENSG00000232811.1 | 977915 | 158 | 79 | 9.31E-01 | -0.01 | 0.13 | 0.99 |
| Artery_Coronary | ENSG00000273010.1 | 958080 | 173 | 81 | 9.31E-01 | 0.01 | 0.13 | 0.99 |
| Testis | ENSG00000116489.8 | -698415 | 259 | 128 | 9.32E-01 | 0.00 | 0.04 | 0.99 |
| Adipose_Visceral_Omentum | ENSG00000238975.1 | -731206 | 355 | 172 | 9.32E-01 | 0.01 | 0.07 | 0.99 |
| Thyroid | ENSG00000156171.10 | 781166 | 446 | 224 | 9.33E-01 | 0.00 | 0.04 | 0.99 |
| Brain_Caudate_basal_ganglia | ENSG00000231246.1 | -439146 | 160 | 88 | 9.33E-01 | -0.01 | 0.11 | 0.99 |
| Muscle_Skeletal | ENSG00000260948.1 | 488314 | 564 | 276 | 9.33E-01 | 0.00 | 0.04 | 0.99 |
| Lung | ENSG00000143110.7 | 447590 | 427 | 222 | 9.34E-01 | 0.00 | 0.03 | 0.99 |
| Esophagus_Mucosa | ENSG00000171385.5 | -67773 | 407 | 208 | 9.36E-01 | 0.00 | 0.04 | 0.99 |
| Liver | ENSG00000273483.1 | -597059 | 175 | 88 | 9.36E-01 | 0.01 | 0.08 | 0.99 |
| Spleen | ENSG00000215867.4 | 271838 | 162 | 55 | 9.40E-01 | -0.01 | 0.16 | 0.99 |
| Esophagus_Mucosa | ENSG00000273010.1 | 958080 | 407 | 208 | 9.40E-01 | 0.01 | 0.09 | 0.99 |
| Uterus | ENSG00000116489.8 | -698415 | 111 | 63 | 9.40E-01 | -0.01 | 0.09 | 0.99 |
| Adipose_Subcutaneous | ENSG00000231346.1 | 312659 | 442 | 214 | 9.41E-01 | 0.00 | 0.05 | 0.99 |
| Testis | ENSG00000143079.10 | -474799 | 259 | 128 | 9.41E-01 | 0.00 | 0.04 | 0.99 |
| Whole_Blood | ENSG00000235299.2 | -704842 | 407 | 195 | 9.41E-01 | 0.00 | 0.07 | 0.99 |
| Testis | ENSG00000231346.1 | 312659 | 259 | 128 | 9.41E-01 | 0.00 | 0.06 | 0.99 |
| Brain_Frontal_Cortex_BA9 | ENSG00000085465.11 | 493605 | 129 | 73 | 9.43E-01 | 0.00 | 0.06 | 0.99 |
| Ovary | ENSG00000184599.9 | -799037 | 133 | 71 | 9.44E-01 | -0.01 | 0.09 | 0.99 |
| Liver | ENSG00000231346.1 | 312659 | 175 | 88 | 9.44E-01 | 0.01 | 0.09 | 0.99 |
| Skin_Not_Sun_Exposed_Suprapubic | ENSG00000260948.1 | 488314 | 387 | 200 | 9.44E-01 | 0.00 | 0.05 | 0.99 |
| Adipose_Subcutaneous | ENSG00000064886.9 | 720611 | 442 | 214 | 9.45E-01 | 0.00 | 0.06 | 0.99 |
| Brain_Anterior_cingulate_cortex_BA24 | ENSG00000064886.9 | 720611 | 121 | 65 | 9.45E-01 | 0.01 | 0.10 | 0.99 |
| Brain_Cerebellum | ENSG00000231346.1 | 312659 | 173 | 96 | 9.46E-01 | -0.01 | 0.11 | 0.99 |
| Esophagus_Muscularis | ENSG00000162777.12 | 716847 | 370 | 188 | 9.47E-01 | 0.00 | 0.04 | 0.99 |
| Artery_Aorta | ENSG00000243960.1 | 482557 | 299 | 152 | 9.47E-01 | 0.01 | 0.09 | 0.99 |
| Thyroid | ENSG00000155366.12 | -786052 | 446 | 224 | 9.48E-01 | 0.00 | 0.03 | 0.99 |
| Ovary | ENSG00000155367.11 | -794095 | 133 | 71 | 9.48E-01 | 0.01 | 0.10 | 0.99 |
| Prostate | ENSG00000227179.2 | 529563 | 152 | 75 | 9.49E-01 | -0.01 | 0.10 | 0.99 |
| Skin_Not_Sun_Exposed_Suprapubic | ENSG00000116489.8 | -698415 | 387 | 200 | 9.51E-01 | 0.00 | 0.03 | 0.99 |
| Artery_Coronary | ENSG00000064703.7 | 166137 | 173 | 81 | 9.52E-01 | 0.00 | 0.05 | 0.99 |
| Brain_Nucleus_accumbens_basal_ganglia | ENSG00000162777.12 | 716847 | 147 | 77 | 9.52E-01 | 0.01 | 0.09 | 0.99 |
| Whole_Blood | ENSG00000197852.8 | 168128 | 407 | 195 | 9.52E-01 | 0.00 | 0.02 | 0.99 |
| Esophagus_Mucosa | ENSG00000155366.12 | -786052 | 407 | 208 | 9.54E-01 | 0.00 | 0.04 | 1.00 |
| Colon_Transverse | ENSG00000116473.10 | 379164 | 274 | 119 | 9.55E-01 | 0.00 | 0.03 | 1.00 |
| Esophagus_Gastroesophageal_Junction | ENSG00000197852.8 | 168128 | 244 | 110 | 9.56E-01 | 0.00 | 0.06 | 1.00 |
| Small_Intestine_Terminal_Ileum | ENSG00000231346.1 | 312659 | 137 | 54 | 9.56E-01 | 0.00 | 0.09 | 1.00 |
| Brain_Frontal_Cortex_BA9 | ENSG00000155363.14 | -751759 | 129 | 73 | 9.56E-01 | 0.00 | 0.06 | 1.00 |
| Artery_Tibial | ENSG00000233337.1 | 483868 | 441 | 219 | 9.57E-01 | 0.00 | 0.07 | 1.00 |
| Skin_Sun_Exposed_Lower_leg | ENSG00000121931.11 | 968421 | 473 | 233 | 9.57E-01 | 0.00 | 0.04 | 1.00 |
| Uterus | ENSG00000121931.11 | 968421 | 111 | 63 | 9.58E-01 | -0.01 | 0.11 | 1.00 |
| Colon_Transverse | ENSG00000121933.13 | 357420 | 274 | 119 | 9.59E-01 | 0.00 | 0.06 | 1.00 |
| Skin_Sun_Exposed_Lower_leg | ENSG00000116473.10 | 379164 | 473 | 233 | 9.60E-01 | 0.00 | 0.03 | 1.00 |
| Cells_EBV-transformed_lymphocytes | ENSG00000121933.13 | 357420 | 130 | 68 | 9.60E-01 | -0.01 | 0.13 | 1.00 |
| Heart_Atrial_Appendage | ENSG00000121931.11 | 968421 | 297 | 147 | 9.61E-01 | 0.00 | 0.05 | 1.00 |
| Liver | ENSG00000064703.7 | 166137 | 175 | 88 | 9.62E-01 | 0.00 | 0.08 | 1.00 |

| | | | | | | | | |
|---|---|---|---|---|---|---|---|---|
| Nerve_Tibial | ENSG00000231346.1 | 312659 | 414 | 202 | 9.62E-01 | 0.00 | 0.06 | 1.00 |
| Brain_Putamen_basal_ganglia | ENSG00000224167.1 | -955558 | 124 | 71 | 9.62E-01 | 0.01 | 0.15 | 1.00 |
| Brain_Caudate_basal_ganglia | ENSG00000155366.12 | -786052 | 160 | 88 | 9.63E-01 | 0.00 | 0.04 | 1.00 |
| Pancreas | ENSG00000155363.14 | -751759 | 248 | 108 | 9.64E-01 | 0.00 | 0.04 | 1.00 |
| Spleen | ENSG00000260948.1 | 488314 | 162 | 55 | 9.65E-01 | -0.01 | 0.12 | 1.00 |
| Breast_Mammary_Tissue | ENSG00000227179.2 | 529563 | 290 | 148 | 9.65E-01 | 0.00 | 0.07 | 1.00 |
| Brain_Cerebellar_Hemisphere | ENSG00000116455.9 | 472519 | 136 | 74 | 9.66E-01 | 0.00 | 0.09 | 1.00 |
| Colon_Sigmoid | ENSG00000064886.9 | 720611 | 233 | 107 | 9.67E-01 | 0.00 | 0.07 | 1.00 |
| Adipose_Visceral_Omentum | ENSG00000064703.7 | 166137 | 355 | 172 | 9.68E-01 | 0.00 | 0.04 | 1.00 |
| Testis | ENSG00000134216.14 | 630520 | 259 | 128 | 9.68E-01 | 0.00 | 0.09 | 1.00 |
| Pancreas | ENSG00000116473.10 | 379164 | 248 | 108 | 9.68E-01 | 0.00 | 0.05 | 1.00 |
| Artery_Tibial | ENSG00000116489.8 | -698415 | 441 | 219 | 9.68E-01 | 0.00 | 0.04 | 1.00 |
| Whole_Blood | ENSG00000116459.6 | 472490 | 407 | 195 | 9.68E-01 | 0.00 | 0.02 | 1.00 |
| Stomach | ENSG00000064703.7 | 166137 | 262 | 107 | 9.71E-01 | 0.00 | 0.05 | 1.00 |
| Uterus | ENSG00000085465.11 | 493605 | 111 | 63 | 9.72E-01 | 0.00 | 0.10 | 1.00 |
| Uterus | ENSG00000162777.12 | 716847 | 111 | 63 | 9.72E-01 | 0.00 | 0.10 | 1.00 |
| Adipose_Visceral_Omentum | ENSG00000121931.11 | 968421 | 355 | 172 | 9.73E-01 | 0.00 | 0.05 | 1.00 |
| Brain_Cerebellum | ENSG00000143110.7 | 447590 | 173 | 96 | 9.73E-01 | 0.00 | 0.07 | 1.00 |
| Adipose_Subcutaneous | ENSG00000203878.7 | 641323 | 442 | 214 | 9.74E-01 | 0.00 | 0.06 | 1.00 |
| Adipose_Visceral_Omentum | ENSG00000184599.9 | -799037 | 355 | 172 | 9.74E-01 | 0.00 | 0.06 | 1.00 |
| Ovary | ENSG00000143079.10 | -474799 | 133 | 71 | 9.76E-01 | 0.00 | 0.07 | 1.00 |
| Esophagus_Gastroesophageal_Junction | ENSG00000134255.9 | 781156 | 244 | 110 | 9.76E-01 | 0.00 | 0.05 | 1.00 |
| Brain_Cerebellum | ENSG00000134255.9 | 781156 | 173 | 96 | 9.76E-01 | 0.00 | 0.06 | 1.00 |
| Small_Intestine_Terminal_Ileum | ENSG00000085465.11 | 493605 | 137 | 54 | 9.77E-01 | 0.00 | 0.07 | 1.00 |
| Brain_Substantia_nigra | ENSG00000121931.11 | 968421 | 88 | 49 | 9.80E-01 | 0.00 | 0.12 | 1.00 |
| Breast_Mammary_Tissue | ENSG00000173947.9 | 575094 | 290 | 148 | 9.80E-01 | 0.00 | 0.04 | 1.00 |
| Nerve_Tibial | ENSG00000134245.13 | -545159 | 414 | 202 | 9.81E-01 | 0.00 | 0.04 | 1.00 |
| Muscle_Skeletal | ENSG00000121933.13 | 357420 | 564 | 276 | 9.81E-01 | 0.00 | 0.04 | 1.00 |
| Brain_Caudate_basal_ganglia | ENSG00000232811.1 | 977915 | 160 | 88 | 9.81E-01 | 0.00 | 0.12 | 1.00 |
| Colon_Transverse | ENSG00000232811.1 | 977915 | 274 | 119 | 9.83E-01 | 0.00 | 0.08 | 1.00 |
| Brain_Anterior_cingulate_cortex_BA24 | ENSG00000171385.5 | -67773 | 121 | 65 | 9.84E-01 | 0.00 | 0.06 | 1.00 |
| Adipose_Subcutaneous | ENSG00000134255.9 | 781156 | 442 | 214 | 9.85E-01 | 0.00 | 0.03 | 1.00 |
| Stomach | ENSG00000162777.12 | 716847 | 262 | 107 | 9.86E-01 | 0.00 | 0.04 | 1.00 |
| Brain_Anterior_cingulate_cortex_BA24 | ENSG00000064703.7 | 166137 | 121 | 65 | 9.86E-01 | 0.00 | 0.10 | 1.00 |
| Skin_Not_Sun_Exposed_Suprapubic | ENSG00000243960.1 | 482557 | 387 | 200 | 9.86E-01 | 0.00 | 0.07 | 1.00 |
| Liver | ENSG00000260948.1 | 488314 | 175 | 88 | 9.86E-01 | 0.00 | 0.11 | 1.00 |
| Esophagus_Muscularis | ENSG00000134245.13 | -545159 | 370 | 188 | 9.87E-01 | 0.00 | 0.06 | 1.00 |
| Testis | ENSG00000007341.14 | -699443 | 259 | 128 | 9.88E-01 | 0.00 | 0.05 | 1.00 |
| Brain_Cerebellar_Hemisphere | ENSG00000143110.7 | 447590 | 136 | 74 | 9.88E-01 | 0.00 | 0.06 | 1.00 |
| Pituitary | ENSG00000273010.1 | 958080 | 183 | 95 | 9.88E-01 | 0.00 | 0.12 | 1.00 |
| Esophagus_Gastroesophageal_Junction | ENSG00000243960.1 | 482557 | 244 | 110 | 9.89E-01 | 0.00 | 0.11 | 1.00 |
| Uterus | ENSG00000173947.9 | 575094 | 111 | 63 | 9.90E-01 | 0.00 | 0.09 | 1.00 |
| Minor_Salivary_Gland | ENSG00000197852.8 | 168128 | 97 | 49 | 9.90E-01 | 0.00 | 0.12 | 1.00 |
| Brain_Cortex | ENSG00000116489.8 | -698415 | 158 | 79 | 9.91E-01 | 0.00 | 0.06 | 1.00 |
| Nerve_Tibial | ENSG00000233337.1 | 483868 | 414 | 202 | 9.92E-01 | 0.00 | 0.06 | 1.00 |
| Artery_Tibial | ENSG00000162777.12 | 716847 | 441 | 219 | 9.92E-01 | 0.00 | 0.03 | 1.00 |
| Adipose_Subcutaneous | ENSG00000155366.12 | -786052 | 442 | 214 | 9.93E-01 | 0.00 | 0.03 | 1.00 |

| | | | | | | | | |
|---|---|---|---|---|---|---|---|---|
| Vagina | ENSG00000064703.7 | 166137 | 115 | 59 | 9.94E-01 | 0.00 | 0.08 | 1.00 |
| Heart_Left_Ventricle | ENSG00000116473.10 | 379164 | 303 | 154 | 9.95E-01 | 0.00 | 0.03 | 1.00 |
| Adipose_Subcutaneous | ENSG00000121933.13 | 357420 | 442 | 214 | 9.95E-01 | 0.00 | 0.03 | 1.00 |
| Small_Intestine_Terminal_Ileum | ENSG00000261654.1 | 979029 | 137 | 54 | 9.95E-01 | 0.00 | 0.10 | 1.00 |
| Skin_Sun_Exposed_Lower_leg | ENSG00000225075.1 | -775239 | 473 | 233 | 9.96E-01 | 0.00 | 0.07 | 1.00 |
| Brain_Nucleus_accumbens_basal_ganglia | ENSG00000260948.1 | 488314 | 147 | 77 | 9.97E-01 | 0.00 | 0.10 | 1.00 |
| Skin_Sun_Exposed_Lower_leg | ENSG00000184599.9 | -799037 | 473 | 233 | 9.97E-01 | 0.00 | 0.06 | 1.00 |
| Brain_Cortex | ENSG00000156171.10 | 781166 | 158 | 79 | 9.97E-01 | 0.00 | 0.10 | 1.00 |
| Prostate | ENSG00000171385.5 | -67773 | 152 | 75 | 9.98E-01 | 0.00 | 0.09 | 1.00 |
| Small_Intestine_Terminal_Ileum | ENSG00000215866.3 | -929261 | 137 | 54 | 9.99E-01 | 0.00 | 0.15 | 1.00 |
| Brain_Anterior_cingulate_cortex_BA24 | ENSG00000227811.2 | 181541 | 121 | 65 | 9.99E-01 | 0.00 | 0.12 | 1.00 |
| Small_Intestine_Terminal_Ileum | ENSG00000273483.1 | -597059 | 137 | 54 | 9.99E-01 | 0.00 | 0.12 | 1.00 |

tss_distance: distance in bp to transcription start site; slope_se: standard error of the estimated regression slope; MAC: minor allele count; FDR: false discovery rate

***Supplementary Table 5: Co-localization results for all tissues with significant cis eQTL associations of rs1545300***

| ***Heart Left Ventricle*** **transcript** | **Gene** | **b_SMR** | **se_SMR** | **p_SMR** | **p_HEIDI** | **nsnp_HEIDI** |
|---|---|---|---|---|---|---|
| ENSG00000232811.1 | *RP11-96K19.2* | 4.01 | 8.08 | 6.20E-01 | NA | NA |
| ENSG00000121931.11 | *LRIF1* | 13.53 | 44.82 | 7.63E-01 | NA | NA |
| ENSG00000273010.1 | *RP11-96K19.5* | -2.51 | 2.82 | 3.73E-01 | NA | NA |
| ENSG00000156171.10 | *DRAM2* | 2.66 | 2.17 | 2.20E-01 | NA | NA |
| ENSG00000134255.9 | *CEPT1* | 2.86 | 2.36 | 2.26E-01 | NA | NA |
| ENSG00000064886.9 | *CHI3L2* | -4.24 | 6.68 | 5.26E-01 | NA | NA |
| ENSG00000162777.12 | *DENND2D* | 3.76 | 3.93 | 3.39E-01 | NA | NA |
| ENSG00000173947.9 | *PIFO* | 9.21 | 18.14 | 6.12E-01 | NA | NA |
| ENSG00000085465.11 | *OVGP1* | 2.50 | 1.57 | 1.10E-01 | NA | NA |
| ENSG00000260948.1 | *RP11-552M11.8* | -2.21 | 1.99 | 2.65E-01 | NA | NA |
| ENSG00000233337.1 | *UBE2FP3* | 8.53 | 37.89 | 8.22E-01 | NA | NA |
| ENSG00000243960.1 | *RP11-552M11.4* | 8.27 | 35.09 | 8.14E-01 | NA | NA |
| ENSG00000116455.9 | *WDR77* | -50.98 | 390.71 | 8.96E-01 | NA | NA |
| ENSG00000116459.6 | *ATP5F1* | -58.83 | 429.76 | 8.91E-01 | NA | NA |
| ENSG00000143110.7 | *C1orf162* | 1.33 | 0.52 | 1.10E-02 | NA | NA |
| ENSG00000116473.10 | *RAP1A* | -1019.00 | 151743.00 | 9.95E-01 | NA | NA |
| ENSG00000121933.13 | *ADORA3* | 4.91 | 6.73 | 4.66E-01 | NA | NA |
| ENSG00000231346.1 | *RP5-836N10.1* | -10.55 | 33.21 | 7.51E-01 | NA | NA |
| ENSG00000227811.2 | *RP4-773A18.4* | -1.64 | 0.96 | 8.84E-02 | NA | NA |
| ENSG00000197852.8 | *FAM212B* | -31.29 | 212.48 | 8.83E-01 | NA | NA |
| ENSG00000064703.7 | *DDX20* | -8.29 | 16.20 | 6.09E-01 | NA | NA |
| **ENSG00000171385.5** | ***KCND3*** | **1.69** | **0.53** | **1.37E-03** | **1.87E-01** | **19** |
| ENSG00000231437.3 | *RP11-88H9.2* | -10.26 | 33.65 | 7.60E-01 | NA | 3 |
| ENSG00000231246.1 | *RP5-965F6.2* | 6.37 | 16.71 | 7.03E-01 | NA | NA |
| ENSG00000143079.10 | *CTTNBP2NL* | -8.98 | 16.40 | 5.84E-01 | NA | NA |
| ENSG00000134245.13 | *WNT2B* | 3.79 | 4.35 | 3.84E-01 | NA | NA |
| ENSG00000273483.1 | *RP4-671G15.2* | -7.66 | 18.89 | 6.85E-01 | NA | NA |
| ENSG00000116489.8 | *CAPZA1* | 7.45 | 8.68 | 3.91E-01 | NA | NA |
| ENSG00000007341.14 | *ST7L* | -19.04 | 135.17 | 8.88E-01 | NA | NA |
| ENSG00000155363.14 | *MOV10* | 10.16 | 17.21 | 5.55E-01 | NA | NA |
| ENSG00000225075.1 | *RP11-426L16.3* | -3.62 | 6.40 | 5.72E-01 | NA | NA |
| ENSG00000155366.12 | *RHOC* | 6.08 | 5.97 | 3.08E-01 | NA | NA |
| ENSG00000155367.11 | *PPM1J* | -4.03 | 4.17 | 3.34E-01 | NA | NA |

| *Artery Tibial* | | | | | | |
|---|---|---|---|---|---|---|
| **transcript** | **Gene** | **b_SMR** | **se_SMR** | **p_SMR** | **p_HEIDI** | **nsnp_HEIDI** |
| ENSG00000261654.1 | *RP11-96K19.4* | 2.44 | 2.03 | 2.30E-01 | NA | NA |
| ENSG00000232811.1 | *RP11-96K19.2* | 1.72 | 1.11 | 1.19E-01 | NA | NA |
| ENSG00000121931.11 | *LRIF1* | 5.32 | 5.56 | 3.39E-01 | NA | NA |
| ENSG00000156171.10 | *DRAM2* | 3.60 | 2.63 | 1.71E-01 | NA | NA |
| ENSG00000134255.9 | *CEPT1* | 5.08 | 4.92 | 3.02E-01 | NA | NA |
| ENSG00000064886.9 | *CHI3L2* | 4.08 | 4.51 | 3.65E-01 | NA | NA |
| ENSG00000162777.12 | *DENND2D* | 625.65 | 63124.30 | 9.92E-01 | NA | NA |
| ENSG00000173947.9 | *PIFO* | 6.96 | 13.02 | 5.93E-01 | NA | NA |
| ENSG00000085465.11 | *OVGP1* | -30.16 | 156.96 | 8.48E-01 | NA | NA |
| ENSG00000260948.1 | *RP11-552M11.8* | 3.00 | 2.40 | 2.11E-01 | NA | NA |
| ENSG00000233337.1 | *UBE2FP3* | 48.75 | 893.91 | 9.57E-01 | NA | NA |
| ENSG00000243960.1 | *RP11-552M11.4* | -5.26 | 10.68 | 6.22E-01 | NA | NA |
| ENSG00000116455.9 | *WDR77* | -15.72 | 37.74 | 6.77E-01 | NA | NA |
| ENSG00000116459.6 | *ATP5F1* | -63.05 | 493.02 | 8.98E-01 | NA | NA |
| ENSG00000143110.7 | *C1orf162* | 12.96 | 28.64 | 6.51E-01 | NA | NA |
| ENSG00000116473.10 | *RAP1A* | 7.10 | 6.51 | 2.76E-01 | NA | NA |
| ENSG00000121933.13 | *ADORA3* | 32.65 | 232.74 | 8.88E-01 | NA | NA |
| ENSG00000231346.1 | *RP5-836N10.1* | -4.12 | 3.08 | 1.81E-01 | NA | NA |
| ENSG00000227811.2 | *RP4-773A18.4* | -1.90 | 1.40 | 1.74E-01 | NA | NA |
| ENSG00000197852.8 | *FAM212B* | 5.15 | 5.05 | 3.08E-01 | NA | NA |
| ENSG00000064703.7 | *DDX20* | -5.71 | 5.05 | 2.58E-01 | NA | NA |
| **ENSG00000171385.5** | ***KCND3*** | **1.02** | **0.26** | **1.15E-04** | **4.41E-02** | **12** |
| ENSG00000231437.3 | *RP11-88H9.2* | 7.84 | 16.99 | 6.44E-01 | NA | NA |
| ENSG00000231246.1 | *RP5-965F6.2* | 9.26 | 28.04 | 7.41E-01 | NA | NA |
| ENSG00000143079.10 | *CTTNBP2NL* | 5.90 | 6.23 | 3.43E-01 | NA | NA |
| ENSG00000134245.13 | *WNT2B* | -17.94 | 67.70 | 7.91E-01 | NA | NA |
| ENSG00000273483.1 | *RP4-671G15.2* | 3.58 | 4.36 | 4.11E-01 | NA | NA |
| ENSG00000116489.8 | *CAPZA1* | 140.56 | 3549.05 | 9.68E-01 | NA | NA |
| ENSG00000007341.14 | *ST7L* | 1.67 | 0.87 | 5.50E-02 | NA | NA |
| ENSG00000155363.14 | *MOV10* | 4.60 | 3.59 | 2.01E-01 | NA | NA |
| ENSG00000155366.12 | *RHOC* | 6.92 | 5.57 | 2.14E-01 | NA | NA |
| ENSG00000155367.11 | *PPM1J* | -3.18 | 2.17 | 1.43E-01 | NA | NA |

| *Minor Salivary Gland* transcript | Gene | b_SMR | se_SMR | p_SMR | p_HEIDI | nsnp_HEIDI |
|---|---|---|---|---|---|---|
| ENSG00000261654.1 | *RP11-96K19.4* | -0.60 | 0.28 | 3.31E-02 | NA | NA |
| ENSG00000232811.1 | *RP11-96K19.2* | -0.58 | 0.27 | 3.06E-02 | NA | NA |
| ENSG00000121931.11 | *LRIF1* | -1.28 | 0.94 | 1.76E-01 | NA | NA |
| ENSG00000273010.1 | *RP11-96K19.5* | 1.67 | 1.99 | 4.03E-01 | NA | NA |
| ENSG00000156171.10 | *DRAM2* | 0.73 | 0.33 | 2.48E-02 | 2.69E-02 | 3 |
| **ENSG00000134255.9** | ***CEPT1*** | **0.57** | **0.18** | **1.51E-03** | **1.70E-03** | **12** |
| ENSG00000064886.9 | *CHI3L2* | 1.73 | 1.11 | 1.18E-01 | NA | NA |
| ENSG00000162777.12 | *DENND2D* | 1.85 | 1.57 | 2.38E-01 | 5.91E-02 | 6 |
| ENSG00000173947.9 | *PIFO* | -0.87 | 0.36 | 1.44E-02 | NA | NA |
| ENSG00000085465.11 | *OVGP1* | -1.70 | 1.34 | 2.04E-01 | NA | NA |
| ENSG00000260948.1 | *RP11-552M11.8* | -0.80 | 0.37 | 2.87E-02 | NA | NA |
| ENSG00000233337.1 | *UBE2FP3* | -1.53 | 1.13 | 1.74E-01 | NA | NA |
| ENSG00000243960.1 | *RP11-552M11.4* | -2.03 | 2.40 | 3.97E-01 | NA | NA |
| ENSG00000116455.9 | *WDR77* | 6.57 | 18.71 | 7.26E-01 | NA | NA |
| ENSG00000116459.6 | *ATP5F1* | -6.38 | 13.12 | 6.27E-01 | NA | NA |
| ENSG00000143110.7 | *C1orf162* | -2.04 | 1.92 | 2.87E-01 | NA | NA |
| ENSG00000116473.10 | *RAP1A* | 5.59 | 16.79 | 7.39E-01 | NA | NA |
| ENSG00000121933.13 | *ADORA3* | 23.76 | 245.92 | 9.23E-01 | NA | NA |
| ENSG00000231346.1 | *RP5-836N10.1* | 0.77 | 0.39 | 4.74E-02 | NA | NA |
| ENSG00000197852.8 | *FAM212B* | 132.35 | 10650.60 | 9.90E-01 | NA | NA |
| ENSG00000064703.7 | *DDX20* | -3.78 | 8.65 | 6.62E-01 | NA | NA |
| ENSG00000171385.5 | *KCND3* | 1.51 | 1.14 | 1.83E-01 | NA | NA |
| ENSG00000231437.3 | *RP11-88H9.2* | 1.72 | 1.87 | 3.58E-01 | NA | NA |
| ENSG00000231246.1 | *RP5-965F6.2* | 11.43 | 82.98 | 8.90E-01 | NA | NA |
| ENSG00000143079.10 | *CTTNBP2NL* | 5.68 | 8.57 | 5.07E-01 | NA | NA |
| ENSG00000134245.13 | *WNT2B* | 1.97 | 2.02 | 3.29E-01 | NA | NA |
| ENSG00000273483.1 | *RP4-671G15.2* | -0.89 | 0.57 | 1.21E-01 | NA | NA |
| ENSG00000116489.8 | *CAPZA1* | 4.24 | 5.45 | 4.36E-01 | NA | NA |
| ENSG00000007341.14 | *ST7L* | 0.71 | 0.42 | 9.49E-02 | NA | NA |
| ENSG00000238975.1 | *snoU13* | 2.05 | 3.03 | 4.97E-01 | NA | NA |
| ENSG00000155363.14 | *MOV10* | 1.97 | 1.70 | 2.46E-01 | NA | NA |
| ENSG00000225075.1 | *RP11-426L16.3* | 6.87 | 35.70 | 8.47E-01 | NA | NA |
| ENSG00000155366.12 | *RHOC* | -3.20 | 3.09 | 3.01E-01 | NA | NA |
| ENSG00000155367.11 | *PPM1J* | 3.40 | 6.75 | 6.14E-01 | NA | NA |
| ENSG00000184599.9 | *FAM19A3* | -1.49 | 1.60 | 3.53E-01 | NA | NA |

b_SMR, se_SMR, p_SMR: effect, standard error, p-value of the SMR test; p_HEIDI, nsnp_HEIDI: p-value, number of SNPs of the HEIDI test for heterogeneity. Co-localizations with p_SMR <0.01 are shown in bold.

### Supplementary Figures

#### *Supplementary Figure 1: Manhattan plot of the combined GWAS meta-analysis*

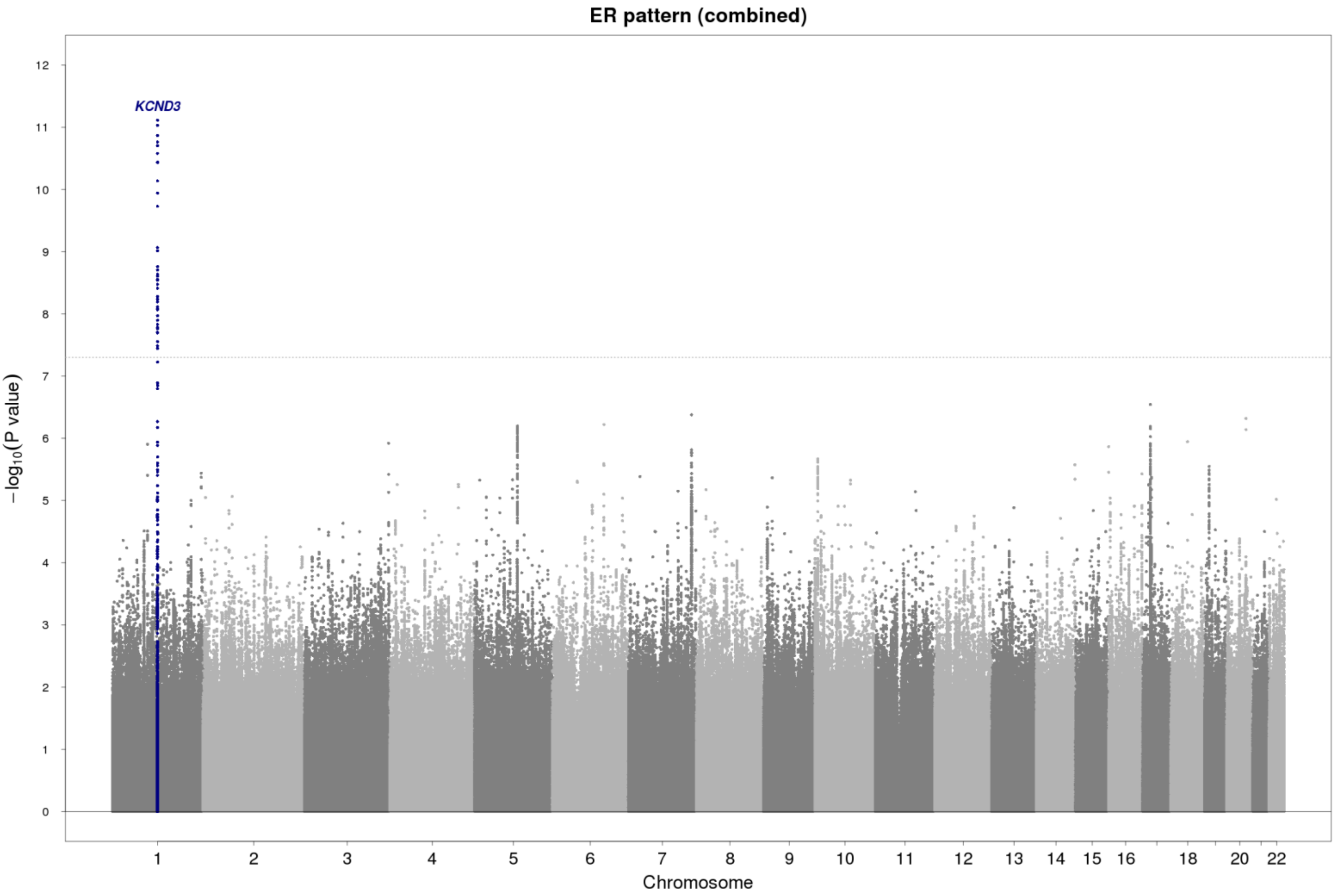

SNPs are plotted on the x-axis according to their position on each chromosome with the -$\log_{10}$(p-value) of the association test on the y-axis. The solid horizontal line indicates the threshold for genome-wide significance, $5 \times 10^{-8}$. The labels show the closest gene.

*Supplementary Figure 2: QQ plot of the combined GWAS meta-analysis*

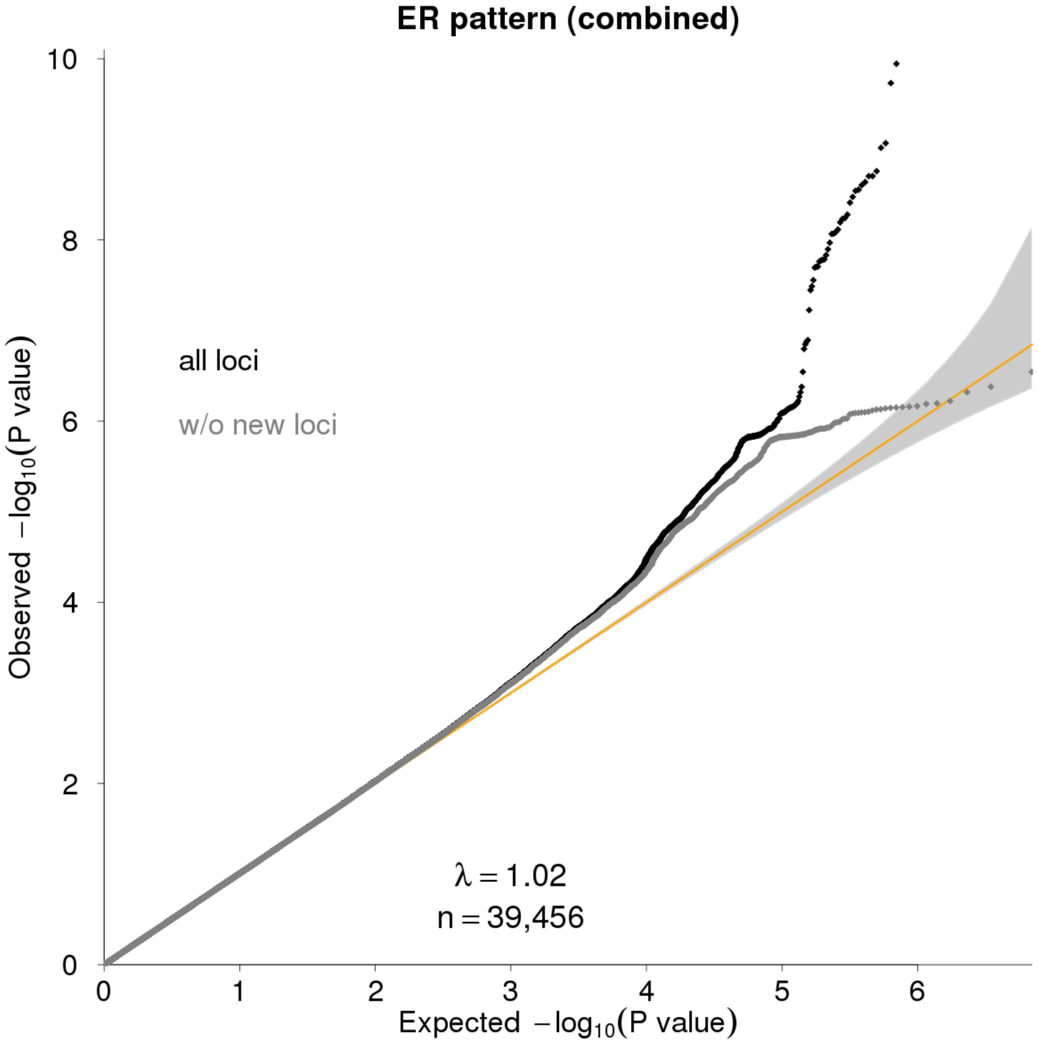

The observed p-values of the association test are plotted on the y-axis against their expected distribution under the null hypothesis of no association on the x-axis. Results for all SNPs are shown in black, and results after removal of loci (±500kb of the lead SNP) genome-wide significantly ($p<5x10^{-8}$) associated with the trait are shown in gray. Gray bands represent 95% confidence intervals. λ: lambda, genomic control parameter; n: sample size.

***Supplementary Figure 3: Regional association plot of rs17029069***

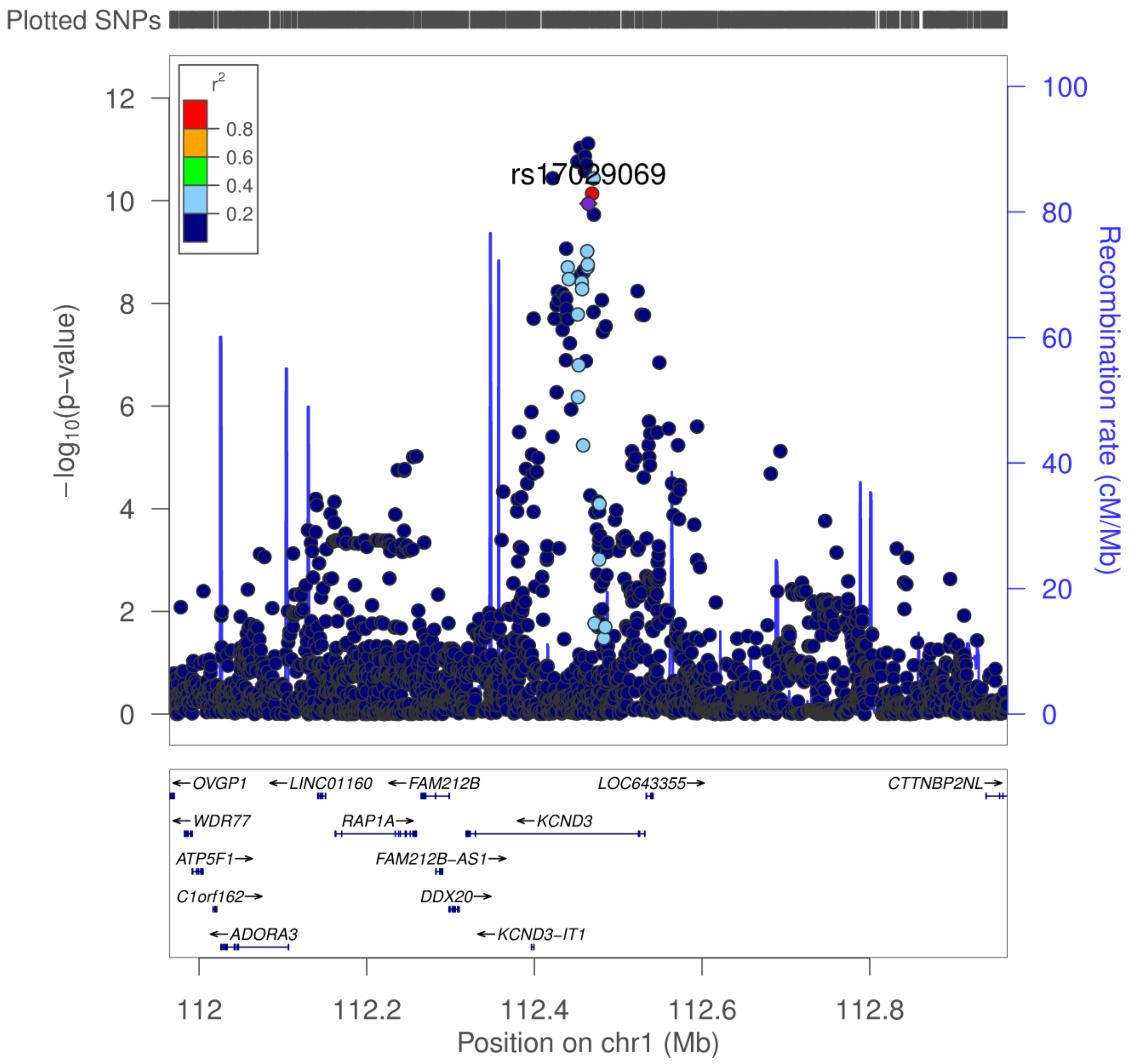

Regional association plots are shown for a reported candidate SNP rs17029069 of a previous GWAS. Correlation with the SNP (purple) is estimated based on the 1000 Genomes EUR reference samples. Plots were generated using the website of LocusZoom (Pruim, R. J. *et al.* Bioinformatics*,* 2010). Genetic positions refer to GRCh37/hg19 coordinates.
